# Endogenous SMN Heterogeneity Defines Dynamic States of Motor Neuron Vulnerability and Resilience

**DOI:** 10.64898/2026.09.23.753804

**Authors:** Joshua Thomas, Antonia Ramme, Jungyen Jun, Antonio Caldarelli, Sofia Vrettou, Fabian Rost, Zeynep Dokuzluoglu, Tobias Grass, Brunhilde Wirth, Natalia Rodriguez-Muela

## Abstract

Selective neuronal vulnerability, whereby some neurons degenerate while others remain resilient despite a shared genetic and disease context, is a defining feature of neurodegenerative diseases, yet the intrinsic mechanisms underlying these divergent fates remain poorly understood. Here, we identify naturally occurring heterogeneity in endogenous Survival Motor Neuron (SMN) protein abundance as a determinant of dynamic motor neuron (MN) vulnerability states that predict neuronal fate. Longitudinal live-cell imaging of human SMN-Clover reporter MNs shows that neurons with high endogenous SMN survive significantly longer than neighboring low-SMN neurons across healthy, spinal muscular atrophy (SMA) and amyotrophic lateral sclerosis (ALS) genetic backgrounds. These vulnerability states are not fixed: a substantial proportion of MNs progressively increase endogenous SMN abundance, transitioning toward a more resilient state associated with enhanced survival. Low-SMN neurons also exhibit increased spontaneous activity, linking endogenous SMN abundance to neuronal physiology and survival. In vivo, selectively vulnerable MN populations display corresponding shifts in endogenous SMN abundance during disease progression, supporting the relevance of SMN-defined vulnerability states beyond in vitro models. Integrated transcriptomic and proteomic analyses reveal distinct molecular programs associated with resilient and vulnerable states, including RNA metabolism, proteostasis, cytoskeletal organization, neuronal activity and stress responses. Among conserved molecular differences, the anti-apoptotic regulator LMO3 emerged as a candidate mediator of resilience. LMO3 overexpression reduced expression of p53-responsive genes, increased MN survival and shifted neurons toward higher endogenous SMN states. Together, these findings establish endogenous protein heterogeneity as a determinant of dynamic neuronal vulnerability and reveal neuronal resilience as a plastic molecular state that can be identified and therapeutically reinforced.

## Introduction

Selective neuronal vulnerability is a defining feature of virtually all neurodegenerative diseases, yet its biological basis remains one of the central unresolved questions in neuroscience. Within the same nervous system, genetically identical neuronal populations exposed to the same pathogenic mutation often exhibit remarkably different susceptibilities to degeneration, with some neurons dying early while others remain functional throughout disease progression. Understanding why certain neurons are intrinsically vulnerable whereas others are naturally resistant is therefore essential for identifying mechanisms that preserve neuronal function and for developing therapies that enhance endogenous resilience rather than merely slowing degeneration ^1–3^.

Motor neuron diseases (MNDs) provide an excellent paradigm to investigate selective neuronal vulnerability. Although Amyotrophic Lateral Sclerosis (ALS) and Spinal Muscular Atrophy (SMA) differ substantially in age of onset, genetics and clinical presentation, both disorders are characterized by the progressive degeneration of specific subsets of upper and lower motor neurons (MNs), ultimately leading to paralysis and premature death from respiratory failure ^4–6^. Importantly, degeneration does not occur uniformly across the motor system. Distinct spinal MN populations reproducibly display differential vulnerability, whereas ocular MNs and selected brainstem nuclei remain remarkably resistant in both diseases ^2,7–11^. Elegant anatomical studies have further demonstrated that even neighboring lumbar MN pools exhibit reproducible differences in vulnerability, highlighting that selective degeneration reflects intrinsic neuronal properties rather than simple differences in genetic background or disease exposure ^9,12,13^.

The remarkable coexistence of vulnerable and resistant MNs has motivated considerable efforts to identify endogenous protective mechanisms. Comparative analyses between resistant and vulnerable neuronal populations have uncovered several candidate resilience factors, including IGF-1/2 and GDF15, whose increased expression enhances MN survival across multiple disease models ^14,15^. These studies demonstrate that naturally resistant neurons can reveal therapeutically relevant survival programs. However, comparisons between anatomically distinct neuronal populations are inherently confounded by developmental origin, connectivity and physiological specialization, making it difficult to distinguish mechanisms that specifically regulate neuronal resilience from those reflecting normal subtype identity. Whether selective vulnerability can instead be studied within a genetically and developmentally homogeneous MN population remains largely unexplored.

One molecule of particular interest is the Survival of Motor Neuron (SMN) protein. SMN is a ubiquitously expressed multifunctional protein that participates in diverse cellular processes including snRNP assembly, RNA processing, mRNA transport, protein translation, cytoskeletal organization, bioenergetics and proteostasis ^16–23^. Homozygous loss of *SMN1* causes SMA, whereas variable *SMN2* copy number determines disease severity by modulating the amount of full-length SMN protein produced ^24–27^. The central importance of SMN is underscored by the remarkable clinical success of therapies that restore SMN expression ^28–30^. Beyond SMA, increasing evidence suggests that SMN also modulates MN vulnerability in ALS, where interactions with disease-associated RNA-binding proteins such as TDP-43 and FUS, together with genetic and experimental studies, support a broader neuroprotective role despite the absence of SMN deficiency as the primary genetic lesion.

Although SMN has traditionally been studied as a disease-associated protein, we previously discovered that endogenous SMN abundance varies substantially among genetically identical human MNs, generating naturally occurring populations expressing either relatively high or low SMN levels ^38^. Notably, MNs with higher endogenous SMN were significantly more resistant to toxic insults despite sharing the same genetic background, suggesting that endogenous protein heterogeneity itself may influence neuronal resilience. Such cell-to-cell variation is increasingly recognized as a fundamental feature of biological systems rather than experimental noise. Naturally occurring differences in gene or protein abundance can shape cell fate, stress responses and disease progression across diverse biological contexts, including cancer, immunology and neurodegeneration ^44–50^. We therefore hypothesized that endogenous SMN heterogeneity may not simply correlate with selective vulnerability but instead define dynamic cellular states that regulate MN resilience.

Here, we combine longitudinal single-cell live imaging, quantitative functional analyses, in vivo validation and integrated transcriptomic and proteomic profiling to determine how endogenous SMN abundance influences MN fate. We demonstrate that naturally occurring differences in endogenous SMN define dynamic vulnerability states that predict survival, neuronal excitability and disease progression across healthy, SMA and ALS genetic backgrounds. Furthermore, by comparing resilient and vulnerable MNs within genetically identical cultures, we identify LMO3 as a candidate regulator of endogenous resilience and demonstrate that enhancing this pathway promotes a resilient MN program. More broadly, our work establishes endogenous protein heterogeneity as a powerful strategy to uncover intrinsic mechanisms of selective neuronal vulnerability and identify therapeutic targets that enhance neuronal resilience across NDs.

## Results

### Endogenous SMN abundance predicts selective motor neuron survival

Although SMN deficiency causes selective motor neuron degeneration in SMA, individual motor neurons within the same culture express markedly different amounts of endogenous SMN protein ^38^. Whether this intrinsic heterogeneity merely correlates with, or directly determines, neuronal survival has remained unknown because endogenous SMN abundance could not previously be monitored in living cells. To address this, we generated healthy hiPSC lines carrying a Clover tag at one endogenous *SMN1* allele ^51^, allowing longitudinal quantification of SMN abundance in individual MNs. Two independent SMN-Clover knock-in clones were generated in each of two healthy hiPSC backgrounds (BJ and 1016A) and differentiated into spinal motor neurons using an established protocol ^19,51^ (Figure S1A-B). Reporter lines retained normal differentiation efficiency, survival, and the progressive increase in SMN abundance previously described for unedited neurons (Figure S1C-F). SMN protein levels were measured using the endogenous SMN-Clover reporter fluorescence and via antibody staining (recognizing both Clover-tagged and unedited SMN). On the MN population level, the average abundance of total SMN and SMN-Clover increased ∼4 times in the MNs derived from all lines after 10 days in culture, matching what we previously reported in unedited hiPSC lines ^38^ (Figure S1E-F). Single-cell quantification revealed a broad, continuous distribution of endogenous SMN expression spanning approximately five-fold across genetically identical motor neurons (Figure 1B-C). As cultures matured, the distribution progressively shifted toward higher SMN levels, suggesting that neurons surviving prolonged culture either already expressed higher SMN or progressively increased endogenous SMN abundance (Figure 1B-C, S1G-J). To directly determine whether endogenous SMN predicts survival, individual motor neurons were continuously tracked for four days, beginning at DIV4, while simultaneously quantifying SMN-Clover fluorescence (Figure 1D-E). Cells were objectively classified as high-, mid-, or low-SMN expressors using a standardized single-cell analysis pipeline before survival analysis was performed (Figure 1D). High-SMN motor neurons consistently exhibited a markedly greater probability of survival than mid- or low-expressing neurons across both independent healthy backgrounds (Figure 1F-G, S1K-L). Thus, endogenous differences in SMN abundance identify distinct pre-degenerative motor neuron states with profoundly different intrinsic survival probabilities. SMA is not classically considered to affect cortical neurons, but some evidence from living and post-mortem patient brains and SMA mouse models has revealed alterations in brain anatomy (e.g. multi-region atrophy and SMN-dependent signaling alterations) and even degeneration (e.g. motor cortex, thalamus, and cerebellum) ^52–56^. Therefore, to determine whether SMN protein heterogeneity has a MN specific impact, we performed a similar live imaging study on hiPSC-derived cortical neurons (CNs). WT BJ and 1016A SMN-Clover CNs were generated using doxycycline induced NGN2 overexpression as previously described ^38,57^ (Figure S1M). This protocol produced cultures with 75-80% BRN2^+^ CNs that degenerated overtime, increased SMN and SMN-Clover levels after 4 days in culture and displayed an analogous SMN and SMN-Clover heterogeneity as the MN cultures (Figure S1N-Q, 1H-J, S1R, respectively). Live, longitudinal imaging and analysis of single CNs revealed that SMN-Clover levels do not consistently determine the probability of survival of individual CNs (Figure 1K-M). Despite exhibiting endogenous SMN heterogeneity comparable to MNs, cortical neurons showed no consistent relationship between SMN abundance and survival (Figure 1K-M). Therefore, endogenous SMN abundance predicts neuronal vulnerability selectively in motor neurons rather than representing a universal feature of all neuronal populations.

**Figure 1.**
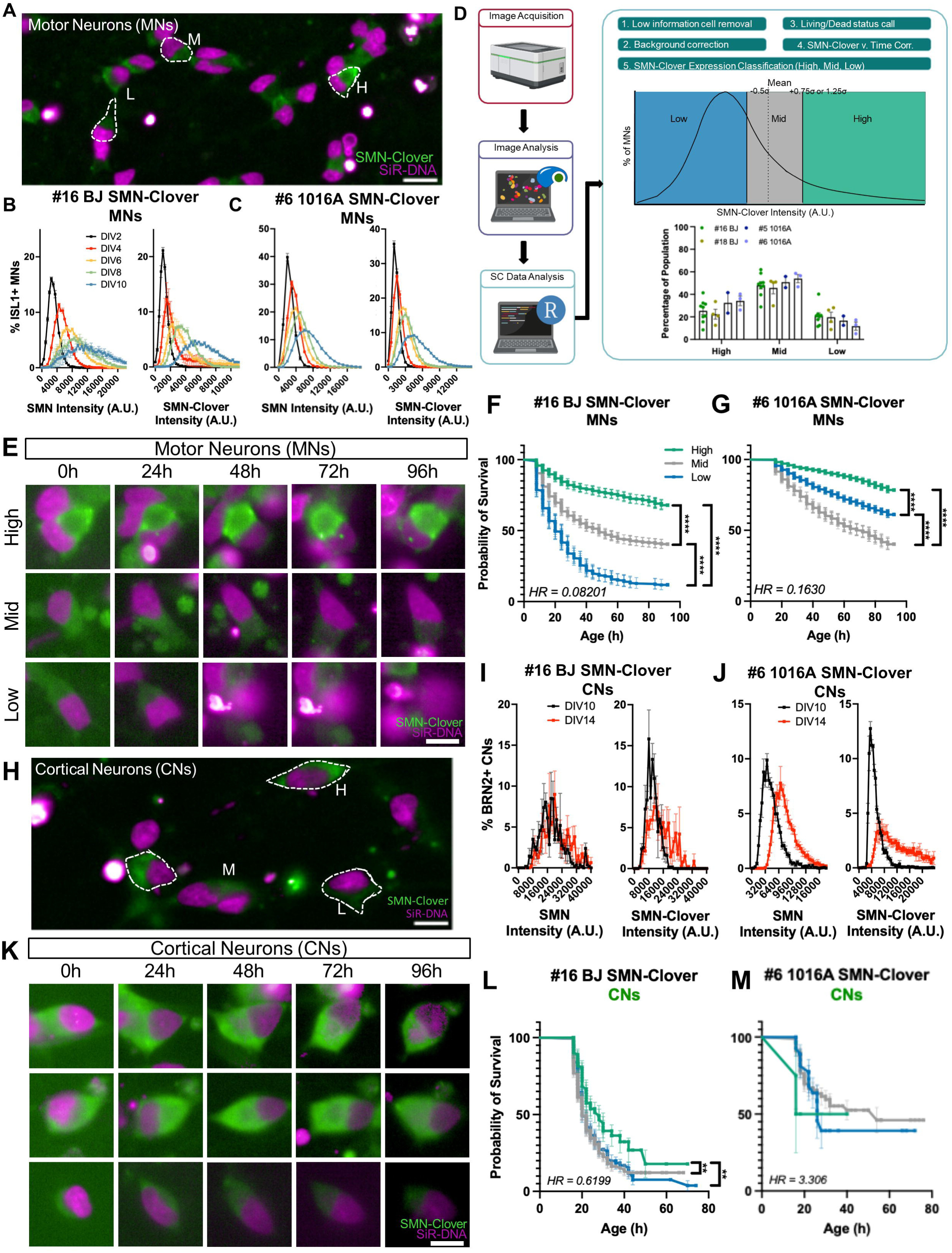
Endogenous SMN abundance predicts selective motor neuron survival A. Representative image of #16 BJ SMN-Clover MNs showing the heterogeneity of SMN-Clover expression. Representative MNs are outlined and classified as high (H)-, mid (M)-, or low (L)-SMN expressors at the indicated timepoints during a timelapse imaging experiment. SMN-Clover represented in green, SiR-DNA nuclear dye shown in magenta. Scale Bar = 20µm. B. Representative SMN (left) and SMN-Clover (right) Intensity (A.U.) histograms of #16 BJ SMN-Clover, hiSPC-derived ISL1+ MNs at timepoints from DIV2 to DIV10. C. Same as (B) for #6 1016A SMN-Clover. D. Schematic of protocol to generate hiPSC-derived MNs using small molecule patterning. Representative images of hiPSCs, embryoid bodies, and dissociated, plated hiPSC-derived MNs. E. Representative images of #16 BJ SMN-Clover MNs classified as high-, mid-, or low-SMN expressors at the indicated timepoints during a timelapse imaging experiment. SMN-Clover represented in green, SiR-DNA nuclear dye shown in magenta. Scale Bar = 20µm. F. Representative Kaplan-Meyer survival plot of #16 BJ SMN-Clover MNs plotting the probability of survival of MNs classified by their SMN-Clover expression level at each timepoint of the experiment (N=7). HR: hazard ratio between low- and high-expressors. Error bars: 95% confidence interval. Comparisons between survival curves was done using the Log-rank (Mantel-Cox) test. Significance represented as: 0.1234 (ns), 0.0332 (*), 0.0021 (**), 0.0002 (***), <0.0001 (****). G. Kaplan-Meyer survival plot of #6 1016A SMN-Clover MNs (N=3). See (F). H. Representative image of #16 BJ SMN-Clover MNs showing the heterogeneity of SMN-Clover expression. Representative MNs are outlined and as classified as high (H)-, mid (M)-, or low (L)-SMN expressors at the indicated timepoints during a timelapse imaging experiment. SMN-Clover represented in green, SiR-DNA nuclear dye shown in magenta. Scale Bar = 20µm. I. Representative SMN (left) and SMN-Clover (right) Intensity (A.U.) histogram of #16 BJ SMN-Clover, hiSPC-derived BRN2+ CNs at DIV10 and DIV14. J. Same as (I) for #6 1016A SMN-Clover. K. Representative images of #16 BJ SMN-Clover CNs classified as high-, mid-, or low-SMN expressors at the indicated timepoints during a timelapse imaging experiment. SMN-Clover represented in green, SiR-DNA nuclear dye shown in magenta. Scale Bar = 20µm. L. Kaplan-Meyer survival plot of #16 BJ SMN-Clover CNs (N=3). See (F). M. Kaplan-Meyer survival plot of #6 1016A SMN-Clover CNs (N=3). See (F).

We next asked whether endogenous SMN-defined vulnerability states simply reflected differential protein degradation. As we and others have previously stablished that SMN protein turnover is regulated by the ubiquitin-proteasome system (UPS) ^58,59^ and selective autophagy ^19,38^, we quantified the E3 ubiquitin ligase CULLIN5 and the selective autophagy receptor p62 across high-, mid-, and low-SMN MNs as proxies to determine UPS-mediated SMN degradation ^38^and selective autophagy-mediated SMN degradation. Neither marker differed significantly between SMN populations (Figure S2), arguing that intrinsic differences in protein degradation are unlikely to account for endogenous SMN-defined vulnerability states. This indicates that in WT MNs, SMN protein heterogeneity is not determined by cell intrinsic differences in SMN protein degradation rates. Endogenous SMN abundance therefore identifies distinct intrinsic motor neuron vulnerability states before degeneration begins.

### Endogenous SMN predicts motor neuron survival across SMA and ALS

Having established that endogenous SMN abundance determines survival in healthy motor neurons, we next asked whether this principle extends across genetically distinct motor neuron diseases (MNDs). As fusing the Clover reporter to the endogenous *SMN1* locus is not possible in our SMA hiPSC lines since no *SMN1* loci were detected ^38^, we fixed hiPSC-derived MNs at multiple time-points to determine if intrinsic heterogeneous SMN protein level determined SMA ISL1^+^ MN survival (Figure S3A-B). As expected, Type II (51N) and Type I (38D) and (33A) ISL1^+^ MNs showed significant degeneration and a marked reduction in SMN compared to WT BJ (Figure 2A, S3C). We observed a marked shift in the average and distribution of SMN protein expression as MNs were kept in culture, which correlated with a significant reduction in their survival (Figure 2B-C, S3D), matching our previous study ^38^. Despite the overall reduction in SMN abundance, the surviving SMA motor neurons became progressively enriched for higher SMN expressors, consistent with endogenous SMN continuing to predict neuronal resilience even in a disease context. Since high-SMN expressing MNs were consistently shown to be selectively resistant to degeneration, we next assessed whether there was an SMN threshold for survival. To address this, we classified DIV2 healthy WT and SMA MNs as high-, mid- and low-expressors and compared their population distribution according to their SMN expression levels (see *Star Methods*). As expected, the majority of SMA MNs were classified as mid- and low-expressors compared to healthy MNs due to their global SMN reduction (Figure 2D). Furthermore, low-SMN expressing healthy-WT MNs expressed only 40% the amount of SMN present in the high-SMN expressing healthy-WT MNs. Such levels of SMN in healthy-WT low expressors-MNs matched the mid-SMN expressing Type II and Type I SMA-MNs indicating that these MN subpopulations may be similarly vulnerable to degeneration. The low-SMN expressing SMA MNs expressed 20-30% of the SMN levels detected in the healthy high-SMN WT neurons, which aligns with the previously reported 15% SMN threshold at which SMA symptoms appear, as determined in the SMA *Smn^2B/-^,*mouse model ^60^ (Figure 2D).

**Figure 2.**
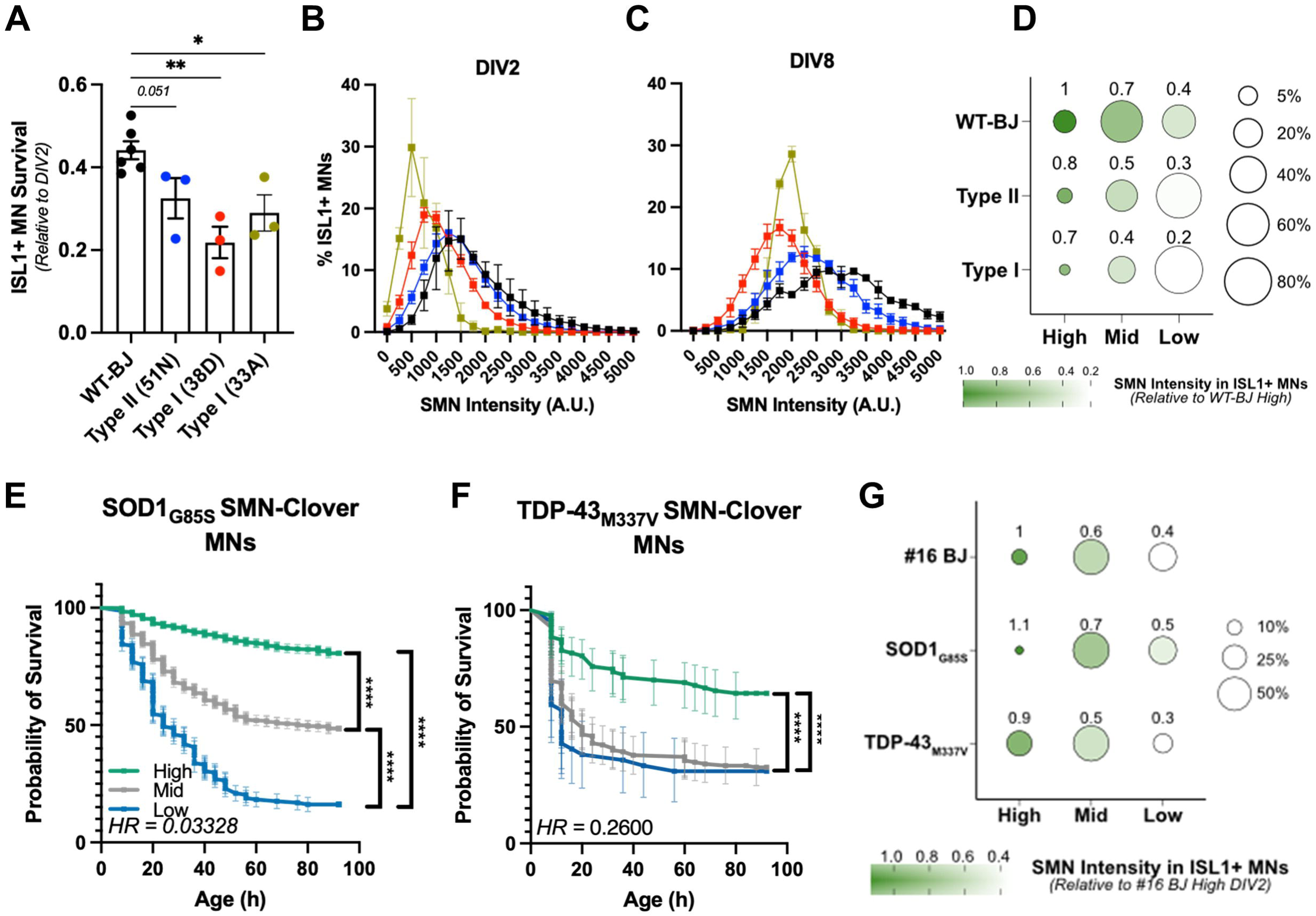
Endogenous SMN predicts motor neuron survival across ALS and SMA A. Fraction of ISL1+ WT-BJ, Type II (51N), Type I (38D), and Type I (33A) MNs that survived until DIV8 relative to the number quantified at DIV2. Statistical significance was calculated using an unpaired, nonparametric Kruskall-Wallis test with Uncorrected Dunn’s post hoc test. WT-BJ N=6; Type II (51N) N=3; Type I (38D) N=3; Type I (33A) N=3. B. Representative SMN Intensity (A.U.) histogram of WT (WT-BJ) and SMA (Type II-51N, Type I-38D, Type I-33A) hiSPC-derived ISL1+ MNs demonstrating both heterogeneous expression and increase in SMN abundance at DIV2. C. Representative SMN Intensity (A.U.) histogram of WT (WT-BJ) and SMA (Type II-51N, Type I-38D, Type I-33A) hiSPC-derived ISL1+ MNs at DIV8. D. Dot plot comparing SMN Intensity (color) and percentage of classified (size) from WT-BJ and SMA ISL1+ MNs classified as high-, mid-, and low-SMN expressors based on WT-BJ SMN expression at DIV2. E. Kaplan-Meyer survival plot of *SOD1*-G85S SMN-Clover MNs plotting the probability of survival of MNs classified by their SMN-Clover expression level at each timepoint of the experiment (N=3). HR: hazard ratio between low- and high-expressors. Error bars: 95% confidence interval. Comparisons between survival curves was done using the Log-rank (Mantel-Cox) test. Significance represented as: 0.1234 (ns), 0.0332 (*), 0.0021 (**), 0.0002 (***), <0.0001 (****). F. Representative Kaplan-Meyer survival plot of TDP-43_M337V_ SMN-Clover MNs (N=3). See (E). G. Dot plot comparing SMN Intensity (color) and percentage of classified (size) from #16 BJ and ALS ISL1+ MNs classified as high-, mid-, and low-SMN expressors based on #16 BJ SMN expression at DIV2.

To directly assess the impact of SMN protein levels on MN survival in ALS, two familial ALS (fALS)-causing mutations, SOD1_G85S_ and TDP-43_M337V_ ^61–63^ were introduced into the BJ SMN-Clover hiPSC line using CRISPR/Cas9-mediated genome editing (Figure S3E). Subjecting them to the same differentiation protocol, fALS SMN-Clover hiPSC lines produced 40-50% ISL1^+^ MNs and exhibited SMN protein expression levels and heterogeneity profiles analogous to the isogenic healthy controls (WTBJ and #16 BJ) (Figure S3F-K). To investigate whether the introduced fALS mutations led to a reported disease phenotype, healthy and SMN-Clover fALS MNs we exposed to oxidative stress, an established degeneration trigger in ALS ^64,65^, via Sodium Arsenite (NaARS) treatment. Longitudinal live imaging showed that both SOD1_G85S_ and TDP-43_M337V_ MNs were significantly more vulnerable to this stress than healthy MNs (Figure S3L-M). To determine whether intrinsically high-SMN levels were also predictive of better survival in fALS, SOD1_G85S_ and TDP-43_M337V_ MNs were live-imaged and classified by their SMN-Clover expression level. Similarly to the healthy MNs, high-SMN expressors were significantly more resistant to degeneration in both fALS backgrounds (Figure 2E-F). Analysis of an SMN survival threshold in fALS MNs revealed that the low-SMN expressors in both fALS genetic backgrounds had 60% less SMN compared to their high-SMN counterparts, a pattern similar to healthy BJ SMN-Clover MNs (Figure 2G). Thus, endogenous SMN abundance predicts survival not only in healthy MNs but also across mechanistically distinct MNDs, indicating that intrinsic SMN-dependent vulnerability represents a shared feature of MN degeneration.

We then explored whether endogenous SMN protein levels predicted MN survival in SMA and fALS selectively or also in CNs. BRN2^+^ SMA CNs were generated at similar rates across lines (Figure S4A-B). Fixation followed by SMN staining and single-cell unbiased quantification, as described above for SMA MNs, revealed that SMA CNs did not show SMN-dependent survival despite harboring SMN protein heterogeneity profiles analogous to the MN cultures (Figure S4C-D). Additionally, and like WT BJ CNs, SMA CNs showed a mild increased in average SMN expression (Figure S4D-F). SMN threshold analysis of SMA CNs showed that these cultures had the same proportion and expression patterns as MNs but still lack the SMN-dependent survival phenotype (Figure S4G). In contrast to MNs, neither SMA nor ALS CNs exhibited a strong relationship between endogenous SMN abundance and survival despite displaying comparable SMN heterogeneity (Figure S4). Therefore, the predictive value of endogenous SMN abundance is largely restricted to selectively vulnerable MNs. All together, these findings indicate that natural cell-to-cell variation in endogenous SMN expression defines a continuum of intrinsic neuronal vulnerability and provides a unifying mechanism that helps explain selective MN degeneration across distinct MNDs.

### Motor neurons dynamically transition toward vulnerable or resilient states

Having established that endogenous SMN abundance predicts motor neuron survival, we next asked whether these pre-degenerative vulnerability states are fixed properties established during differentiation or instead reflect dynamic cellular states that individual neurons can transition between over time (Figure 3A). Under the first model, neurons would maintain a relatively constant level of SMN throughout their lifetime, and the progressive enrichment of high-SMN neurons observed in culture would simply result from preferential loss of vulnerable low-SMN neurons. Alternatively, individual MNs might actively modulate endogenous SMN expression, allowing transitions between vulnerable and resilient states (Figure 3A). To distinguish between these possibilities, we continuously tracked individual healthy and familial ALS SMN-Clover MNs and classified them according to changes in endogenous SMN abundance over time as SMN-increasing, SMN-constant, or SMN-decreasing (Figure 3B). Across all genetic backgrounds, approximately 30–40% of MNs progressively increased endogenous SMN abundance, whereas roughly half maintained relatively stable SMN levels (Figure 3C). In contrast, only a small fraction of neurons (5–10%) exhibited decreasing SMN levels, the majority of which subsequently underwent cell death during the imaging period (Figure 3B-C). Because declining SMN expression largely reflected neurons already undergoing degeneration, these cells were excluded from subsequent analyses.

**Figure 3.**
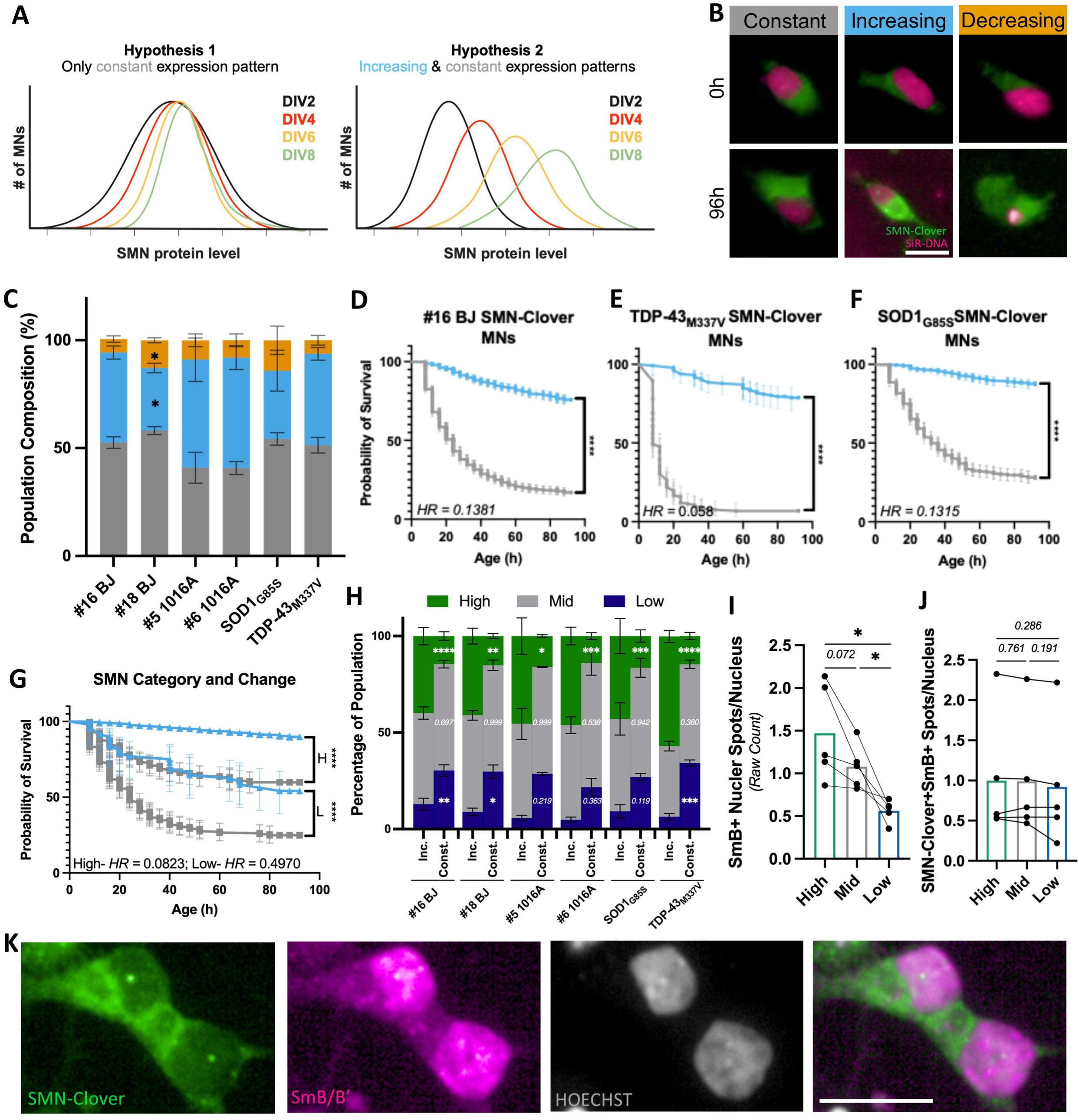
Dynamic increases in endogenous SMN define a resilient motor neuron state A. Schematic of two hypothesis of SMN intensity distribution changes overtime. Hypothesis 1 demonstrating a fixed level of SMN expression in cells and hypothesis 2 representing both constant and increasing expression patterns in cells. B. Representative #16 WT-BJ MNs in three dynamic SMN-expression categories at three timepoints with SMN-Clover in green and SiR-DNA in hot pink. Scale Bar = 10µM. C. Quantification of SMN-Clover MNs classified into each dynamic category. Significance was determined using RM two-way ANOVA with the Geisser-Greenhous correction and matched values and a Dunnett’s post hoc multiple comparisons test against #16 BJ. #16 BJ N=8; #18 BJ N=4; SOD1_G85S_ N=3; TDP-43_M337V_ N=4; WT-1016A N=3; #5 1016A N=2; #6 1016A N=3. Significance represented as: 0.1234 (ns), 0.0332 (*), 0.0021 (**), 0.0002 (***), <0.0001 (****). D. Representative Kaplan-Meyer survival plot of #16 BJ SMN-Clover MNs plotting the probability of survival of MNs classified by whether they increase or decrease their SMN-Clover intensity at each timepoint of the experiment (N=8). HR: hazard ratio between increasing- and constant-expressors. Error bars: 95% confidence interval. Comparisons between survival curves was done using the Log-rank (Mantel-Cox) test. Significance represented as: 0.1234 (ns), 0.0332 (*), 0.0021 (**), 0.0002 (***), <0.0001 (****). E. Representative Kaplan-Meyer survival plot of TDP-43_M337V_ SMN-Clover MNs (N=4). See (D). F. Representative Kaplan-Meyer survival plot of SOD1_G85S_ SMN-Clover MNs (N=3). See (D). G. Representative Kaplan-Meyer survival plot of #16 BJ SMN-Clover MNs where MNs are classified by whether the MN was first classified as a high- or low-expressor before SMN-Clover intensity increased or remained constant. H. Quantification of the percentage of SMN-Clover MNs first classified high- or low-expressors and whether SMN-Clover intensity increased or remained constant overtime. #16 BJ N=8; #18 BJ N=4; SOD1_G85S_ N=3; TDP-43_M337V_ N=4; WT-1016A N=3; #5 1016A N=2; #6 1016A N=3. Statistical significance calculated independently for each line between the increasing and constant MNs from each classification (high, mid, and low). An RM two-way ANOVA with matched high, mid, and low values was performed with a post hoc Turkey’s multiple comparisons test with single pooled variance. I. Quantification of SmB+ nuclear spots/nucleus in high, mid, and low #16 BJ SMN-Clover MNs (N=5). Significance was calculated using an RM one-way ANOVA with Geisser-Greenhouse correction and a post hoc uncorrected Fisher’s LSD test with variances computed for each comparison. Significance represented as: 0.1234 (ns), 0.0332 (*), 0.0021 (**), 0.0002 (***), <0.0001 (****). J. Quantification of SMN-Clover+SmB+ nuclear spots/nucleus in high, mid, and low #16 BJ SMN-Clover MNs (N=5). Significance was calculated using an RM one-way ANOVA with Geisser-Greenhouse correction and a post hoc uncorrected Fisher’s LSD test with variances computed for each comparison. Significance represented as: 0.1234 (ns), 0.0332 (*), 0.0021 (**), 0.0002 (***), <0.0001 (****).

We next asked whether increasing endogenous SMN abundance alters neuronal resilience. Kaplan-Meier survival analysis revealed that SMN-increasing MNs consistently survived significantly longer than neurons maintaining constant SMN expression, irrespective of genetic background (Figure 3D-F, S5). Importantly, this survival advantage was observed even after stratifying neurons according to their initial SMN abundance. Both high- and low-SMN neurons that subsequently increased endogenous SMN survived significantly longer than their respective constant-expression counterparts (Figure 3G). Motor neurons retain the capacity to transition from a vulnerable toward a more resilient state through endogenous SMN upregulation. Thus, MN vulnerability is not irreversibly predetermined but reflects a dynamic cellular state. These findings extend previous observations that increasing SMN abundance through gene replacement, splice modulation or genetic correction improves motor neuron survival ^28–30,37,38,51^ by demonstrating that a substantial proportion of motor neurons can spontaneously activate this protective mechanism without therapeutic intervention.

We next sought to identify mechanisms that might underlie endogenous SMN upregulation. Previous studies have proposed a positive feedback loop in which elevated SMN protein promotes increased *SMN2* exon 7 inclusion, thereby further enhancing full-length SMN production ^66,67^. Notably, neurons already expressing high endogenous SMN were nearly three times more likely to further increase SMN abundance than neurons initially expressing low SMN (Figure 3H). Specifically, approximately 40% of neurons initially classified as high-SMN continued to increase endogenous SMN expression during live imaging, whereas only ∼15% of neurons initially expressing low SMN were classified as SMN-increasing (Figure 3H). These observations are consistent with a self-reinforcing mechanism that stabilizes the resilient state. To investigate whether this resilient state is accompanied by enhanced spliceosomal maturation, as previously reported ^66,67^, we stained healthy #16 BJ SMN-Clover motor neurons for SmB (SNRPB), a core Sm protein assembled onto snRNAs by the SMN complex during snRNP biogenesis before incorporation into mature spliceosomal complexes ^23,68,69^. High-SMN neurons contained significantly more SmB+ nuclear puncta than low-SMN neurons, whereas the number of SMN-Clover/SmB double+ puncta remained unchanged (Figure 3I-J). These findings are consistent with enhanced SMN-dependent snRNP maturation and a previously proposed positive-feedback mechanism in which improved SMN-dependent splicing promotes increased *SMN2* exon 7 inclusion and sustained endogenous SMN production ^67,69^. Together, these findings demonstrate that MN resilience is not a fixed intrinsic property but rather that individual MNs retain the capacity to transition from a vulnerable toward a more resilient state through endogenous SMN upregulation.

### Low endogenous SMN defines a hyperexcitable neuronal state

Having established that endogenous SMN abundance defines dynamic MN vulnerability states, we next asked whether these states also differ in neuronal function. Motor neuron hyperexcitability has been reported in SMA mouse models and hiPSC-derived MNs ^9,70^, although whether this phenotype reflects a cell-autonomous consequence of reduced SMN within MNs or arises secondarily from alterations in the surrounding motor circuit remains controversial ^9,12,71,72^. We therefore investigated the relationship between endogenous SMN abundance and neuronal activity using a high-throughput calcium imaging platform that enabled automated single-cell quantification of MN activity in healthy and disease contexts (Figure 4A-B). All cell lines displayed spontaneous calcium transients that reflected functional neuronal activity, as demonstrated by the marked reduction in active cells following tetrodotoxin (TTX) treatment, the increased calcium peak amplitude following neuronal stimulation with 4-aminopyridine and bicuculline (4APBIC), and the presence of mature synaptic structures identified by PSD-95 and Synapsin co-localization (Figure S6A-D).

**Figure 4.**
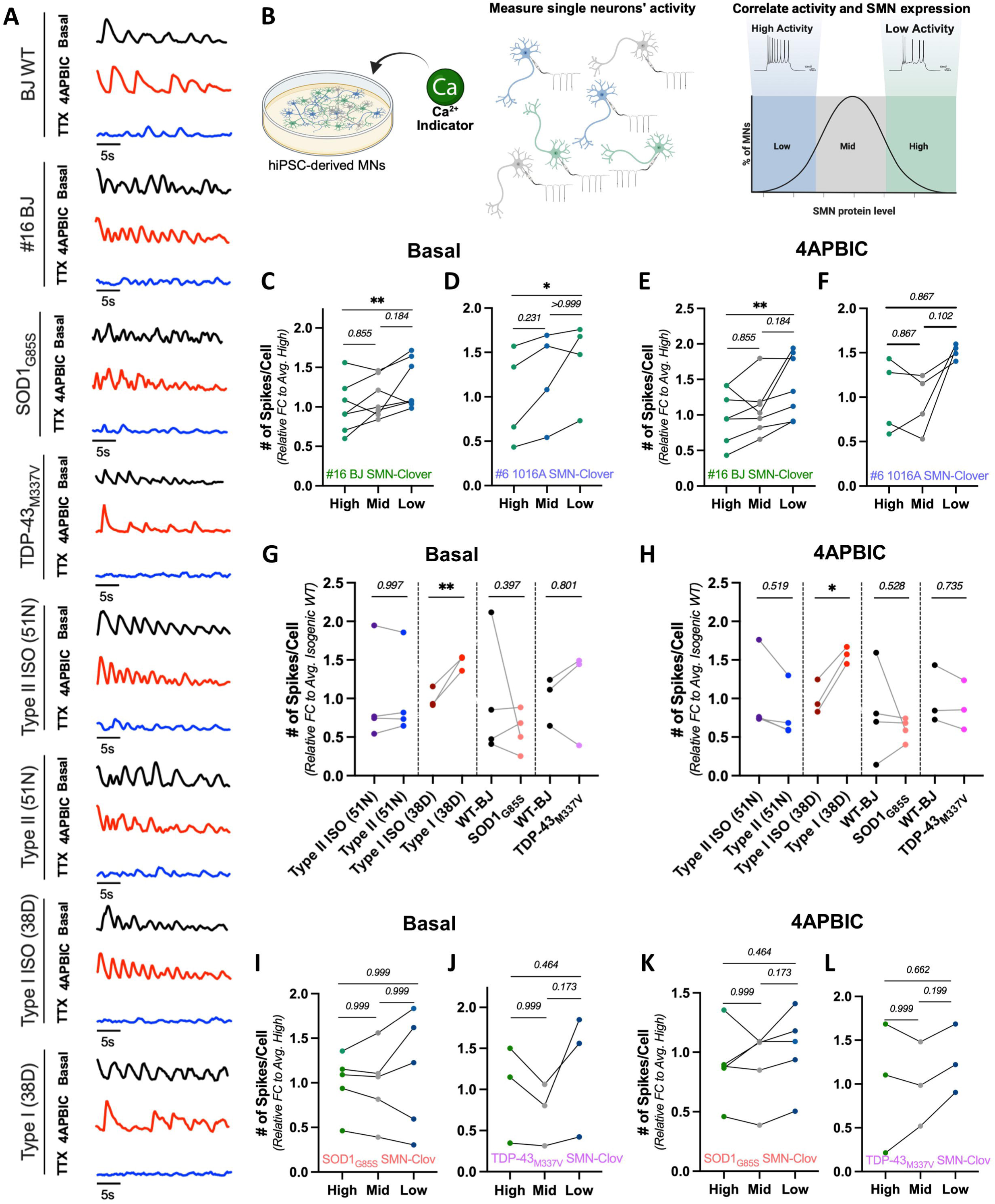
Endogenous SMN abundance predicts neuronal function A. Representative calcium traces of 1 cell/condition/cell line used in the study to demonstrate spiking activity. B. Schematic representation of measurement of calcium in individual hiPSC-derived MNs and the result that low-SMN expressors display higher levels of activity compared to high-SMN expressors. C. Quantification of spikes/cell in each SMN expression category relative to the average high-expressors from #16 BJ SMN-Clover MNs under basal conditions. Statistical significance was calculated using a pair Friedman test followed by a Dunn’s multiple comparisons post hoc test. Significance represented as: 0.1234 (ns), 0.0332 (*), 0.0021 (**), 0.0002 (***), <0.0001 (****). D. Same as (C) for #6 1016A SMN-Clover MNs under basal conditions. E. Same as (C) for #16 BJ SMN-Clover MNs under activating (4APBIC) conditions. F. Same as (C) for #6 1016A SMN-Clover MNs under activating (4APBIC) conditions. G. Quantification of spikes/cell normalized to the average isogenic WT for SMA and ALS isogenic pairs under basal conditions (Type II ISO (51N) N=4; Type II (51) N=4; Type I (38D) ISO N=3; Type I (38D) N=3; WT-BJ N=4; SOD1_G85S_ N=4; WT-BJ N=3; SOD1_G85S_ N=3). Statistical significance was calculated with an Ordinary one-way ANOVA followed by Tukey’s multiple comparisons post hoc test for each isogenic pair. Significance represented as: 0.1234 (ns), 0.0332 (*), 0.0021 (**), 0.0002 (***), <0.0001 (****). H. Same as (G) for activating (4APBIC) conditions). I. Same as (C) for SOD1_G85S_ SMN-Clover MNs under basal conditions. J. Same as (C) for TDP-43_M337V_ SMN-Clover MNs under basal conditions. K. Same as (C) for SOD1_G85S_ SMN-Clover MNs under activating (4APBIC) conditions. L. Same as (C) for TDP-43_M337V_ SMN-Clover MNs under activating (4APBIC) conditions.

To determine whether neuronal activity differs between resilient and vulnerable MN states, we compared high-, mid- and low-SMN-expressing motor neurons from the healthy #16 BJ and #6 1016A SMN-Clover lines. Under basal conditions, low-SMN MNs exhibited significantly greater spontaneous activity than high-SMN neurons in both genetic backgrounds (Figure 4C-D). Following pharmacological stimulation, low-SMN #16 BJ MNs remained significantly more excitable than high-SMN neurons, whereas low-SMN #6 1016A MNs showed the same trend without reaching statistical significance (Figure 4E-F). These findings demonstrate that reduced endogenous SMN abundance is associated with a hyperexcitable functional state, extending our previous observations that low-SMN neurons also exhibit reduced survival.

Because neuronal excitability is influenced by multiple intrinsic and extrinsic factors including soma size, dendritic morphology, synaptic connectivity and surrounding neuronal populations ^12,73–75^–-we next asked whether these variables could account for the relationship between endogenous SMN abundance and neuronal activity. Soma size did not differ between high-, mid- and low-SMN MNs in any SMN-Clover line analyzed (#16 BJ, #6 1016A, SOD1_G85S_ and TDP-43_M337V)_ (Figure S6E-H). Likewise, dendritic complexity, assessed by MAP2+ neurite staining, was indistinguishable between each pair of isogenic SMA lines (Figure S6I-J). Finally, although previous studies proposed that SMA MN hyperexcitability arises indirectly through the loss of proprioceptive interneuron inputs ^12^, the differentiation protocol used here has not been shown to generate proprioceptive interneurons capable of influencing MN activity ^76^. These observations indicate that the association between low endogenous SMN abundance and neuronal hyperexcitability is predominantly cell intrinsic.

We next asked whether endogenous SMN regulates neuronal excitability, or conversely whether differences in neuronal activity contribute to heterogeneous SMN expression. Intracellular calcium signaling has previously been implicated in regulating SMN abundance ^77–79^, raising the possibility that highly active neurons might secondarily alter endogenous SMN levels. To test this, #16 BJ MNs were exposed to prolonged activity blockade (TTX) or activation (4AP+BIC) under either continuous treatment conditions or following a two-day recovery period after compound washout (Figure S6E). Automated single-cell quantification revealed no significant change in endogenous SMN-Clover expression during continuous treatment, while only a modest (∼8%) reduction was observed after recovery from prolonged TTX exposure (Figure S6F). These experiments revealed that physiological modulation of neuronal activity has minimal impact on endogenous SMN abundance, supporting a model in which SMN acts upstream of altered neuronal excitability rather than being regulated by it.

To determine whether this relationship extends to disease models, we next analyzed neuronal activity in our isogenic SMA and familial ALS MNs. Restoration of SMN expression in the severe SMA Type I (38D) line significantly reduced MN activity under both basal and stimulated conditions, whereas the milder SMA Type II (51N) line showed no significant difference from its isogenic corrected counterpart (Figure 4G-H), suggesting that the impact of SMN on neuronal excitability is threshold dependent. We next asked whether endogenous SMN abundance similarly predicts neuronal activity in fALS, where both hyper- and hypoexcitability have been reported depending on disease stage and genetic background ^80^. Neither SOD1_G85S_ nor TDP-43_M337V_ MNs differed significantly from their isogenic controls, and stratification according to endogenous SMN abundance likewise revealed no differences in neuronal activity (Figure 4G-L). This may reflect the relatively immature developmental stage of our cultures ^81,82^, which was intentionally selected to permit longitudinal analysis of both vulnerable low-SMN and resilient high-SMN MNs before widespread degeneration occurs. Alternatively, the effects of ALS-causing mutations may dominate over any contribution of endogenous SMN to neuronal excitability. Together, these findings identify neuronal hyperexcitability as a functional feature of the vulnerable low-SMN state in healthy and SMA MNs and indicate that endogenous SMN abundance acts upstream of altered neuronal activity rather than being regulated by it.

### Intrinsic SMN-dependent vulnerability is conserved in vivo

Having established that endogenous SMN abundance defines dynamic motor neuron vulnerability states in vitro, we next asked whether this principle is also reflected during disease progression in vivo. Selective degeneration of spinal motor neuron populations is a hallmark of SMA in both patients and animal models ^83,84^, with lumbar L1/L2 motor neurons and medial motor column (MMC) neurons of the L4/L5 spinal cord being considerably more vulnerable than lateral motor column (LMC) neurons ^9,12,13^. We therefore investigated whether endogenous SMN abundance correlates with these established patterns of selective vulnerability using the SMNΔ7 mouse model (Smn^-/-^;hSMN2^+/+^;hSMNΔ7^+/+^;HB9:GFP^+^) and healthy littermate controls (Smn^+/-^;hSMN2^+/+^;hSMNΔ7^+/+^;HB9:GFP^+^). Spinal cords were collected before MN loss (P1), at disease onset (P4), during disease progression (P7/8) and at end-stage (P10/12), followed by immunostaining for SMN in the L1/L2 and L4/L5 spinal cord segments (Figure 5A-B, Figure S7A). Consistent with previous reports, vulnerable L1/L2 MNs and L4/L5 MMC MNs progressively degenerated throughout disease progression, whereas L4/L5 LMC MNs remained comparatively resistant (Figure 5C-E). Although absolute L1/L2 MN counts in healthy animals decreased modestly after P4, this reflects the sectioning and quantification strategy previously described by Sowoidnich et al. ^13^and does not affect comparisons between healthy and SMA mice. Thus, our analyses faithfully recapitulated the established pattern of selective motor neuron vulnerability in the SMNΔ7 model.

**Figure 5.**
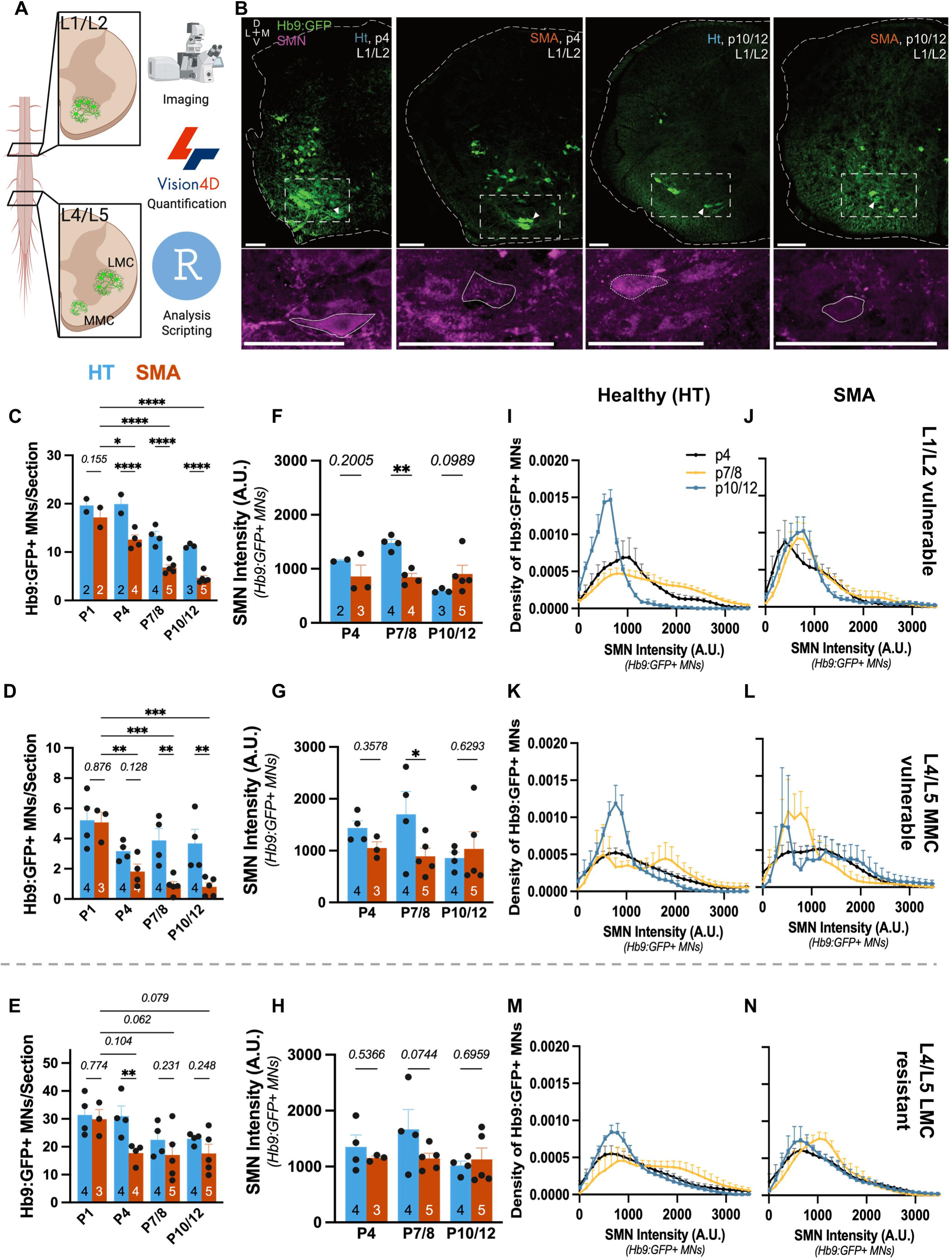
Intrinsic SMN-dependent vulnerability is conserved in vivo A. Schematic showing a representation of the mouse spinal cord and sections for the L1/L2 and L4/L5 segments of interest and the motor neuron distribution. Spinal cord sections were confocally imaged, motor neurons and SMN intensity were quantified using the ARIVIS Vision4D software, and then compiled using R-script based analysis. LMC: Lateral motor column. MMC: medial motor column. B. Representative images of p4 and p10/12 L1/L2 spinal cord segments. Upper image panels show outlines of spinal cord section and Hb9:GFP+ (green) MNs. Outlined box represents the MNs of interest in this segment and the below zoomed in region showing the SMN (magenta) staining. MNs are labelled with a dotted outline for clarity. Scale bar = 100µm. C. Quantification of vulnerable L1/L2 Hb9:GFP+ MNs/section in HT and SMA mice at one post-natal day (p1), p4, p7/8, and p10/12 ages. Each dot represents one mouse and the total number (N) is shown with a black and white number at the base of each bar. Statistical significance was calculated using an Ordinary two-way ANOVA with Tukey’s multiple comparisons post hot test. Significance represented as: 0.1234 (ns), 0.0332 (*), 0.0021 (**), 0.0002 (***), <0.0001 (****). D. Quantification of vulnerable L4/L5-MMC Hb9:GFP+ MNs/section in HT and SMA mice at one post-natal day (p1), p4, p7/8, and p10/12 ages. Same as (C). E. Quantification of resistant L4/L5-LMC Hb9:GFP+ MNs/section in HT and SMA mice at one post-natal day (p1), p4, p7/8, and p10/12 ages. Same as (C). F. Quantification of SMN Intensity (A.U.) in vulnerable L1/L2 Hb9:GFP+ MNs/section in HT and SMA mice at four post-natal days (p4), p7/8, and p10/12 ages. Each dot represents one mouse and the total number (N) is shown with a black and white number at the base of each bar. Statistical significance was calculated using an Ordinary two-way ANOVA with Tukey’s multiple comparisons post hot test. Significance represented as: 0.1234 (ns), 0.0332 (*), 0.0021 (**), 0.0002 (***), <0.0001 (****). G. Quantification of SMN Intensity (A.U.) in vulnerable L4/L5-MMC Hb9:GFP+ MNs/section in HT and SMA mice at one post-natal day (p1), p4, p7/8, and p10/12 ages. Same as (B). H. Quantification of SMN Intensity (A.U.) in resistant L4/L5-LMC Hb9:GFP+ MNs/section in HT and SMA mice at one post-natal day (p1), p4, p7/8, and p10/12 ages. Same as (B). I. Density plot showing a predictive proportion of vulnerable L1/L2 Hb9:GFP+ HT MNs that fall within a discrete level of SMN Intensity (A.U.) at p4 (N=2), p7/8 (N=4), and p10/12 (N=3). J. Same as (F) for vulnerable L4/L5-MMC Hb9:GFP+ HT MNs. K. Same as (F) for resistant L4/L5-LMC Hb9:GFP+ HT MNs. L. Density plot showing a predictive proportion of vulnerable L1/L2 Hb9:GFP+ SMA MNs that fall within a discrete level of SMN Intensity (A.U.) at p4 (SMA N=3), p7/8 (N=5), and p10/12 (N=5). M. Same as (I) for vulnerable L4/L5-MMC Hb9:GFP+ SMA MNs. N. Same as (I) for resistant L4/L5-LMC Hb9:GFP+ SMA MNs.

To address whether differences in endogenous SMN abundance accompany these distinct vulnerability profiles we measured population level SMN expression. Average SMN expression in vulnerable SMA motor neuron populations was reduced at P4 and significantly reduced at P7/8 compared with their healthy counterparts, whereas resistant LMC MNs maintained comparable SMN levels throughout disease progression (Figure 5F-H). By end-stage disease (P10/12), average SMN expression no longer differed significantly between healthy and SMA vulnerable MN populations, consistent with the preferential loss of low-SMN MNs during disease progression. These findings suggest that MNs maintaining relatively higher endogenous SMN are preferentially preserved in vivo, mirroring our observations in cultured MNs. Because average protein measurements cannot distinguish whether these changes reflect uniform SMN reduction across all neurons or selective loss of low-SMN cells, we next quantified endogenous SMN abundance at single-cell resolution within each motor neuron population. Similar to our in vitro observations, healthy and SMA MNs exhibited remarkably broad endogenous SMN distributions spanning approximately a four- to five-fold range of expression levels (Figure 5I-N). Importantly, the SMN distributions of vulnerable SMA MN populations progressively shifted toward higher SMN expression during disease progression, with a marked enrichment of high-SMN neurons at P7/8 and P12 compared with P4, although this shift was less pronounced than that observed in vitro (Figure 5J, L). In contrast, resistant LMC MNs maintained comparatively stable SMN distributions throughout disease progression (Figure 5N). Interestingly, the highly vulnerable L4/L5 MMC population exhibited a bimodal SMN distribution at end-stage disease, with a small population of neurons persisting at relatively low SMN levels while the majority clustered within a substantially higher SMN expression range (Figure 5M), suggesting that only a subset of low-SMN neurons escapes degeneration. In summary, the longitudinal analysis of MN survival, average endogenous SMN abundance and single-cell SMN distributions demonstrates that the intrinsic SMN-dependent vulnerability states identified by live-cell imaging in vitro are conserved during SMA disease progression in vivo. Across both experimental systems, MNs maintaining relatively high endogenous SMN are preferentially preserved, whereas neurons expressing lower SMN exhibit a substantially greater probability of degeneration.

### Distinct molecular programs characterize resilient and vulnerable motor neuron states

Having established that endogenous SMN abundance defines dynamic motor neuron vulnerability states both in vitro and in vivo, we next sought to identify the molecular programs that distinguish resilient from vulnerable MNs. To identify molecular determinants of intrinsic MN resilience independent of genetic background, we compared genetically identical MNs that differed primarily in their endogenous SMN abundance. This experimental design provides a uniquely controlled comparison in which differences in gene and protein expression are associated with the resilient and vulnerable states themselves, rather than with disease-causing mutations or differences in cellular identity. To this end, SMN-Clover MNs derived from two independent healthy BJ hiPSC clones (#16 and #18) were FACS-sorted into high-, mid- and low-SMN populations at an early (DIV4-5) and a late (DIV12) developmental stage spanning the transition in endogenous SMN-defined vulnerability states (Figure 6A). Sorted populations were subsequently analyzed by bulk RNA sequencing and quantitative mass spectrometry. FACS sorting reliably separated MNs according to endogenous SMN abundance, as confirmed by western blotting, *SMN1* transcript abundance and SMN peptide intensity (Figure S8A, Figure 6B-C). Differential expression analysis identified 99 differentially expressed genes (DEGs) and 369 differentially expressed proteins (DEPs) between high- and low-SMN MNs across both developmental stages (Figure 6D). As expected, most molecular differences were stage-specific, with only 14 DEGs and 21 DEPs shared between the early and late time points (Figure 6D-E). Hierarchical clustering of the transcriptomic data further demonstrated reproducible separation of high- and low-SMN MNs at both developmental stages (Figure S8B), whereas manual functional annotation grouped the conserved DEGs into biological categories including neuronal identity, neuronal maturation, RNA processing, cytoskeletal organization, metabolism/autophagy, neuronal activity and cell survival (Figure 6F, S8C, Supplementary Table 1). Because proteins represent the functional effectors of cellular state, we next analyzed the proteomic dataset. Volcano plots confirmed robust differential protein expression between high- and low-SMN MNs at both developmental stages (Figure S8D). Functional network analysis of the 369 DEPs identified 38 interconnected biological modules encompassing pathways involved in cytoskeletal organization, neuronal differentiation, glutamatergic signaling, autophagy, metabolism and stress responses (Figure 6G, S9 and Supplementary Table 2-3). To facilitate biological interpretation of the conserved proteomic signature, overlapping proteins were further classified according to the consistency of their regulation across clones and developmental stages (Supplementary Table 2; see *Star Methods*). This analysis identified a directionally concordant core proteomic signature enriched for proteins involved in RNA regulation, proteostasis, neuronal differentiation, metabolism and stress signaling. Importantly, both the transcriptomic and proteomic datasets independently converged on biological processes previously implicated in selective MN vulnerability in SMA and ALS, including RNA metabolism, proteostasis, cytoskeletal regulation and neuronal activity (Figure 6F-G) ^7,18–21,85–89^. Thus, despite relatively modest molecular differences between high- and low-SMN MNs, both datasets independently identified coordinated molecular programs associated with the resilient and vulnerable states.

**Figure 6.**
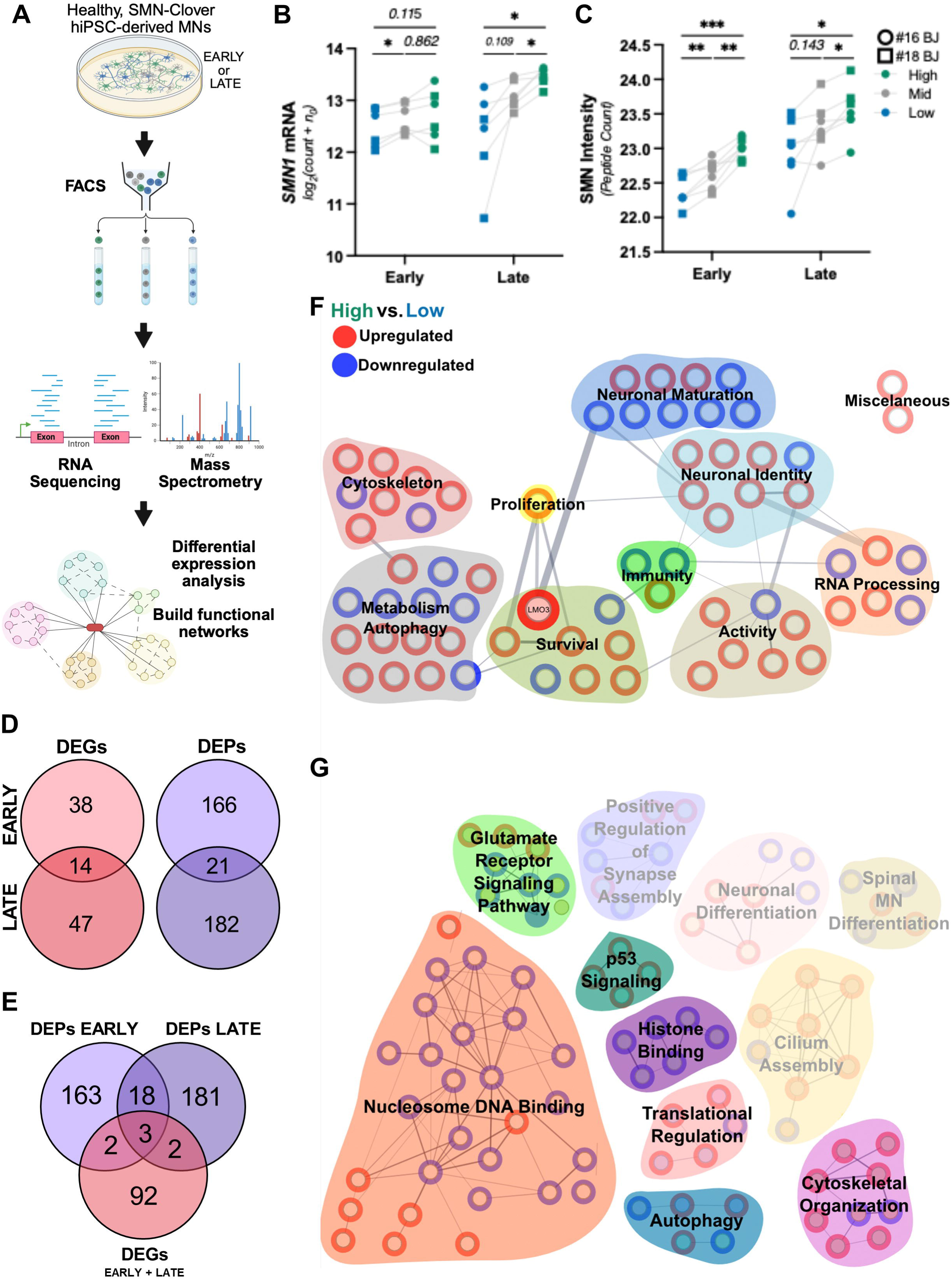
Distinct molecular programs underlie resilient and vulnerable motor neuron states A. Schematic of FACS experiment to sort WT SMN-Clover MNs into high-, mid-, and low-expresser categories based upon fluorescence measurements and then downstream processing of bulk RNA-sequencing or proteomics. B. Quantification of *SMN1* mRNA measured by bulk RNA-sequencing and expressed as the log2(count + n_0_) of sequencing reads (Early: #16 BJ N=3, #18 BJ N=3; Late: #16 BJ N=3, #18 BJ N=3). Statistical significance was calculated using an RM one-way ANOVA with the Geisser-Greenhouse correction and a Tukey’s multiple comparison post hoc test. Each time point was assessed separately with #16 BJ and #18 BJ replicates pooled as was done in the DEG analysis. Significance represented as: 0.1234 (ns), 0.0332 (*), 0.0021 (**), 0.0002 (***), <0.0001 (****). C. Quantification of SMN protein measured by bulk mass spectrometry and expressed as the number of peptide counts (Early: #16 BJ N=4, #18 BJ N=3; Late: #16 BJ N=3, #18 BJ N=3). Statistical significance was calculated using an RM one-way ANOVA with the Geisser-Greenhouse correction and a Tukey’s multiple comparison post hoc test. Each time point was assessed separately with #16 BJ and #18 BJ replicates pooled as was done in the DEG analysis. Significance represented as: 0.1234 (ns), 0.0332 (*), 0.0021 (**), 0.0002 (***), <0.0001 (****). D. Venn diagram of significantly differentially expressed genes (DEGs, red) and differentially expressed proteins (DEPs, gray) identified when comparing high- and lows-expressors at the early and late time points. E. Venn diagram showing the overlap between DEGs (both time points), early DEPs, and late DEPs. F. String diagram of DEGs between high- and low-expressing SMN-Clover MNs. Each node represents one DEG and is colored by whether they were upregulated (red) or downregulated (down) in the high-expressers. DEGs were hand classified into functional categories represented here by colored shapes and names. G. String diagram of the most interesting functional clusters determined from the DEPs between high- and low-expressing SMN-Clover MNs. Upper left hand corner shows the entire string diagram. Each node represents one DEP and is colored by whether they were upregulated (red) or downregulated (down) in the high-expressers. DEPs were classified using StringDB and associated GO Terms, KEGG pathways, and Reactome pathways. Highlighted networks represent biological processes discussed in the main text.

Among the conserved transcriptional changes, several genes reproducibly distinguished high- and low-SMN MNs across developmental stages, including genes involved in neuronal identity (*ISL1*, *HMX3*), neuronal activity (*SNCA*), cytoskeletal organization (*DLC1*) and transcriptional regulation (*NFATC1*) (Supplementary Table 1). Because our objective was to identify candidate regulators of MN resilience, we next focused on genes with established links to neuronal survival. Among these candidates, *LMO3* was unique in having a direct reported role in suppressing p53-mediated apoptosis ^90^. Together with the consistent enrichment of LMO3 in high-SMN MNs at both developmental stages, this made it a compelling candidate regulator of the resilient MN state for subsequent functional validation. Consistent with the transcriptional enrichment of survival-associated genes, proteomic network analysis independently identified a p53-signalling module (Figure 6G, S8). Among the proteins contributing to this network was MDM4, a regulator whose aberrant splicing under SMN-deficient conditions has previously been shown to activate p53-mediated degeneration in selectively vulnerable MNs ^91^. Together, these findings support the notion that differential regulation of stress-response pathways contributes to intrinsic MN resilience. In parallel, both transcriptomic and proteomic analyses identified coordinated changes in pathways regulating neuronal activity. These molecular alterations complement our functional observations that low-SMN MNs exhibit increased spontaneous activity (Figure 4) and further support the concept that neuronal excitability represents an integral component of the vulnerable MN state rather than an independent phenotype. Collectively, these integrated transcriptomic and proteomic analyses demonstrate that resilient and vulnerable MNs occupy distinct molecular states enriched for pathways controlling survival, RNA metabolism and neuronal activity, and identify LMO3 as a candidate regulator of MN resilience.

### LMO3 promotes a resilient motor neuron program

Having identified LMO3 as one of the few genes consistently enriched in high-SMN motor neurons at both early and late developmental stages (Figure 6, Supplementary Table 1), we next investigated whether increasing LMO3 expression is sufficient to promote features of the resilient MN state. LMO3 has previously been reported to inhibit p53-dependent apoptosis through repression of p53 transcriptional activity ^90^, making it an attractive candidate mediator of the survival-associated molecular program identified in our transcriptomic analyses. We therefore generated lentiviral vectors expressing either mCherry alone (control) or LMO3-P2A-mCherry and transduced healthy (#16 BJ) and SMA type I (33A) embryoid bodies during the onset of MN differentiation (EB9), followed by analysis after six days of transgene expression (EB15) (Figure 7A). To verify pathway engagement, we first quantified LMO3 together with two downstream p53-responsive genes. LMO3 overexpression markedly increased LMO3 transcript abundance while reducing expression of the pro-apoptotic genes CDKN1A and PMAIP1 in both healthy and SMA cultures, consistent with suppression of p53-dependent apoptotic signaling (Figure 7B-C).

**Figure 7.**
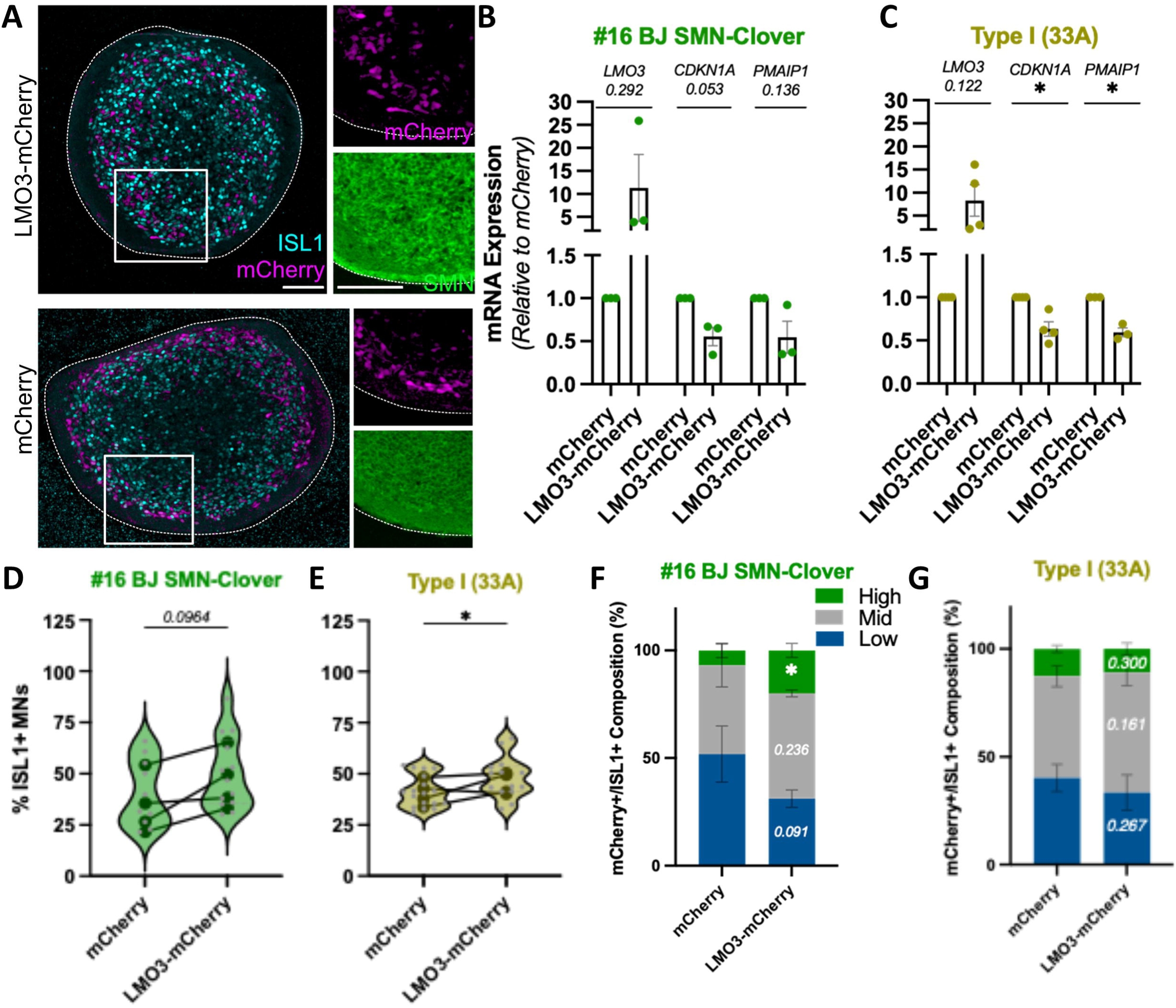
LMO3 promotes a resilient motor neuron program A. Representative images of D15 #16 BJ embryoid bodies transduced with mCherry or LMO3-mCherry lentiviruses. Images show mCherry (gray), ISL1 (cyan), SMN (magenta). Scale bar = 100µm for whole image and 50µm for zooms. B. mRNA expression of *LMO3* and p53 pro-apoptotic target genes (*CDKA1* and *PMAIP1*) in #16 BJ SMN-Clover embryoid bodies (D15) transduced with control mCherry or LMO3-mCherry lentiviruses and allowed to express each construct for 6 days (N=3). Statistical significance was calculated for each gene using a parametric, one-tailed paired t-test. Significance represented as: 0.1234 (ns), 0.0332 (*), 0.0021 (**), 0.0002 (***), <0.0001 (****). C. Same as (B) for Type I (33A) embryoid bodies (D15) (N=4). D. Quantification of ISL1+ MNs in D15 embryoid bodies transduced with mCherry or LMO3-mCherry lentiviruses (N=4, n=1-6). Violin plots represent the % of ISL1+ MNs in each organoid (small gray points) while each point is an average of all embryoid bodies in a biological replicate. Statistical significance was calculated against all EBs in each condition via a parametric, one-tailed unpaired t-test. Significance represented as: 0.1234 (ns), 0.0332 (*), 0.0021 (**), 0.0002 (***), <0.0001 (****). E. Same as (D) for Type I (33A) embryoid bodies (D15) (N=5, n=1-6). F. Quantification of mCherry+/ISL1+ MNs classified as High, Mid, and Low repressors in D15 EBs transduced with mCherry or LMO3-mCherry lentiviruses (N=4, n=1-6). Error bars represent SEM. Statistical significance was calculated between all EBs of each class across each condition via a parametric, one-tailed unpaired t-test. Significance represented as: 0.1234 (ns), 0.0332 (*), 0.0021 (**), 0.0002 (***), <0.0001 (****). G. Same as (F) for Type I (33A) embryoid bodies (D15) (N=4, n=1-6). H. Schematic representation of our hypothesis where high SMN expression is representative of a resilient MN program exemplified by the protective effect of LMO3 overexpression.

To determine whether activation of this program influences MN survival, cryosections from control- and LMO3-transduced EBs were stained for ISL1 and the proportion of ISL1+ MNs was quantified across four independent differentiations. LMO3 overexpression increased the proportion of ISL1+ MNs in both genetic backgrounds, reaching statistical significance in SMA cultures while showing the same trend in healthy cultures (Figure 7D-E). These findings demonstrate that activation of LMO3 is sufficient to promote maintenance of MNs during the developmental period when vulnerable neurons are normally lost.

Finally, we asked whether LMO3 influences the endogenous SMN-defined vulnerability states identified throughout this study. Endogenous SMN abundance was quantified specifically within mCherry+;ISL1+ MNs following LMO3 overexpression. In healthy cultures, LMO3 reduced the proportion of low-SMN MNs while increasing the representation of high-SMN neurons (Figure 7F), with a similar trend emerging in SMA cultures (Figure 7G). Importantly, this shift occurred simultaneously with an increase in the total number of surviving ISL1+ MNs (Figure 7D-E), arguing against a simple enrichment of high-SMN neurons through preferential loss of low-SMN cells. Instead, these findings are most consistent with LMO3 promoting the acquisition or stabilization of a higher SMN, resilient MN state. Collectively, these experiments provide functional validation of the molecular analyses presented in Figure 6 and demonstrate that LMO3 is not only associated with the higher SMN molecular states but is sufficient to induce key features of that program, including suppression of pro-apoptotic signaling, increased MN survival and enrichment of high endogenous SMN states.

## Discussion

Selective neuronal vulnerability is a defining feature of neurodegenerative disease, yet why genetically identical neurons exposed to the same pathogenic environment exhibit markedly different probabilities of degeneration remains one of the central unresolved questions in neuroscience. Here, we demonstrate that endogenous SMN abundance defines dynamic motor neuron vulnerability states that predict both neuronal function and survival across healthy, spinal muscular atrophy (SMA) and amyotrophic lateral sclerosis (ALS) genetic backgrounds. By combining longitudinal single-cell imaging, functional phenotyping, in vivo validation and integrated transcriptomic and proteomic analyses, we show that naturally occurring variation in endogenous SMN abundance identifies biologically meaningful cellular states that predict neuronal fate, transition over time and can be manipulated to promote neuronal resilience. Together, these findings establish endogenous protein heterogeneity as a framework for identifying intrinsic mechanisms of selective neuronal vulnerability.

Our study substantially extends previous observations linking endogenous SMN abundance to MN survival. Earlier work demonstrated that neurons expressing relatively high endogenous SMN are more resistant to experimentally induced stress ^38^. Here, we show that endogenous SMN predicts spontaneous survival in the absence of exogenous insults, that this relationship is conserved across healthy, SMA and ALS genetic backgrounds, and that it is recapitulated in selectively vulnerable MN populations in vivo. Importantly, longitudinal imaging revealed that endogenous SMN-defined states are dynamic rather than fixed. A substantial proportion of neurons progressively increased endogenous SMN abundance and displayed enhanced survival, whereas neurons exhibiting constant or declining SMN preferentially degenerated. Thus, endogenous SMN abundance should not be viewed simply as a biomarker of vulnerability but rather as a dynamic cellular state that influences neuronal fate.

The dynamic nature of these states has important implications for our understanding of selective neuronal vulnerability. Vulnerability is often considered a property imposed by disease-causing mutations ^2,3^; however, our data suggest that genetically identical MNs already occupy different resilience states before degeneration becomes apparent. This concept is reinforced by recent work demonstrating that vulnerable α-MNs in an ALS mouse model transition into a distinct disease-associated motor neuron state before cell death, with transcriptional regulatory networks governing this transition and related features conserved in human ALS ^92^. Together with our findings, these observations support an emerging view in which motor neuron vulnerability reflects dynamic cellular states that precede overt degeneration rather than simply the terminal consequences of disease. Importantly, whereas Gautier et al. identified disease-associated state transitions during ALS progression, our results reveal naturally occurring vulnerability and resilience states among otherwise equivalent MNs before degeneration and demonstrate that individual neurons can transition between SMN-defined states over time. These observations are also consistent with developmental priming models proposing that subtle differences established during neuronal specification influence later disease susceptibility ^51,93–95^. Rather than representing irreversible neuronal identities, our data indicate that these states remain plastic during maturation, allowing individual neurons to transition toward increased resilience.

One potential mechanism underlying this plasticity is a positive-feedback loop regulating endogenous SMN abundance. Previous studies demonstrated that efficient SMN-dependent snRNP assembly promotes *SMN2* exon 7 inclusion, thereby increasing full-length SMN production ^66,67^. Consistent with this model, neurons that initially expressed relatively high endogenous SMN were considerably more likely to further increase SMN abundance during longitudinal imaging, and high-SMN neurons exhibited increased numbers of mature SmB+ nuclear structures. Although our experiments do not directly establish causality, together with previous work these findings support the existence of a self-reinforcing mechanism capable of stabilizing resilient motor neuron states.

Our functional analyses further demonstrate that endogenous SMN abundance predicts not only survival but also neuronal physiology. Low-SMN MNs exhibited increased spontaneous activity despite being genetically identical to neighboring high-SMN neurons. Previous studies associated reduced SMN with MN hyperexcitability in SMA models ^9,12,70–72^; our work extends those observations by demonstrating that excitability varies according to endogenous SMN abundance within homogeneous neuronal populations. Whether altered excitability actively contributes to degeneration or instead represents an early manifestation of the vulnerable state remains unresolved, but the concordance between calcium imaging and molecular profiling suggests that altered neuronal activity forms part of a broader vulnerability program.

Transcriptomic and proteomic analyses further demonstrated that resilient and vulnerable MNs occupy distinct molecular states. Although the molecular differences between high- and low-SMN neurons were relatively modest, both datasets independently converged on pathways regulating RNA metabolism, proteostasis, cytoskeletal organization, neuronal activity, metabolism and stress responses, all processes repeatedly implicated in motor neuron disease ^7,19,20,85–89^. Importantly, these comparisons were performed between genetically identical neurons differing primarily in endogenous SMN abundance, minimizing confounding effects of developmental identity or disease-causing mutations and allowing intrinsic resilience programs to be resolved with unique precision. Among the conserved transcriptional changes, LMO3 emerged as a particularly compelling candidate because it was consistently enriched in high-SMN neurons at both developmental stages and has a documented role in suppressing p53-dependent apoptosis. Functional validation demonstrated that LMO3 overexpression reduced expression of canonical p53 target genes, increased the abundance of surviving ISL1+ MNs and shifted endogenous SMN distributions toward the higher SMN state. Together, these findings indicate that molecular programs associated with endogenous resilience are not merely correlative but can be functionally manipulated to promote motor neuron survival.

Our study has several limitations. The molecular analyses were performed in developing human MNs, and the relative contribution of individual pathways may evolve during long-term maturation. Likewise, although endogenous SMN-defined vulnerability states were conserved across SMA and ALS models, additional disease-specific mechanisms undoubtedly shape neuronal degeneration at later stages. Indeed, the emergence of disease-associated MN states during ALS progression ^92^ suggests that the intrinsic vulnerability states identified here may subsequently interact with disease-induced transcriptional and epigenetic programs. Future longitudinal epigenetic and single-cell multi-omic studies will be important to define how vulnerability states are established during development, maintained throughout maturation and remodeled during disease progression.

More broadly, our findings suggest a conceptual shift in how selective neuronal vulnerability can be studied. Rather than treating naturally occurring cell-to-cell variability as experimental noise, endogenous protein heterogeneity can be exploited to identify intrinsic resilience mechanisms within genetically homogeneous neuronal populations. This strategy reveals molecular programs that distinguish neurons destined to survive from those predisposed to degenerate while avoiding many of the confounding variables inherent to comparisons between anatomically distinct neuronal subtypes. We propose that endogenous protein heterogeneity is not merely a source of biological variability but a general experimental strategy for discovering intrinsic resilience mechanisms across neurodegenerative diseases. Ultimately, transforming endogenous cellular variability from an experimental nuisance into a biological discovery tool may provide new opportunities for identifying therapeutic targets capable of enhancing neuronal resilience before irreversible degeneration occurs.

## Resource Availability

### Data and Code Availability

All data supporting the findings of this study are available from the corresponding author upon reasonable request. Raw bulk RNA-sequencing and mass spectrometry proteomics data are being deposited in the European Genome-phenome Archive (EGA) and the PRIDE Proteomics Identifications Database, respectively. Accession numbers will be provided as soon as they are issued and prior to publication. Accession numbers are currently pending and will be provided as soon as they are issued and, in all cases, prior to publication.

## Materials Availability

Materials used in this project are listed in the *Reagents and Tools Table* and any materials generated by the lab are available upon request with appropriate MTA approvals.

## Code Availability

The code used in this study is available on Github (https://github.com/Rodriguez-MuelaLab). This includes the live longitudinal image analysis pipeline (R), calcium imaging pipeline (KNIME), script for analysis of mouse spinal cord data (R), script for analysis of EB data (R), and macros for EB quantification (FIJI).

## Acknowledgements

We thank Lee Rubin (Harvard University) for providing the MTAs enabling the work with the WT and SMA hiPSC lines. We thank all members of the DRESDEN-concept Genome Center (DcGC) (CMCB Technology Platform, TUD) for performing the bulk RNA-seq experiments and analysis. We thank Dr. Silke White (DZNE Imaging Platform) for her support and training on confocal imagine. We thank Sandra Günther, Jens Bergmann, and all animal users of the DZNE Animal Facility for their assistance in maintaining our mouse colony. We thank the CECAD Mass Spectrometry Facility in Cologne for performing the bulk proteomics experiment. We thank Dr. Martin Stöter and Dr. Rico Barsacchi of the MPI-CBG Technology Development Studio (TDS) in developing our calcium imagine analysis pipeline, experimental design and imaging with the Yakogawa CV7000. We thank Julia Jarrels and Jessica Hernández of the MPI-CBG Cell Technologies facility for their support in FACS sorting our hiPSC-derived MNs and confirming our RNA integrity for the bulk transcriptomics experiments. We thank Mike Karl (DZNE) for access to the Leica Cryostat and Michael Siewecke (CRTD) for access to the Applied Biosystem qPCR machine. We thank all current and past members of the Rodriguez-Muela Laboratory for their contributions and insightful discussions.

## Author Contributions

Conceptualization of project: NRM, JT; Data acquisition: JT, AR, JY, AC, SV; Data Analysis: JT, AR, JY, FR, SV; Funding acquisition: NRM, BW; Primary supervision: NRM, TG, BW; Writing: JT, NRM; Review & Editing: NRM, JT, TG and BW.

## Supplemental Information

Document S1. Supplemental Figures S1-S9, Supplemental Tables 1-2 Document S2. Supplemental Table 3

## Supplemental Figure Legends

**Supp. Figure 1.** Healthy high-SMN expressors are more likely to survive

A. Immunoblot of EB15 protein lysates from the indicated lines probed for SMN-Clover, untagged SMN, and beta-Tubulin (N=3/line).

B. Schematic of protocol to generate hiPSC-derived MNs using small molecule patterning. Representative images of hiPSCs, embryoid bodies, and dissociated, plated hiPSC-derived MNs.

C. Percentage of ISL1+ MNs at DIV6 of dissociated hiPSC-derived MN cultures. Each point represents a biological replicate. WT-BJ N=11; #16 BJ N=7; #18 BJ N=4; SOD1_G85S_ N=5; TDP-43_M337V_ N=3; WT-1016A N=3; #5 1016A N=3; #6 1016A N=3.

D. Relative # of ISL1+ MNs from DIV2 to DIV10 of hiPSC-derived MN cultures. Statistical comparisons were calculated between DIV2 to DIV10 within each line using a Mixed-Effects Analysis with the Geisser-Greenhouse correction and the Uncorrected Fisher’s LSD post-hoc test. Significance represented by the color of each cell line: 0.1234 (ns), 0.0332 (*), 0.0021 (**), 0.0002 (***), <0.0001 (****). WT-BJ N=11; #16 BJ N=7; #18 BJ N=4; SOD1_G85S_ N=5; TDP-43_M337V_ N=3; WT-1016A N=3; #5 1016A N=2; #6 1016A N=3.

E. Relative SMN Intensity (A.U.) in ISL1+ MNs from DIV2 to DIV10 of hiPSC-MN cultures. Statistics compare DIV2 to DIV10 within each line using a Mixed-Effects Analysis with the Geisser-Greenhouse correction and the Uncorrected Fisher’s LSD post-hoc test. Significance represented by the color of each cell line: 0.1234 (ns), 0.0332 (*), 0.0021 (**), 0.0002 (***), <0.0001 (****). WT-BJ N=11; #16 BJ N=7; #18 BJ N=4; SOD1_G85S_ N=5; TDP-43_M337V_ N=3; WT-1016A N=3; #5 1016A N=3; #6 1016A N=3.

F. Relative SMN-Clover Intensity (A.U.) in ISL1+ MNs from DIV2 to DIV10 of hiPSC-MN cultures. Statistics compare DIV2 to DIV10 within each line using a Mixed-Effects Analysis with the Geisser-Greenhouse correction and the Uncorrected Fisher’s LSD post-hoc test. Significance represented by the color of each cell line: 0.1234 (ns), 0.0332 (*), 0.0021 (**), 0.0002 (***), <0.0001 (****). WT-BJ N=11; #16 BJ N=7; #18 BJ N=4; SOD1_G85S_ N=5; TDP-43_M337V_ N=3; WT-1016A N=3; #5 1016A N=3; #6 1016A N=3.

G. Representative SMN Intensity (A.U.) histogram of WT-BJ, hiSPC-derived ISL1+ MNs at time points from DIV2 to DIV10.

H. Same as (G) for WT-1016A.

I. Representative SMN (left) and SMN-Clover (right) Intensity (A.U.) histogram of #18 BJ SMN-Clover, hiSPC-derived ISL1+ MNs at time points from DIV2 to DIV10.

J. Same as (I) for #5 1016A SMN-Clover.

K. Representative Kaplan-Meyer survival plot of #18 BJ SMN-Clover MNs plotting the probability of survival of MNs classified by their SMN-Clover expression level at each time point of the experiment (N=4). HR: hazard ratio between low- and high-expressors. Error bars: 95% confidence interval. Comparisons between survival curves was done using the Log-rank (Mantel-Cox) test. Significance represented as: 0.1234 (ns), 0.0332 (*), 0.0021 (**), 0.0002 (***), <0.0001 (****).

L. Same as (K) for #5 1016A SMN-Clover (N=2).

M. Schematic of protocol to generate hiPSC-derived cortical neurons using lentiviral transduction of a TetOn NGN2 overexpression vector and small molecule support. Representative images of hiPSCs undergoing differentiation into cortical neurons in 2D.

N. Percentage of BRN2+ CNs at DIV10 of lentiviral derived hiPSC-CN cultures. Each point represents a biological replicate. WT-BJ N=6; #16 BJ N=6; #6 1016A N=3; SOD1_G85S_ N=8; TDP-43_M337V_ N=3.

O. Relative # of BRN2+ CNs from DIV10 to DIV14 of hiPSC-derived CN cultures. Statistical comparisons were calculated between DIV10 to DIV14 within each line using a Mixed-Effects Analysis with the Geisser-Greenhouse correction and the Uncorrected Fisher’s LSD post-hoc test. Significance represented by the color of each cell line: 0.1234 (ns), 0.0332 (*), 0.0021 (**), 0.0002 (***), <0.0001 (****). WT-BJ N=6; #16 BJ N=3; #6 1016A.

P. Relative SMN Intensity (A.U.) in BRN2+ CNs from DIV10 to DIV14 of hiPSC-derived CN cultures. Statistical comparisons were calculated between DIV10 to DIV14 within each line using a Mixed-Effects Analysis with the Geisser-Greenhouse correction and the Uncorrected Fisher’s LSD post-hoc test. Significance represented by the color of each cell line: 0.1234 (ns), 0.0332 (*), 0.0021 (**), 0.0002 (***), <0.0001 (****). WT-BJ N=6; #16 BJ N=3; #6 1016A.

Q. Relative SMN-Clover Intensity (A.U.) in BRN2+ CNs from DIV10 to DIV14 of hiPSC-derived CN cultures. Statistical comparisons were calculated between DIV10 to DIV14 within each line using a Mixed-Effects Analysis with the Geisser-Greenhouse correction and the Uncorrected Fisher’s LSD post-hoc test. Significance represented by the color of each cell line: 0.1234 (ns), 0.0332 (*), 0.0021 (**), 0.0002 (***), <0.0001 (****). #16 BJ N=3; #6 1016A.

R. Representative SMN Intensity (A.U.) histogram of WT-BJ, lentiviral derived hiPSC-CN cultures at DIV10.

**Supp. Figure 2.** SMN degradation does not result in variable SMN expression levels

A. Representative images of DIV 8 WT-BJ, MNs stained for SMN (orange), p62 (purple), CULLIN5 (teal), and HOECHST (gray).

B. Quantification of SMN Intensity (A.U.) in SMN expression categories determined independent for each line at DIV8 (N=3). Statistical significance was calculated using a matched Friedman test with an uncorrected Dunn’s post hoc test for each line. Significance represented as: 0.1234 (ns), 0.0332 (*), 0.0021 (**), 0.0002 (***), <0.0001 (****).

C. Quantification of p62 Intensity (A.U.) in SMN expression categories determined independent for each line at DIV8 (N=3). Statistical significance was calculated using a matched Friedman test with an uncorrected Dunn’s post hoc test for each line. Significance represented as: 0.1234 (ns), 0.0332 (*), 0.0021 (**), 0.0002 (***), <0.0001 (****).

D. Quantification of CULLIN5 Intensity (A.U.) in SMN expression categories determined independent for each line at DIV8 (N=3). Statistical significance was calculated using a matched Friedman test with an uncorrected Dunn’s post hoc test for each line. Significance represented as: 0.1234 (ns), 0.0332 (*), 0.0021 (**), 0.0002 (***), <0.0001 (****).

**Supp. Figure 3.** Validation of SMA and Isogenic ALS hiPSC-derived MNs

A. Representative images of WT-BJ, Type II (51N), Type I (38D), and Type I (33A) MNs at DIV2 and stained for SMN (orange), ISL1 (cyan), and HOESCHST (gray).

B. Percentage of ISL1+ MNs at DIV8 of dissociated hiPSC-derived MN cultures. WT-BJ N=6; Type II (51N) N=3; Type I (38D) N=3 Type I (33A) N=3.

C. Immunoblot of EB15 protein lysates from the indicated lines probed for SMN and beta-Tubulin (N=3/line).

D. Relative SMN Intensity (A.U.) in ISL1+ MNs at DIV2 and DIV8 of WT and SMA hiPSC-derived MNs to demonstrate the reduction in SMN protein between healthy and diseased states. Statistical comparisons were calculated within each time point using an unpaired Brown-Forsythe and Welch ANOVA test and the Unpaired t with Welch’s correction post-hoc test. DIV2: WT-BJ N=7; Type II (51N) N=4; Type I (38D) N=4; Type I (33A) N=3. DIV8: WT-BJ N=6; Type II (51N) N=3; Type I (38D) N=3 Type I (33A) N=3.

E. Cladogram demonstrating successful introduction of the SOD1_G85S_ mutation (GGC>AGC) and TDP-43_M337V_ mutation (ATG>GTA) WT-BJ *SOD1* and *TARDBP* gene loci.

F. Percentage of ISL1+ MNs at DIV6 of dissociated hiPSC-derived MN cultures. Each point represents a biological replicate. WT-BJ N=11; #16 BJ N=7; #18 BJ N=4; SOD1_G85S_ N=5; TDP-43_M337V_ N=3; WT-1016A N=3; #5 1016A N=3; #6 1016A N=3.

G. Relative # of ISL1+ MNs from DIV2 to DIV10 of hiPSC-derived MN cultures. Statistical comparisons were calculated between DIV2 to DIV10 within each line using a Mixed-Effects Analysis with the Geisser-Greenhouse correction and the Uncorrected Fisher’s LSD post-hoc test. Significance represented by the color of each cell line: 0.1234 (ns), 0.0332 (*), 0.0021 (**), 0.0002 (***), <0.0001 (****).#16 BJ N=7; SOD1_G85S_ N=5; TDP-43_M337V_ N=3.

H. Relative SMN Intensity (A.U.) in ISL1+ MNs from DIV2 to DIV10 of hiPSC-MN cultures. Statistics compare DIV2 to DIV10 within each line using a Mixed-Effects Analysis with the Geisser-Greenhouse correction and the Uncorrected Fisher’s LSD post-hoc test. Significance represented by the color of each cell line: 0.1234 (ns), 0.0332 (*), 0.0021 (**), 0.0002 (***), <0.0001 (****).#16 BJ N=7; SOD1_G85S_ N=5; TDP-43_M337V_ N=3.

I. Relative SMN-Clover Intensity (A.U.) in ISL1+ MNs from DIV2 to DIV10 of hiPSC-MN cultures. Statistics compare DIV2 to DIV10 within each line using a Mixed-Effects Analysis with the Geisser-Greenhouse correction and the Uncorrected Fisher’s LSD post-hoc test. Significance represented by the color of each cell line: 0.1234 (ns), 0.0332 (*), 0.0021 (**), 0.0002 (***), <0.0001 (****).#16 BJ N=7; SOD1_G85S_ N=5; TDP-43_M337V_ N=3.

J. Representative SMN (left) and SMN-Clover (right) Intensity (A.U.) histograms of SOD1_G85S_ SMN-Clover, hiSPC-derived ISL1+ MNs at time points from DIV2 to DIV10.

K. Same as (J) for TDP-43_M337V_.

L. Kaplan-Meyer survival plot of #16 BJ (N=3) and SOD1_G85S_ (N=3) SMN-Clover MNs exposed to no treatment or 1µM NaARS during the 4 day live imaging experiment (N=3). HR: hazard ratio between low- and high-expressors. Error bars: 95% confidence interval. Comparisons between survival curves was done using the Log-rank (Mantel-Cox) test.

M. Same as (L) but comparing #16 BJ (N=3) and TDP-43_M337V_ (N=3) SMN-Clover MNs.

**Supp. Figure 4.** Endogenous SMN predicts motor neuron survival across SMA and ALS

A. Representative images of WT-BJ, Type II (51N), and Type I (38D) CNs at DIV10 and stained for SMN (orange), ISL1 (cyan), and Hoechst (gray).

B. Percentage of BRN2+ CNs at DIV10 of hiPSC-derived CN cultures. WT-BJ N=6; Type II (51N) N=3; Type I (38D) N=3 Type I (33A) N=3.

C. Fraction of BRN2+ WT-BJ, Type II (51N), Type I (38D), and Type I (33A) MNs that survived until DIV14 relative to the number quantified at DIV10. Statistical significance was calculated using an unpaired, nonparametric Kruskall-Wallis test with Uncorrected Dunn’s post hoc test. WT-BJ N=3; Type II (51N) N=3; Type I (38D) N=3.

D. Relative SMN Intensity (A.U.) in BRN2+ MNs at DIV10, DIV12, and DIV14 of WT and SMA hiPSC-derived CNs to demonstrate the reduction in SMN protein between healthy and diseased states. Statistical comparisons were calculated within each time point using an unpaired Brown-Forsythe and Welch ANOVA test and the Unpaired t with Welch’s correction post-hoc test. WT-BJ N=3 Type II (51N) N=3; Type I (38D) N=3; Type I (33A) N=3.

E. Representative SMN Intensity (A.U.) histogram of WT (WT-BJ) and SMA (Type II-51N and Type I-38D) hiSPC-derived BRN2+ CNs demonstrating both heterogeneous expression and increase in SMN abundance at DIV10.

F. Representative SMN Intensity (A.U.) histogram of WT (WT-BJ) and SMA (Type II-51N, Type I-38D) hiSPC-derived BRN2+ CNs at DIV14.

G. Dot plot comparing SMN Intensity (color) and percentage of classified (size) from WT-BJ and SMA BRN2+ CNs classified as high-, mid-, and low-SMN expressors based on WT-BJ SMN expression at DIV2.

H. Percentage of BRN2+ CNs at DIV10 of hiPSC-derived CN cultures. #16 BJ N=9; SOD1_G85S_ N=11; TDP-43_M337V_ N=3.

I. Relative # of BRN2+ CNs from DIV10 to DIV14 of hiPSC-derived CN cultures. Statistical comparisons were calculated between DIV10 to DIV104within each line using a Mixed-Effects Analysis with the Geisser-Greenhouse correction and the Uncorrected Fisher’s LSD post-hoc test. Significance represented by the color of each cell line: 0.1234 (ns), 0.0332 (*), 0.0021 (**), 0.0002 (***), <0.0001 (****). #16 BJ N=3; SOD1_G85S_ N=3; TDP-43_M337V_ N=3.

J. Relative SMN Intensity (A.U.) in BRN2+ CNs at DIV10, DIV12, and DIV14 of WT and ALS hiPSC-derived CNs to demonstrate the reduction in SMN protein between healthy and diseased states. Statistical comparisons were calculated within each time point using an unpaired Brown-Forsythe and Welch ANOVA test and the Unpaired t with Welch’s correction post-hoc test. #16 BJ N=3; SOD1_G85S_ N=3; TDP-43_M337V_ N=3.

K. Relative SMN-Clover Intensity (A.U.) in BRN2+ CNs at DIV10, DIV12, and DIV14 of WT and ALS hiPSC-derived CNs to demonstrate the reduction in SMN protein between healthy and diseased states. Statistical comparisons were calculated within each time point using an unpaired Brown-Forsythe and Welch ANOVA test and the Unpaired t with Welch’s correction post-hoc test. #16 BJ N=3; SOD1_G85S_ N=3; TDP-43_M337V_ N=3.

L. Representative SMN (left) and SMN-Clover (right) Intensity (A.U.) histograms of SOD1_G85S_ hiSPC-derived BRN2+ CNs demonstrating both heterogeneous expression and increase in SMN abundance at DIV10 to DIV14.

M. Same as (L) for TDP-43_M337V_.

N. Kaplan-Meyer survival plot of *SOD1*-G85S SMN-Clover CNs plotting the probability of survival of MNs classified by their SMN-Clover expression level at each time point of the experiment (N=3). HR: hazard ratio between low- and high-expressors. Error bars: 95% confidence interval. Comparisons between survival curves was done using the Log-rank (Mantel-Cox) test. Significance represented as: 0.1234 (ns), 0.0332 (*), 0.0021 (**), 0.0002 (***), <0.0001 (****).

O. Representative Kaplan-Meyer survival plot of *TARDBP*-M337V SMN-Clover CNs (N=3). See (N).

P. Dot plot comparing SMN Intensity (color) and percentage of classified (size) from WT and ALS BRN2+ CNs classified as high-, mid-, and low-SMN expressors based on #16 BJ SMN expression at DIV10.

**Supp. Figure 5.** Dynamic increases in endogenous SMN define a resilient motor neuron state

A. Representative Kaplan-Meyer survival plot of #18 BJ SMN-Clover MNs plotting the probability of survival of MNs classified by whether they increase or decrease their SMN-Clover intensity at each time point of the experiment (N=8). HR: hazard ratio between increasing- and constant-expressors. Error bars: 95% confidence interval. Comparisons between survival curves was done using the Log-rank (Mantel-Cox) test. Significance represented as: 0.1234 (ns), 0.0332 (*), 0.0021 (**), 0.0002 (***), <0.0001 (****).

B. Representative Kaplan-Meyer survival plot of #6 1016A SMN-Clover MNs (N=3). See (A).

C. Representative Kaplan-Meyer survival plot of #5 1016A SMN-Clover MNs (N=3). See (A).

**Supp. Figure 6.** MN excitability is not determined by extrinsic factors

A. Quantification of the percentage of active cells in each cell line in basal and blocking (TTX) treatments (BJ WT N=4; #16 BJ N=6; #6 1016A N=5; Type II ISO (51N) N=4; Type II (51N) N=4; Type I ISO (38D) N=4; Type I (38D) N=4; SOD1_G85S_ N=5; TDP-43_M337V_ N=3). Statistical significance was calculated using a paired parametric on-tailed T-test for each cell line. Significance represented as: 0.1234 (ns), 0.0332 (*), 0.0021 (**), 0.0002 (***), <0.0001 (****).

B. Quantification of the median peak amplitude in basal and activating (4APBIC) treatments (BJ WT N=4; #16 BJ N=6; #6 1016A N=5; Type II ISO (51N) N=4; Type II (51N) N=4; Type I ISO (38D) N=4; Type I (38D) N=4; SOD1_G85S_ N=5; TDP-43_M337V_ N=3). Each violin plot is constructed from the median peak amplitude of all cells and each point, and the standard error of the mean error bars is the average of all cells per experiment. Statistical significance was calculated on the average of each experiment using a paired parametric on-tailed T-test for each cell line. Significance represented as: 0.1234 (ns), 0.0332 (*), 0.0021 (**), 0.0002 (***), <0.0001 (****).

C. Quantification of the # of Syn1+/PSD95+ along MAP2+ neurites from DIV8 #16 BJ (N=3).

D. Representative image and zoom ins of synaptic staining in #16 WT-BJ MNs with the post-synaptic PSD-95 (cyan) and pre-synaptic Synapsin 1 (magenta) overlaid with the dendritic marker MAP2 (gray).

E. Quantification of #16 BJ SMN-Clover MN soma area (pixel^2^) at DIV8-9 (N=6). Statistical significance calculated using a paired RM one-way ANOVA with the Geisser-Greehouse correction and Tukey’s multiple comparisons post hoc test. Significance represented as: 0.1234 (ns), 0.0332 (*), 0.0021 (**), 0.0002 (***), <0.0001 (****).

F. Same as (E) for #6 1016A SMN-Clover MN soma area (pixel^2^) at DIV8-9 (N=5).

G. Same as (E) for SOD1_G85S_ SMN-Clover MN soma area (pixel^2^) at DIV8-9 (N=4).

H. Same as (E) for TDP-43_M337V_ SMN-Clover MN soma area (pixel^2^) at DIV8-9 (N=4).

I. Representative images of MAP2 (red) stained SMA isogenic hiPSC-derived MNs. Scale bar = 100µM).

J. Quantification of neurite length (µM)/cell in SMN isogenic hiPSC-derived MNs (Type II ISO (51N) N=3; Type I ISO (51N) N=3; Type I ISO (38D) N=3; Type I ISO (38D) N=3). Significance represented as: 0.1234 (ns), 0.0332 (*), 0.0021 (**), 0.0002 (***), <0.0001 (****).

K. Schematic representation of activating (4APBIC) and blocking (TTX) treatments performed on #16 BJ SMN-Clover MNs. Continuously treated cells (white) were fixed on DIV6 after 48hours of treatment, and cells allowed to recover post treatment (gray) were fixed after 48hours of recovery and fixed.

L. Quantification of SMN-Clover Intensity (A.U.) in #16 BJ SMN-Clover ISL1/2+ MNs after continuous (white) and recovery post treatment (gray) with either activating or blocking compounds (N=3). Statistical significance was calculated independently for the continuous and recovery post treatment paradigms using as Friedman test with a post hoc uncorrected Dunn’s test. Only significant comparisons shown. Significance represented as: 0.1234 (ns), 0.0332 (*), 0.0021 (**), 0.0002 (***), <0.0001 (****).

**Supp. Figure 7.** Intrinsic SMN-dependent vulnerability is conserved in vivo

Representative images of p4 and p10/12 L4/L5 spinal cord segments. Upper image panels show outlines of spinal cord sections and Hb9:GFP+ (green) MNs. Outlined box represents the MNs of interest in this segment and the below zoomed in region showing the SMN (magenta) staining. MNs are labelled with a dotted outline for clarity. Scale bar = 100µm.

**Supp. Figure 8.** Survival and activity related genes that may protect high-SMN expressors and reduce their activity

A. Confirmatory western blot probed for SMN and Actin to demonstrate that sorted MNs from #16 BJ and #18 BJ show the corresponding amount of untagged SMN and SMN-Clover based upon their category at both Early and Late time points.

B. Heatmap depicting DEGs from high-SMN vs low-SMN-Clover MNs at early (left) and late (right) time points.

C. Complete string diagram DEPs between high- and low-expressing SMN-Clover MNs. Each node represents one DEP and is colored by whether they were upregulated (red) or downregulated (down) in the high-expressers. DEPs were classified using StringDB and associated GO Terms, KEGG pathways, and Reactome pathways.

D. Volcano plots from MS performed from #16 BJ high-SMN vs low-SMN-Clover MNs at early (left) and late (right) time points. SMN and other proteins are highlight with their names.

E. Same as (D) for #18 BJ.

**Supp. Figure 9.** Survival and activity related genes that may protect high-SMN expressors and reduce their activity

Complete string diagram DEPs between high- and low-expressing SMN-Clover MNs. Each node represents one DEP and is colored by whether they were upregulated (red) or downregulated (down) in the high-expressers. DEPs were classified using StringDB and associated GO Terms, KEGG pathways, and Reactome pathways.

## Supplementary Tables

**Supp. Table 1.** Core differentially expressed genes (DEGs) distinguishing high- and low-SMN motor neurons

**Supp. Table 2.** Conserved differentially expressed proteins (DEPs) distinguishing high- and low-SMN motor neurons

**Supp. Table 3.** Pathway analysis of DEPs between high- and low-SMN neurons

## STAR Methods

### Human iPSC Lines and culture

All experiments involving hiPSCs were performed in accordance with the ethical standards of the institutional and national research committees, as well as the 1964 Helsinki Declaration and its later amendments, and approved by the Ethics Commission at the Technische Universität Dresden (SR-EK 80022020, SR-EK-455112023). The healthy BJ siPSC (HVRDi005-A) and 1016A (HVRDi007-A) lines, and the SMA lines (HVRDi015-A, HVRDi017-A, HVRDi016-A), were kindly provided by Lee L. Rubin (Harvard University) through an MTA. Information regarding both the SMA hiPSC lines and the healthy control lines is enclosed in ^19,38,51^. Our laboratory is certified by the Staatsministerium für Energie, Klimaschutz, Umwelt und Landwirtschaft, Freistaat Sachsen at the biosafety/security level S2 required for handling agents and toxins listed by the CDC as falling under dual use research restrictions (54-8452/111/4). iPS cells were cultured in mTeSR Plus Media supplemented with 1% Penstrep. iPS cells were passaged using 5mM EDTA in 1X PBS onto Matrigel coated plates.

### SMN-Clover hiPSC Line Generation by CRISPR

SMN-Clover hiPSC lines were generated as previously described in Grass et al ^96^. Briefly, hiPSCs were dissociated to single cells using accutase and nucleofected with the pTG-Cr-SMN plasmid and the pTG-HR-SMN targeting vectors using the 4-D nucleofector system (AMAXA) and P3 Primary Cell 4D-Nucleofector Kit (Lonza) following the manufacturer’s instructions (program CB-150). Successfully targeted cells were plated to Matrigel-coated dishes with mTeSR plus supplemented with 4µM ROCK inhibitor before selecting for 1 week with 1µg/mL puromycin. To excise the mRuby-T2A-Puromycin selection cassette, cells were again nucleofected with pCAG-Cre:GFP, and 24hr later, GFP^+^:mRuby-cells were enriched by FACSorting. Post-Cre-excised hiPSCs were plated at clonal densities and clones picked and screened for successful *SMN2* loci targeting using the targeted *SMN1/2* and untargeted *SMN1/2* primer pairs. See the *Reagents and Tools Table* for relevant plasmids and sequences.

### Isogenic ALS hiPSC line generation by CRISPR

#### Isogenic ALS SOD1_G85S_

The isogenic SOD1_G85S_ hiPSC line was generated from the parental BJ WT line by CRISPR/Cas9-mediated homology-directed repair. Approximately 1 × 10^6 hiPSCs were nucleofected using a 4D-Nucleofector™ System (Lonza) with Alt-R™ S.p. HiFi Cas9 Nuclease V3, SOD1-targeting Alt-R™ CRISPR-Cas9 crRNA, ATTO 550-labeled Alt-R™ CRISPR-Cas9 tracrRNA, and a single-stranded Ultramer™ DNA oligonucleotide donor carrying the desired G85S substitution (Integrated DNA Technologies, IDT). Twenty-four hours after nucleofection, ATTO 550-positive cells were enriched by fluorescence-activated cell sorting (FACS) and plated at clonal density on 10-cm culture dishes. Individual colonies were manually picked once they had reached sufficient size and expanded. Genomic DNA was isolated from individual clones, and the SOD1 genomic region encompassing the targeted site was amplified by PCR and analyzed by Sanger sequencing (Microsynth). Correctly targeted clones carrying the G85S substitution were subsequently expanded for further characterization and experimental use. <u>Isogenic TDP-43_M337V_:</u> The Isogenic TDP-43_M337V_ hiPSC line was generated from the healthy #16 BJ SMN-Clover hiPSC line using a two-vector CRISPR-cas9 approach described in Grass et al. ^97^. Briefly, the px458 plasmid expressed both Cas9 and a single-guide RNA (sgRNA) targeted to the specific point mutation locus in exon5 of the *TARDBP* gene. A second plasmid, a modified version of the HR120-PA1 vector, served as a donor template for homology-directed repair while containing the desired point mutation, 420bp homology arms, and a dual-selection cassette (mRuby-T2A-puromycin) flanked by LoxP sites. hiPSCs were nucleofected as previously described with these two vectors and clones were then selected as described for the hiPSC SMN-Clover lines.

### Motor Neuron Differentiation and Culture

hiPSC-derived motor neurons were derived and cultured following the same embryoid body (EB) based protocol described in ^51^ and represented in Figure S1B. Embryoid bodies (EBs) were generated in 10cm ultra-low attachment dishes from accutase, dissociated single cell iPS suspensions. EBs were cultured in mTeSR Plus supplemented with 10µM Y-27632 and 10ng/mL FGF-2 for 3 days. On D0, EBs were gravity precipitated to remove death remaining from EB formation and switched to Neural Induction Media (NIM; 50% Advanced DMEM/F-12, 50% Neurobasal Medium, 1% Penstrep, 1% GlutaMAX (Thermo Scientific, 35050087), 0.1 mM *β*-Mercaptoethanol, 0.5X B-27, 0.5X N2, 0.16% D(+)-glucose and 20 µM ascorbic acid) to begin MN differentiation. Neural induction was performed using dual-SMAD inhibition by supplementing NIM with 10µM SB431542 and 100 nM LDN 193189 between day 0 and day 5. Caudal and ventral patterning is achieved by adding 1µM retinoic acid and 1 µM Sonic Hedgehog Signaling Agonist starting on day 2. To support neural progenitor growth, 10 ng/ml BDNF was supplemented from day 7. From day 9 to day 11, 2.5 µM DAPT was added, followed by 10 ng/ml GDNF and 2 µM cytosine arabinoside (Ara-C) on day 11 to specify motor neuron fate and remove proliferating cells. On day 15, EBs were dissociated to single cells using the Worthington Papain Dissociation System (Worthington, 9035-81-1, 9048-46-8, 9003-98-9, 9001-73-4) and plated on 50µg/mL, 1X Borate Buffer and 3µg/mL Laminin coated plates. MNs were plated at a density of 80,000cells/well (p96), 30,000cells/well (384well plate) or 4.5-5million cells/well (p6). Cells were cultured in human Motor Neuron Media (hMN) comprised of Neurobasal Medium, 1% Penstrep, 1% GlutaMAX, 1% Non-Essential Amino Acids, 25µM ß-Mercaptoethanol, 0.5X B-27, 0.5X N-2, 0.16% D(+)-glucose and 20 µM ascorbic acid. Media was supplemented with 10ng/mL BDNF, 10ng/mL GDNF, and 1-2µM Ara-C (depending on Glial cell content) and changed every 3-4 days unless otherwise specified.

### Cortical Neuron Differentiation and Culture

Human cortical neurons were generated following protocols from ^38,57^ that initiate 2D neuronal cell differentiation through lentiviral mediated, neurogenin-2 (Ngn2) overexpression. iPSCs were dissociated to single cells using accutase and plated to Matrigel-treated multi-well cell culture plates at 5,000-30,000 cells per well, in a line dependent manner, in mTeSR Plus supplemented with 1% PenStrep and 10µM Y-27632. The following day, cells were transduced with two lentiviruses, one carrying a constitutively expressed a tetracycline-controlled transcription activator (rtTA) and another carrying an Ngn2 and puromycin resistance cassette that were both under tetracycline control (tetO promoter). Co-infection thereby enabled doxycycline-inducible Ngn2 expression and puromycin selection. Continuous 0.2µg/mL doxycycline treatment was started alongside transduction to induce tetracycline operator (tetO) promoter activity. Two days later, the virus was removed, and medium switched to cortical neuron media (CNM) (Neurobasal Medium, 1% Pentrep, 1% GlutaMAX, 1X non-essential amino acids, 0.5X N2, 0.5X B27 without vitamin A, 20 µM ascorbic acid and 0.1 mM ß-Mercaptoethanol) supplemented with 10 ng/ml BDNF and 10 ng/ml GDNF. Beginning on day 3, puromycin selection was started (0.5 µg/ml), and 0.2 µg/ml laminin was supplemented to support plate adhesion of neuronal precursors. To remove dead cells following puromycin treatment, a full media change was performed on day 4. Starting on day 6, 2 µM cytosine arabinoside (Ara-C) was supplemented, and 50% of media was changed every two days until the end of the experiment.

### Live Longitudinal Imaging, Quantification, and Analysis

iPSC derived-MNs or -CNs plated in 96-well plates or 384-well plates were allowed to mature to either DIV4 post dissociation or DIV10 post lentiviral transduction to achieve a SMN-Clover distribution with 3-4x range of expression. Neurons were stained for 2 hours at 37°C with 125nM SiR-DNA, the far-red nuclear dye. Following staining, were washed with the respective neuronal medium before adding non-phenol red containing culture media as described for each cell type above. In the case of a longitudinal treatment compounds were added to this media. The plate was acclimated for 1-2hours in the Perkin Elmer (now Revity) Operetta CLS prevent deformation artifacts in the videos. The cells were imaged under at 5% CO2 and 37°C conditions for 4 days 1 frame/4- (MNs) or 2-hours (CNs), to account for differences in cell movement. The cells were imaged using non-confocal mode, a 40X (NA 1.1) Water objective, and the preset filters for Alexa 647 and Clover. Following imaging cells were fixed in 4% PFA and stained for additional markers of interest to confirm experimental reliability (i.e ISLET1, BRN2, MAP2, SMN, etc.).

Videos were analyzed using Perkin Elmer’s proprietary image analysis software Columbus. A high-throughput Columbus script segmented single neuronal nuclei and cytoplasm and measured both intensity and morphological properties overtime by tracking each cell’s movement. Clumped nuclei were removed at this stage. At each time point both morphological and intensity measurements were used to classify cells as living or dead (see *Data Availability* for access to analysis details on Github). All composite and time resolved, single-cell information was output by Columbus for further quality control and analysis in a custom R-markdown notebook (see Data Availability). Briefly this notebook assigns user-defined metadata to the experiment (cell lines, treatments, time points, controls, etc.), removes objects with insufficient information (dependent on the cell type), makes a final live/dead determination for each cell at each time point, subtracts background intensity from measured channels, classifies cells by their static and dynamic SMN-Clover intensity (see *SMN Classification Analysis* section for details), performs Kaplan Meyer survival analysis using the Rpackage *survival*^98^, and outputs spreadsheets that are then plotted and statistically analyzed using Graphpad Prism (see *Statistics* and relevant figure legends).

### SMN Classification Analysis

To determine if a cell was a High, Mid, or Low SMN-Clover expressor in live, *longitudinal imagine experiments*, the average corrected SMN-Clover intensities over all time points of a cell and arbitrarily determined cutoffs were determined based upon the SMN-Clover distribution of each line and treatment. For MNs cutoffs of -0.5sd and +1.25sd of the mean were used, and - 0.5sd and +0.75sd for CNS were used. To identify if the SMN-Clover intensity of a cell remains constant or changes overtime, a linear regression over time of SMN-Clover intensity was used and if it was significantly positive the cell was labelled as “increasing,” significantly negative “decreasing,” and non-significant or undefined “constant.” In the case of *fixation experiments*, classified intensities were background corrected, cutoffs remained the same for each cell type, and classification was done at the indicated time point. However, to perform the analysis present in Figure 2D, 2G, S4G, and S4P the cutoffs were determined on reference data dependent on the desired comparison. To compare the expression of SMN in different classifications between cell lines, cutoffs were determined and applied for each cell line independently. To compare the percentage of cells that fall into a particular SMN classification, cutoffs were determined using the SMN WT BJ or SMN-Clover #16 BJ distribution and then applied to the SMA or ALS distributions.

### Calcium imagining and analysis

iPSC derived-MNs in 96- or 384-well plates were stained with 125nM SiR-DNA at 125nM for 2 hours at 37°C. Cells were subsequently stained for 45-minutes at 37°C with the calcium indicator dye, Cal590 (AAT Bioquest, 20510), which fluoresces when Ca^2+^ ions bind the molecule allowing for visualization of calcium transients within single neurons. Cells were then washed with motor neuron media to remove excess dye and dead cells. Cells were treated for 30minutes before imaging. Treatments applied to individual sets of cells in hMN media minus phenol red in all experiments: (1) basal, standard media, (2) activating, 4-aminopyridine (4-AP, 200µM) and Bicuculine (BIC, 60µM), to confirm cells respond to activity modulation, and (3) blocking, tetrodotoxin (TTX, 1µM) to confirm the calcium influxes result from action-potential like phenomena ^99,100^. Cells were imaged live in the Yokagawa CV7000 under 37°C and 5% C0_2_. 2-4 fields/well were imaged with 3-4 wells used for each treatment. The imaging program operated as follows: (1) calcium influxes as shown by the Cal590 for 1.5minutes at a rate of 5Hz, (2) z-stack of SMN-Clover, (3) SiR-DNA nuclear staining, and (4) a bright field image. A KNIME-based analysis was used to measure calcium transients (hereafter referred to as spikes) in hiPSC-derived MNs. The generalized procedure for calcium trace analysis and spike detection is as follows: (1) correct for generalized noise in the data by normalizing each timepoint by the median ratio of a defined time window’s median intensity and that timepoint’s raw intensity for all cells, (2) transform the data to remove underlying trends by subtracting the 10^th^ quantile of intensity values in a defined window size, (3) transform the data by subtracting the minimum intensity value of a selected window size from each time point, and (4) normalize the data by dividing the sum of the trace produced by the previous two transformations (Steps 2 and 3), and (5) smooth the data using moving average over a defined window size. This results in an approximation of ΔF/F_0_, the most common measure of calcium fluorescence in literature interpreted as the change in fluorescence over a defined baseline. Spikes are detected on the smooth data using the “Find Peaks” function from the *pracma* package ^101^ and multiple parameters were reported for each spike per cell. Where necessary, the KNIME classified MNs by their SMN-Clover expression where the top 10% of cells were classified as high, the lowest 40% as low, and the middle 50% as mid as previously reported ^38^. All composite calculations and statistics were performed in PRISM Graphpad. The KNIME pipeline is available on Github at the link provided in the *Data Availability* section.

### Lentiviral generation and transduction

HEK293T cells were cultured in DMEM containing 10% heat-inactivated FBS, 1% Penstrep and 1% GlutaMAX on cell-culture treated dishes. Cells were passaged using TrypleE. When generating 2^nd^ generation lentivirus, HEK293T cells were passaged to cell-culture treated dishes coated with 0.1% gelatin. HEK293T cells were transfected with lentiviral packaging plasmids (psPAX2 and pMD.G) and the construct containing plasmid using lipfectamine 2000 in Optimem After 6hours of incubation, media was changed fresh DMEM (as previously described) and collected three times over the next 3 days. Media was filtered each time using 45micron filters, mixed with lentiviral concentrator Lenti-X and stored at 4°C with agitation. The three collections were spun at 4°C and 2500rpm for 45 minutes, and the resulting pellet was resuspended in either hMN media or mTeSR and flash frozen on dry ice before storing at -80°C. Lentiviral plasmids used in this study can be found in the *Reagents and Tools Table*.

Embryoid bodies (EBs) were transduced with lentivirus at D9 of the motor neuron differentiation protocol to optimize for the infection of newly born neurons. Transduction was done in a minimal media amount (1mL) in 6-well plastic plates using a 1:100 dilution. After 24 hours on D10 1mL of media with the D9 media formulation as shown in Figure S1B was added reducing the lentiviral dilution to 1:200. After 24 additional hours with lentivirus on D11, a full media change (FMC) to the D11 formulation was performed. At D15, EBs were fixed in 4% PFA diluted in 1X PBS overnight at 4°C before a 24hour wash in 1X PBS.

### SMNΔ7 Mouse Model and Spinal Cord Extraction

All animal studies were approved by the ethics committee of the Technische Universität Dresden and the Landesdirektion Dresden (approval numbers: TVV 4/2022; 25–5131/542/6). All relevant European Union regulations, German laws (Tierschutzgesetz) and the NIH Guide for the Care and Use of Laboratory Animals (National Academies Press, 2011) were strictly followed for all animal work. Heterozygous SMNΔ7 Hb9:GFP mice (Smn+/−;hSMN2+/+; hSMNΔ7+/+;Hb9:GFP+) on an FVB background were kept under standard conditions and bred to harvest healthy littermate controls (Smn+/−;hSMN2+/+; hSMNΔ7+/+;Hb9:GFP+) and SMA (Smn−/−;hSMN2+/+; hSMNΔ7+/+;Hb9:GFP+) mice at P1, P4, P7/8, and P10/12. Spinal cord extraction and segmentation was performed as described in ^13,102^. Briefly, dorsal roots and the spinal cord were isolated under a dissection microscope from the intact back of each mouse. By counting dorsal roots from the sacral region and observing the shape of the cord the location of L1, L2, L4, and L5 segments was identified and confirmed during spinal cord imaging of the Hb9:GFP signal. Each cut lumbar segment was placed into a separate microcentrifuge tube and cryopreserved and sectioned as described above. And immersed in 30% sucrose overnight at 4°C to cryopreserve the tissue. Once tissue was fully saturated and sunken to the bottom it was embedded in OCT compound and frozen at -80 °C for at least 24 h.

### Cryosectioning

EBs and spinal cords were embedded in PolyFreeze Tissue Freezing Medium compound and frozen at -80°C. Cryosections were made with a Liecca CM 3050S at a 12µm (EBs) or 30µm (Spinal Cord) thickness. Sections were transferred to Super Frost Plus positive charged Microscope slides (Epredia), heat-fixed on a benchtop heat plate at 37 °C for 45 min to ensure adhesion before storage at -20 °C or they were immediately processed for immunostaining.

### Immunofluorescence staining and imaging

#### Plated Cells

Cells were fixed in 4% Methanol Free PFA created from mixing hMN Media with 1X Dulbecos PBS -Ca/-Mg PBS diluted 16% PFA for 12-15minutes. Following fixation cells were washed 3 times with 1X Dulbecos PBS -Ca/-Mg. Cells were permeabilized and blocked by a 20minute, room temperature incubation in 0.25% Triton and 1.0% Normal Goat Serum in 1X Dulbecos PBS -Ca/-Mg (referred to as P&B Buffer). Primary antibodies were diluted in P&B Buffer and incubated on cells at 4°C overnight with orbital shaking. The next day, cells were washed 3x with 1X Dulbecos PBS -Ca/-Mg for 10minutes each at room temperature with orbital shaking. Following the washes, secondary antibody solutions were made with all secondary antibodies diluted to 2.5µg/mL and incubated for 2 hours at room temperature. 1X Dulbecos PBS -Ca/-Mg washes were repeated, with the second wash containing 1:10,000 Hoechst 33342 (Invitrogen; H3570) to stain nuclei, and after the last wash the cells were stored in 1X Dulbecos PBS -Ca/-Mg before imaging in the Perkin Elmer Operetta CLS. Stained cells were imaged in the Perkin Elmer Operetta CLS. Imaging was performed using the following settings: non-confocal mode, 2pixel binning, 2 peak autofocus, and the 40X (NA 1.1) or 20X (NA 0.4) water objectives. The DAPI, Alexa 488, Alexa 546, and Alexa 647 preset filters were used as necessary with exposure time and LED power standardized for each protein assayed. <u>Slides</u>: EB and spinal cords were stained the same as fixed cells with the following modifications. Before staining, slides were heat fixed for an additional 45minutes at 37°C and immersed in 1X PBS for ∼15 min to remove OCT. An initial permeabilization step (30minutes in 1X PBS and 0.1% Triton in a humidity chamber at room temperature) for spinal cords was added before permeabilization and blocking to account for its additional thickness compared to EBs. The P&B solution was made with 1% Triton X-100, 5% normal goat serum in 1X PBS was added and slides were incubated for 1.5 h in a humidity chamber. Slides were then stained with 1:1000 Hoechst 33342 (Invitrogen; H3570) and mounted with Fluoromount (Invitrogen; 00-4958-02) before sealing with clear nail polish and drying overnight. Imaging was performed on a Nikon Spinning Disk system on a Ti2-E inverse stand with a Yokogawa CSU-W1 Dual-Disk scan head and a Hamamatsu Orca Fusion camera (Hamamatsu), using a 20x/0.8 CFI Plan Apochromat Lambda D objective (Spinal Cords) and 10X/0.45 CFI Plan-Aprochromat Lambda D (EBs). A z-step of 3.5 μm and 2.5µm was used for spinal cords and embryoid bodies, respectively. Fluorophores were acquired using 405 nm, 488 nm, 546 nm, and 647 nm lasers with exposure time standardized for each protein assayed. See *Reagents and Tools Table* for list of antibodies and used concentrations

### Image Analysis

#### Plated Cells

Images taken in the Operetta CLS were transferred to the Perkin Elmer Columbus server system (v.2.6.0) for analysis. The system allows one to create image analysis scripts from pre-programed “blocks” that performed specific functions. Scripts were structured to find nuclei based upon DNA staining, find cytoplasmic regions based upon various markers, find spots within the nuclei or cytoplasm, calculate intensity of various markers, and determine if cells are living or dead (all dead cells were excluded from the analysis). <u>EBs</u>: Acquired nd2 images were processed and analyzed using semi-automated FIJI macros. Images were first maximum projected and those maximum projections were used to quantify the total number of cells (identified by Hoechst staining), positively transduced cells (identified by mCherry+ signal), and ISL1+ MNs based on immunostaining. This was accomplished by first prompting the user to crop the image to the relevant area. The Hoechst staining was processed first using the Unsharp Mask command (radius = 5, mask = 0.60) and a user defined Threshold was set to identify living nuclei. This threshold was turned into a binary and a Watershed command was run to segment individual nuclei. Both the mCherry and ISL1 images were processed similarly but only used a user defined threshold to determine positive cells. The Image Calculator command was used to identify mCherry+;Hoechst+ (mCherry+ cells), ISL1+;Hoechst+ (ISL1+ MNs), mCherry+;ISL1+;Hoechst+ (mCherry+;ISL1+ MNs). A particle analysis was performed on each channel to calculate the total number of cells from all populations. The resulting spreadsheets were then compiled and further quantified using a custom R script. Code for FIJI macros is available on Github at the link provided in the Data Availability section. <u>Spinal Cords</u>: Spinal cord section images were processed using NIS-Elements (Nikon) and ARIVIS Vision 4D software (Zeiss). Acquired z-stacks were converted to maximum intensity projections in NIS-Elements, with blurred images removed by manual inspection. Using the “Draw Object” tool in ARIVIS single motor neurons were identified by high expression of Hb9::GFP or Hb9 immunostaining in GFP negative tissue. Motor neuron size and position were used to determine their columnar identity (LMC or MMC). Single cell immunofluorescence intensities for each imaged channel, morphological features, and identity tags features and cellular morphology features were exported for each section. These spreadsheets were compiled and processed using an automated R-script for each mouse and each time point allowing for the tabulation of motor neuron counts and SMN intensity which was corrected using a single background measurement obtained from a P4 CTRL mouse stained only with secondary antibodies. Code is available on github at the link provided in the Data Availability section.

### Western Blot

Cells or EB pellets were lysed in RIPA + 2% SDS buffer supplemented with HALT Protease Inhibitor. Protein concentrations of lysates were measured using the Pierce BCA Protein Assay Kit. Western Blots were performed using BioRad AnykD Criterion TGX Precast Midi Protein Gels with appropriate numbers of wells. Gels were run in 1X Tris-Glycine SDS Running Buffer using 60V for ∼30min and 100-110V for ∼2hours. After running, the gel was removed from the gel casing and equilibrated in 1X Tris/Glycine Transfer Buffer for ∼5min. The equilibrated gel was assembled into the pre-made BioRad TransBlot Turbo-Transfer Pack with 0.2µm PVDF membrane, placed in the BioRad Trans-Blot Turbo System, and the proteins transferred to the membrane using either the High MW (2.5A 25V 10min) or Standard SD (25V 1.0A 30min) pre-set programs. Once transferred, the membrane was quickly rinsed with diH_2_O and stained with Ponceau to confirm even loading. Membranes were then washed to remove the stain and blocked with 5% Nonfat Dried Milk Powder in 1X TBST for 1 hour at room temperature and agitation. The desired antibody was then added and incubated at 4°C overnight; the next day the membrane was washed 3x with 1X TBST for 10min each, probed for the secondary antibody diluted in 5% Milk at room temperature with shaking, and washed again. The signal was developed using western blot substrate (SupraSignal West PICO PLUS Chemiluminescent Substrate), X-ray Films, and the Cawomat 2000 IR X-ray developer. Developed films were scanned using the Epson Perfection C470 in 8-bit Black and White and 300dpi settings before quantification using the BioRad Image Lab Software (v6.0.0 Build 26).

### qPCR

Total RNA was extracted using the TRIZOL Reagent Protocol and treated with DNase I. cDNA was synthesized utilizing the High-Capacity cDNA Reverse Transcription Kit with the following program: 10min at 25°C, 2hrs at 37°C, and 5min at 85°C. Quantitative Real-Time PCR was performed in 96- or 384well plates. Each reaction was 10µL comprised of the GoTaq qPCR Master Mix, 1-2.5ng of cDNA, and 100nM of forward and reverse primers. The reaction was processed in the CFX Connect Real-Time PCR Detection System or the QuantStudio Real-Time PCR System. C*_τ_* values were normalized to 18S rRNA expression. See *Reagents and Tools Table* for all qPCR primers.

### FACS of SMN-Clover iPSC-derived MNs

WT-BJ and WT-BJ SMN-Clover hiPSC-MNs were collected from 6-well cultures on DIV4 and DIV12 and sorted into high-, mid-, and low-SMN expressing SMN-Clover populations in the BD Biosciences FUSION FACS ARIA III. Briefly, on each sort day, motor neurons were stained with calcein red-orange AM, dissociated with 10-15minutes incubation of papain at 37°C, detached with chilled detachment media (hMN media supplement with 20µg/mL of BDNF/GDNF and Worthington DNAse), and transferred to a 15mL falcon tube. Detached cells were spun at 4°C for 3minutes at 12000rpm. The resulting cell pellet was resuspended with minimal agitation with detachment media and filtered using 5mL round bottom tubes with 35µm cell strainer caps. 50,000-100,000 living cells were sorted and collected at 4°C into 1.5mL SafeSeal Rnase free microfuge tubes filled with collection media (hMN media supplement with 20µg/mL of BDNF/GDNF and DNAse). Collection media was supplemented with 2.5U Rec Rnasin if samples were for bulk-RNA sequencing or 100X HALT Protease Inhibitor if samples were for bulk-mass spectrometry. After sorting cells were pelleted by spinning at 4°C and 5000rpm for 5 minutes. Living cells were selected for based on their size as determined by Side-scattering and forward-scattering measurements. Viability was determined using calcein red-orange intensity using the PE filter. SMN-Clover expression was measured using the FITC filter and expression level aimed to produce roughly 30% of the population into high-, mid-, or low-SMN-Clover expression. The minimum fluorescence of SMN-Clover was determined by measurements of control WT BJ hiPSC-MNs.

### Bulk-RNA sequencing (SmartSeq 2) and analysis

Pelleted, FACSorted MNs were lysed in 0.5-1mL of TRIZOL Reagent and total RNA immediately extracted as described in *qPCR.* RNA sample quality was assessed using the Qbit Biozanalyzer, and only samples with RIN (RNA integrity) ≥ 6.4 were used in the bulk-RNA sequencing procedure. Total RNA was processed by the Dresden Concept Genome Center as follows. For the cDNA synthesis, up to 12 µl RNA was evaporated in a vacuum centrifuge for 10-45 minutes at room temperature to reduce the volume to ∼2 µl. The RNA was subsequently denatured for 3 minutes at 72 °C in the presence of 2.4 mM dNTP (Invitrogen), 240 nM dT-primer^*1^ and 4 U Rnase Inhibitor (NEB). The reverse transcription and addition of the template switch oligo was performed at 42°C for 90 min after filling up to 10 µl with RT buffer mix for a final concentration of 1x superscript II buffer (Invitrogen), 1 M betaine, 5 mM DTT, 6 mM MgCl2, 1 µM TSO-primer^*2^, 9 U Rnase Inhibitor and 90 U Superscript II, followed by a heat inactivation of the reverse transcriptase (70 °C for 15 min). With 10% of the material, a qPCR on full-length cDNA was performed with universal primers^*3^ to determine the optimal number of cycles to avoid under- or overamplification of the samples (for 22 cycles with the below mentioned protocol). The single stranded cDNA was amplified using 1x Kapa HiFi HotStart Readymix (Roche) containing 0.1 µM UP-primer^*3^ under following cycling conditions: initial denaturation at 98°C for 3 min, 10-16 cycles according to qPCR results [98°C 20 sec, 67°C 15 sec, 72°C 6 min] and final elongation at 72°C for 5 min. The amplified cDNA was purified using 0.6x volume of hydrophobic Sera-Mag SpeedBeads (GE Healthcare), rebuffered in a buffer consisting of 10 mM Tris, 20 mM EDTA, 18.5 % (w/v) PEG 8000 and 2 M sodium chloride solution. The cDNA is eluted in 12 ul nuclease free water and quality and concentration was determined with the Fragment Analyzer (Advanced Analytical). For library preparation, 2 µl of cDNA was tagmented at 55 °C for 15 min in a total volume of 4 µl containing 1x Tagment DNA Buffer and 0.8 µl Tagment DNA Enzyme (from the Illumina DNA Prep – Tagmentation Kit, Illumina). The reaction was stopped by adding 1 µl 0.1% SDS and incubation for 15 min at 37 °C. After removing the supernatant, the samples were resuspended in 1x concentrated KAPA HiFi HotStart Ready Mix and 0.3 µM dual indexing primers and PCR run under following conditions: (72°C 3 min, 98°C 30 sec, 12 cycles [98°C 10 sec, 63°C 20 sec, 72°C 1 min], 72°C 5 min). After PCR, libraries were purified with 0.9x volume of Sample Purification Beads (Illumina), followed by a size selection with 0.6x (right side) and 0.9x (left side) volume of Sample Purification Beads, and a final 0.9x purification to get a fragment size distribution between 200-700 bp. The libraries were quantified with the Fragment Analyzer, and sequenced with a Novaseq 6000 system (Illumina) on a S4 flowcell in 100 bp paired-end XP mode, aiming at minimum sequencing depth of 40 mio reads per library.

^*1^ dT-primer: C6-aminolinker-AAGCAGTGGTATCAACGCAGAGTCGAC TTTTTTTTTTTTTTTTTTTTTTTTTTTTTTVN, where N represents a random base and V any base beside thymidine;

^*2^TSO-primer: AAGCAGTGGTATCAACGCAGAGTACATrGrGrG, where rG stands for ribo-guanosine;

*^3^ UP-primer: AAGCAGTGGTATCAACGCAGAGT

Bulk RNA-sequencing results were processed by Dr. Nat. Fabian Rost in the following manner. FASTQ files were processed using the nf-core/rnaseq pipeline (v3.14.0) ^103^, executed with Nextflow (v23.10.1) ^104^, and leveraging reproducible software environments provided by Bioconda ^105^ and BioContainers^4^ ^106^. The pipeline was run with the following command: nextflow run nf-core/rnaseq -r 3.14.0 -resume -params-file params.yml -c hpc_slurm_cbg.config –outdir output Key parameters included the use of the *Homo sapiens* GRCh38 reference genome (Ensembl release 104; primary assembly FASTA and GTF files) and additional options for adapter trimming (--nextseq 20 –length 15), STAR alignment (--outFilterMismatchNmax 999 – outFilterMismatchNoverLmax 0.1 –alignMatesGapMax 200000 –chimSegmentMin 20 – twopassMode Basic –alignIntronMin 20 –alignIntronMax 200000), and Salmon quantification (--seqBias –gcBias –posBias). Quality control metrics were evaluated using MultiQC. For downstream analysis, Salmon quantification files were imported into R (v4.3.3) using the tximport package (v1.28.0) ^107^. Variance-stabilizing transformation (VST) was applied prior to principal component analysis (PCA). Differential expression analysis was performed using DESeq2 (v1.40.2) ^108^ with the design formula ∼ age_SMN1.sort + batch + cell_line. DEGs were identified as those with a significant adjusted p-value (< 0.05). For directional interpretation, genes were retained when their fold-change direction did not reverse between developmental stages.

### Proteomics mass spectrometry and analysis

Pelleted, FACSorted MNs or washed with chilled 1X PBS and re-pelleted before flash-freezing on dry-ice. Samples were processed and analyzed by Sofia Vrettou and the University of Cologne CECAD Proteomics Facility. Frozen cell pellets were lysed in 50µL SP3 lysis buffer (5% sodium dodecyl sulfate in 1X PBS). Chromatin was degraded by sonication. Samples were reduced with dithiothreitol (DDT) at a final concentration of 5mM, vortexed and incubated at 55°C for 30minutes. Samples were alkylated with 40mM Chloroacetamide (CAA), vortexed, and incubated at room temperature for 30minutes. Samples were frozen at -20°C before processing by the Proteomics Core Facility. Proteins were digested and beads were removed before cleaning by SDB stage tipping ^109^ instead of the described bead-based clean-up. The digested proteins were separated by reversed phase chromatography on the nano-UHPLC system. As the peptides eluted they were ionized, detected, and analyzed on an Orbitrap Exploris 480 (ThermoFisher Scientific, granted by the German Research Foundation under INST 1856/71-1 FUGG) mass spectrometer equipped with a FAIMS pro differential ion mobility device that was coupled to an UltiMate 3000 (ThermoFisher Scientific). A total proteome data-independent acquisition was performed where the whole mass range is scanned in broad windows and all peptides presented are simultaneously fragmented. Fragments were linked pack to peptides with the help of a pre-build spectral library. Proteins were considered detected if ≥1 unique peptide was found in ¾ replicates (#16 BJ) or 2/3 replicates (#18 BJ). Downstream analysis was carried out in PERSEUS v1.5.1.5. Sample quality was assessed by determining whether log normalized (log_2_x) detected values approached a normal distribution. No samples were excluded based upon this assessment, and missing values were imputed utilizing this distribution. Principle component analysis (PCA) was used to identify if samples cluster according to cell line, time, or SMN category differences. To compare differentially expressed proteins (DEPs) between high- and low-expressers two-sample t-tests were applied for each cell line and time point independently. Each cell line was then processed individually by splitting the peptide count matrix. DEPs were considered significantly differentially expressed if the -logFDR ≥ 1.3 and absolute |difference (log2intensity)| > 0.5 (i.e. greater than 1.4 fold change). The q value was also calculated and used to assess the confidence of significantly different proteins. <u>StringDB:</u> StringDB analysis was carried out using the Protiens with Values/Ranks – Functional Enrichment Analysis. All DEGs with their Log2FoldChanges were uploaded and network was built with the following settings selected: Nework type: full STRING network, meaning of network edges – confidence, active interaction sources – all, minimum required interaction score – medium confidence (0.400), max number of interactors to show – 1st and 2nd shell none. All other settings were changed based on the desired visualization. A clustering analysis was performed using the built in “MCL clustering” method with a 1.7 inflation parameter. The 49 found clusters were manually reviewed for relevant ontological terms from multiple embedded sources resulting in 38 reported clusters. In some cases, clusters had no assigned function, and a name was assigned based on the general function of each protein, and in the case of Spinal Neuron Differentiation these genes were taken from the clusters indicated in Supplemental Table 3 as these genes are known for their role in motor neuron differentiation. All clusters that did not have a determinable ontological term were deemed Miscellaneous, and all disconnected nodes (119) were excluded from visualizations but can be found in Supplemental Table 3. <u>Conserved DEGs:</u> Proteins significantly altered in the high- versus low-SMN comparison at both early and late time points were defined as temporally overlapping DEPs. For directional interpretation, proteins were retained when their fold-change direction was concordant between clones wherever detected and did not reverse between developmental stages within an individual clone. Proteins identified in only one clone were retained when their direction was consistent across the available comparisons. Temporally overlapping proteins with opposing regulation between clones or developmental stages were reported but excluded from pathway-level interpretation.

### Statistics

Data analysis was performed using specific R packages and Prism Graphpad (v10, GraphPad Software, Inc). The Reagents and Tools Table lists key packages and software used for statistical analysis. The data/graphs are presented as mean with the standard error of the mean (SEM) where appropriate. Kaplan-Meier survival curves are presented as the probability of survival against age (h) and 95% confidence intervals. Biological replicates (N) are termed as independent batches of differentiated hiPSCs, passages of hiPSCs, or individual mice. Where appropriate, the number of cells or EBs quantified is signified with n. Statistical tests and post hoc tests are indicated in each Figure legend. The level of conventional statistical significance was set at p<0.05 and displayed as p<0.1234 (ns), p<0.0332 (*), p<0.0021 (**), p<0.0002 (***), p<0.0001 (****).

## Reagents and Tools Table

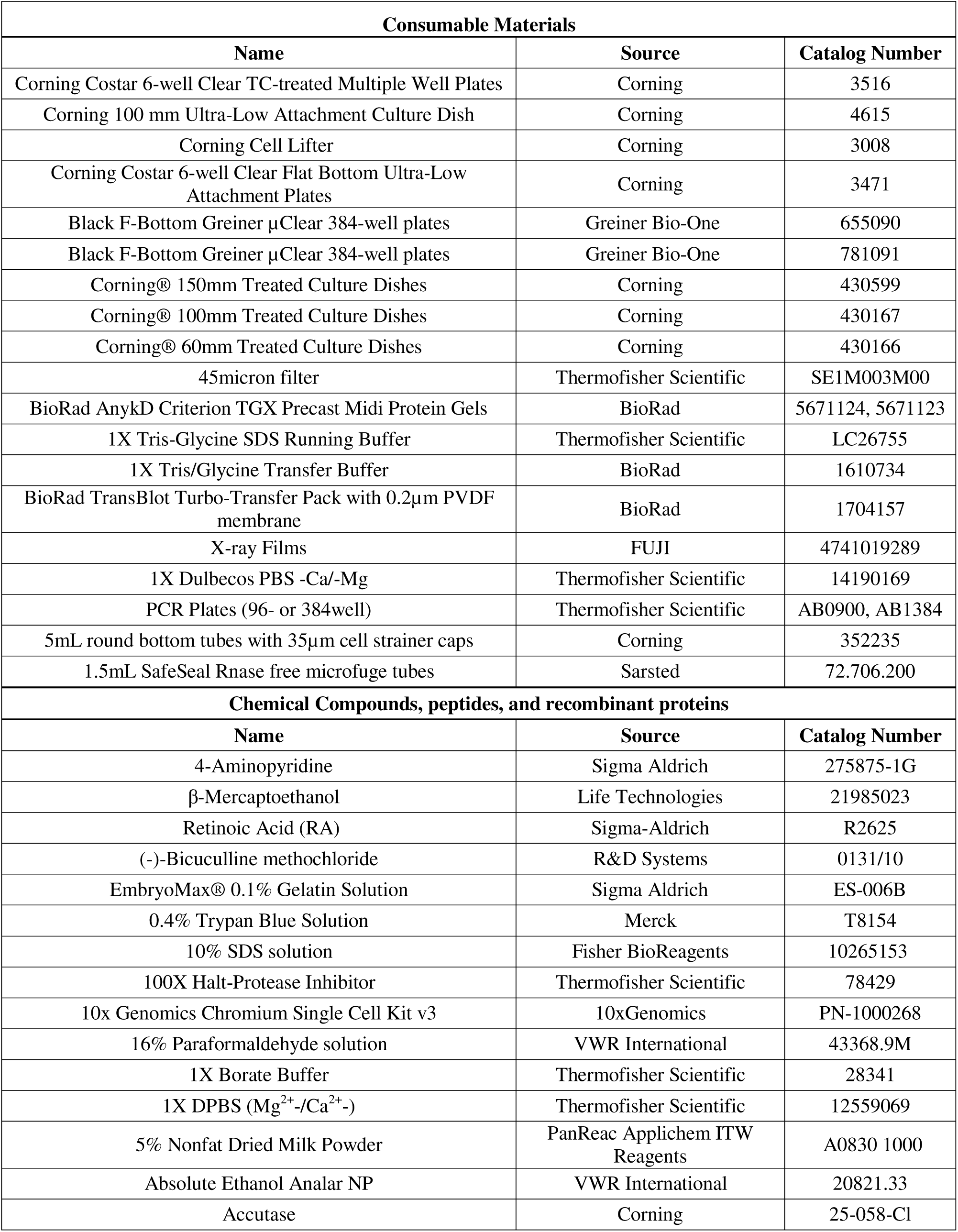
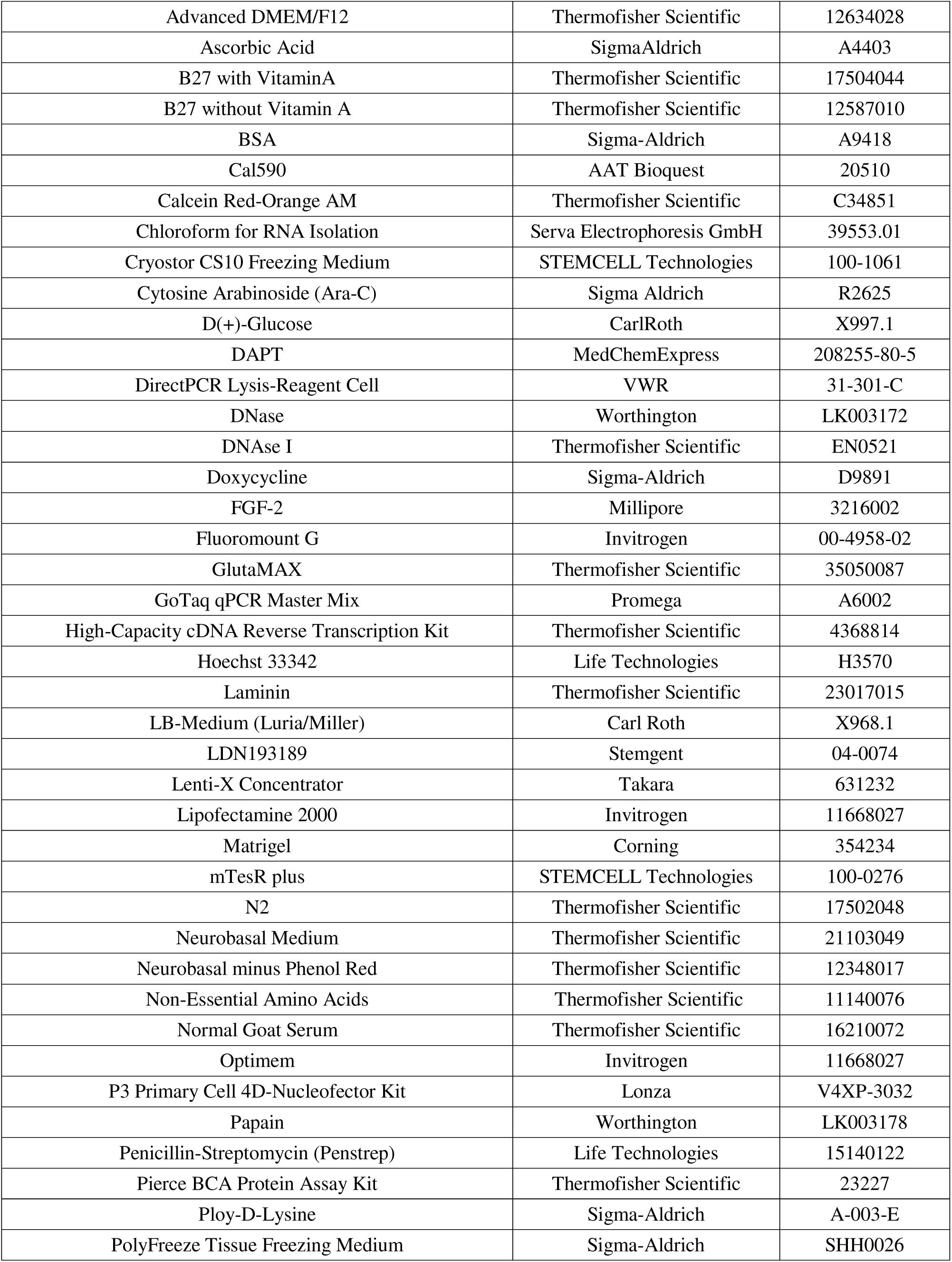
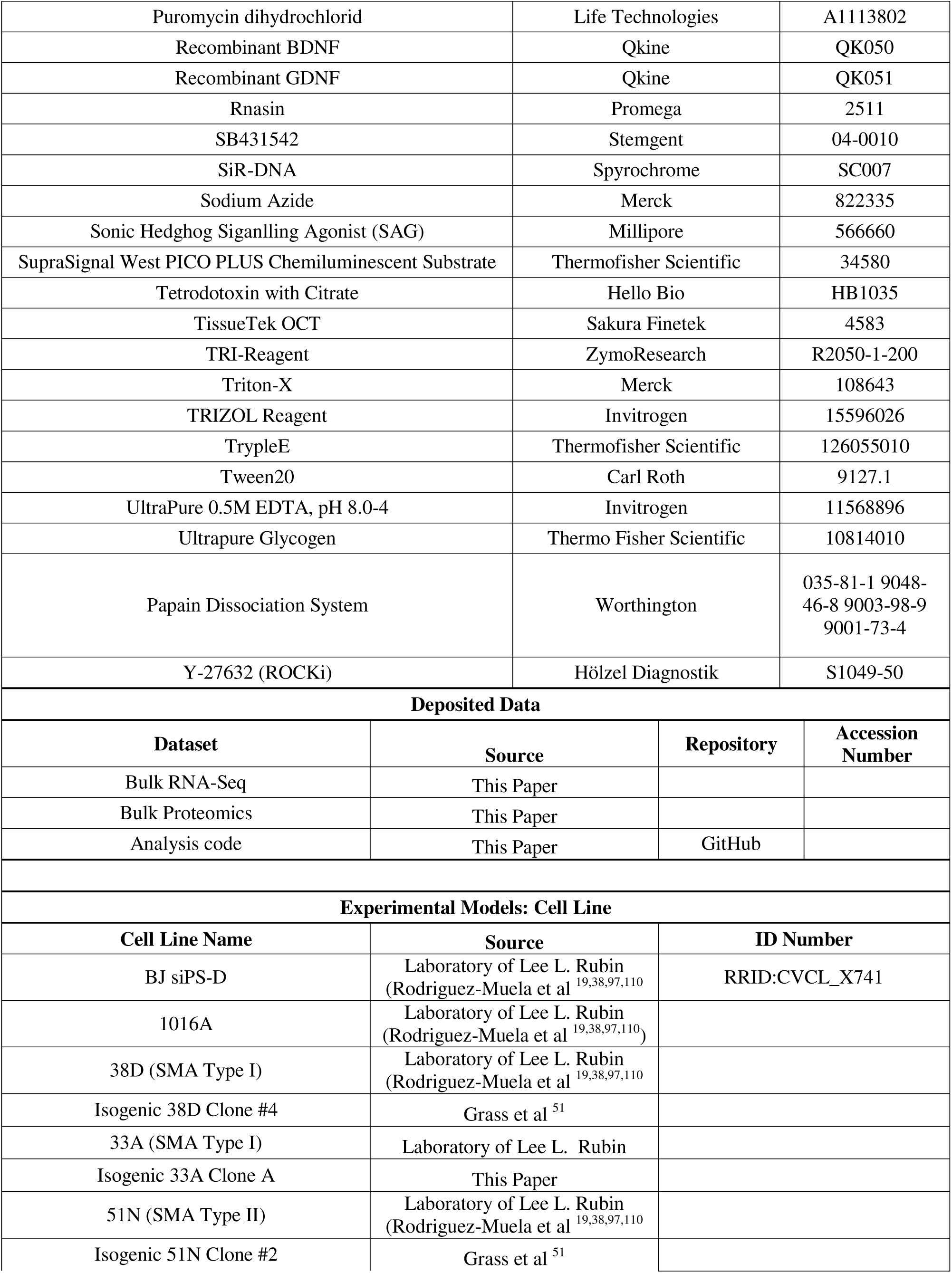
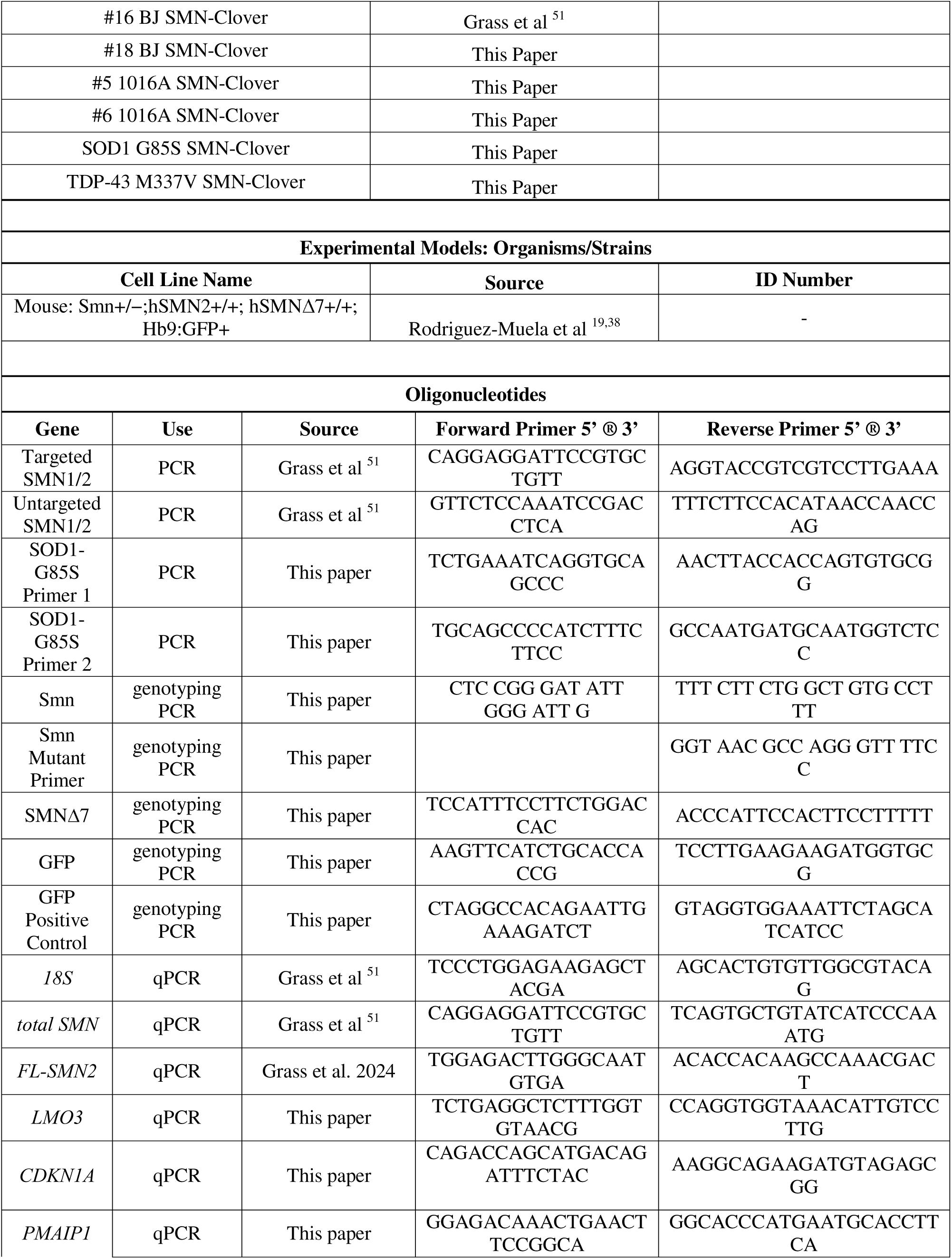
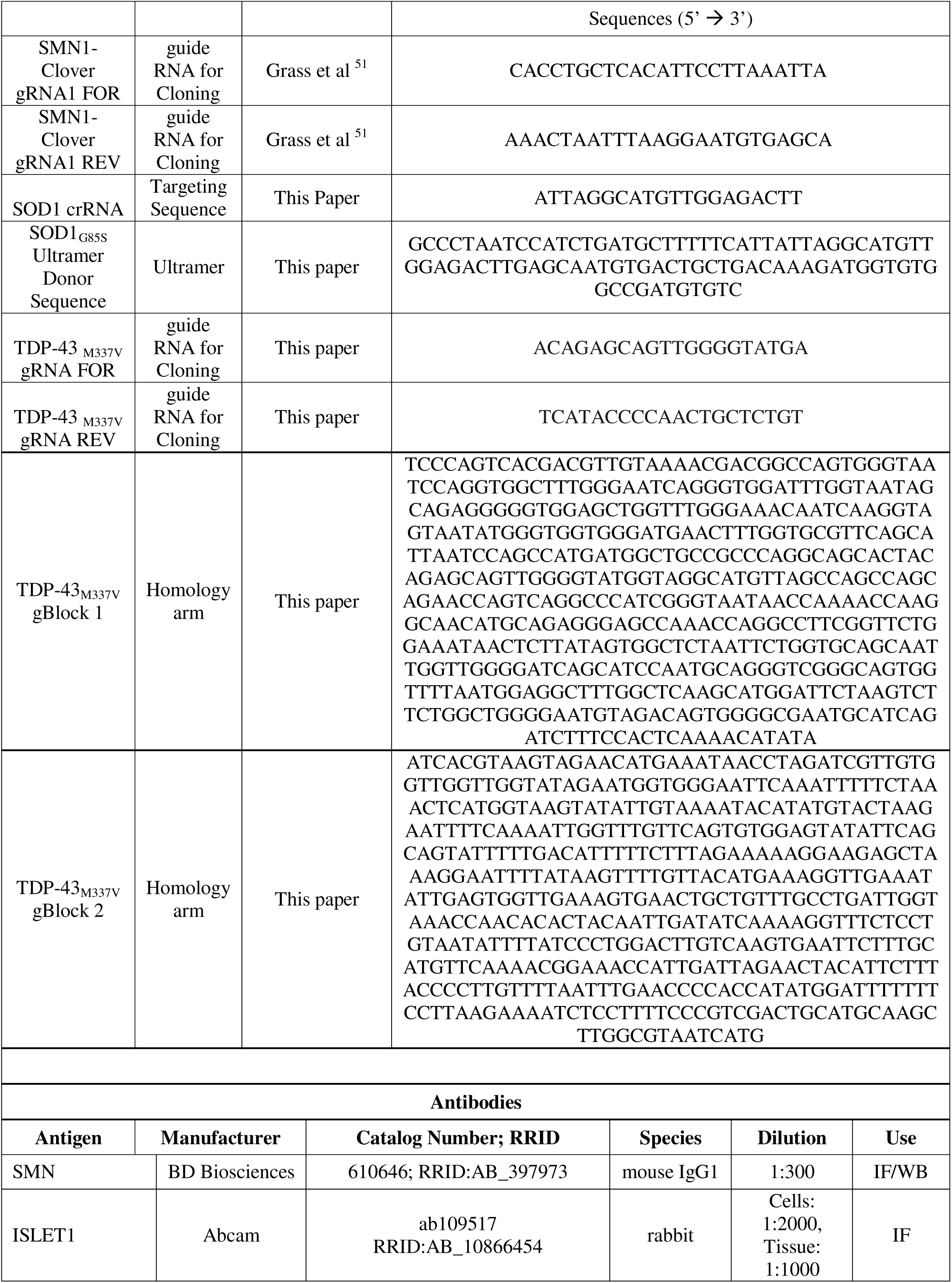
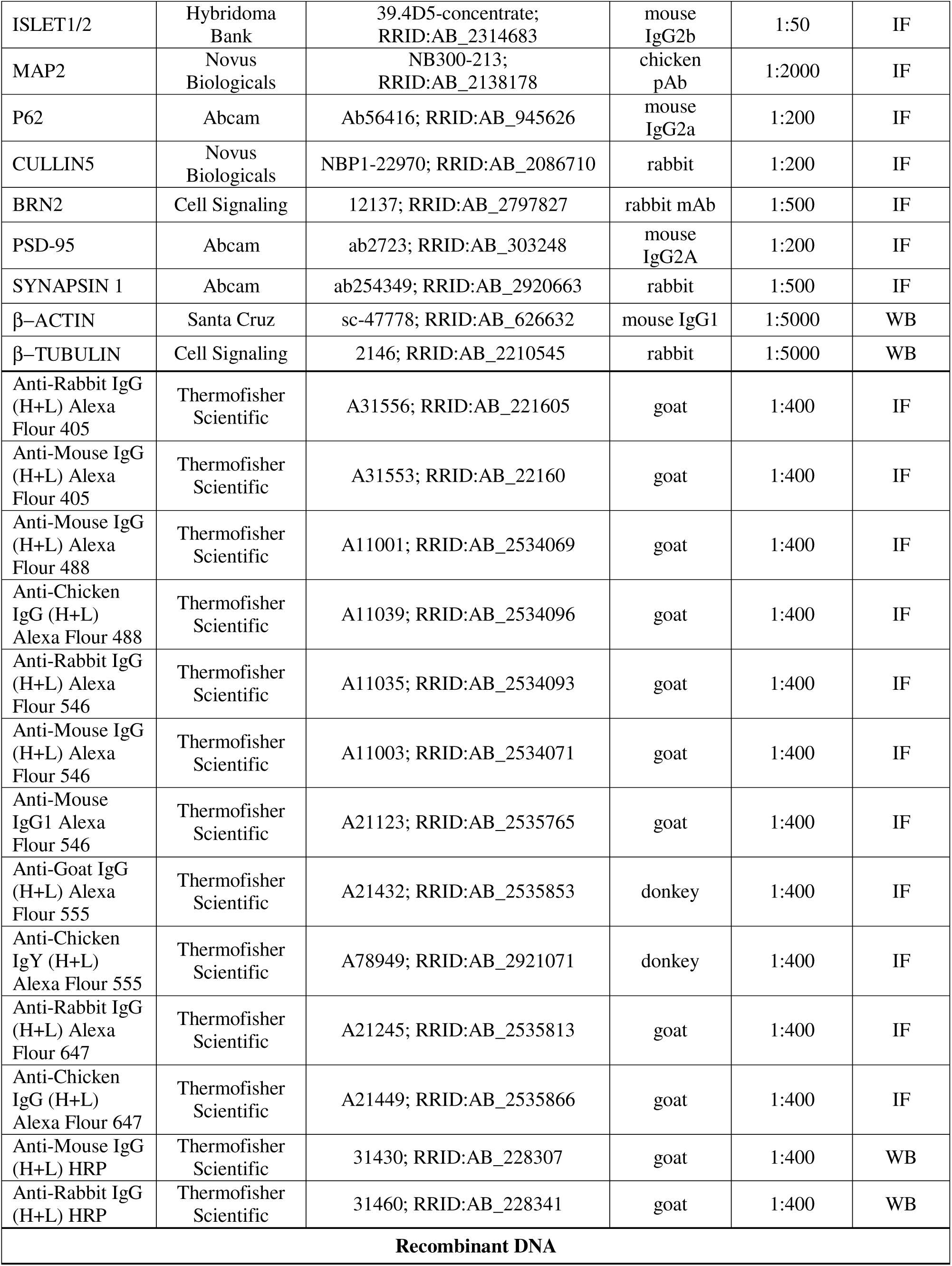
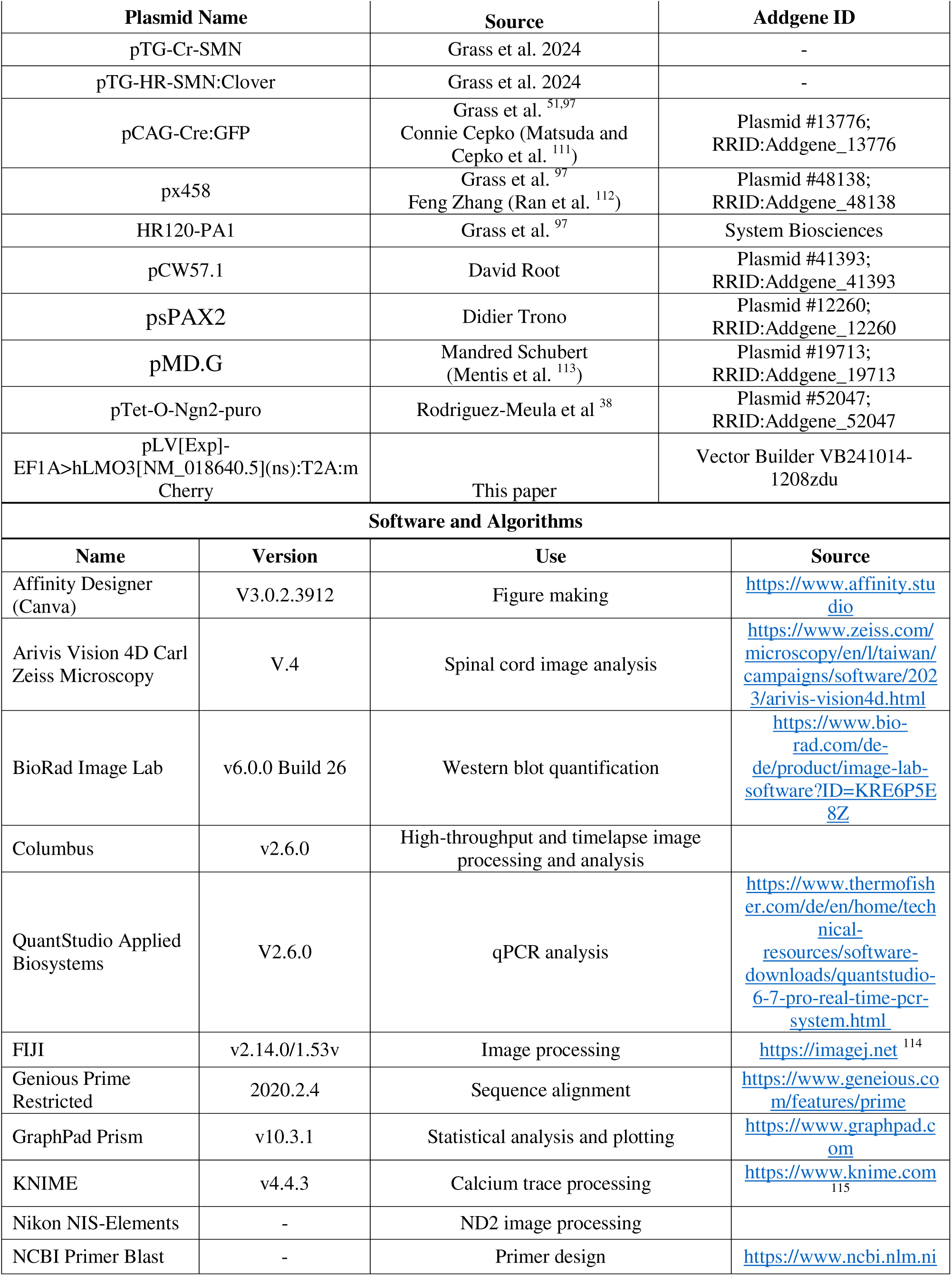
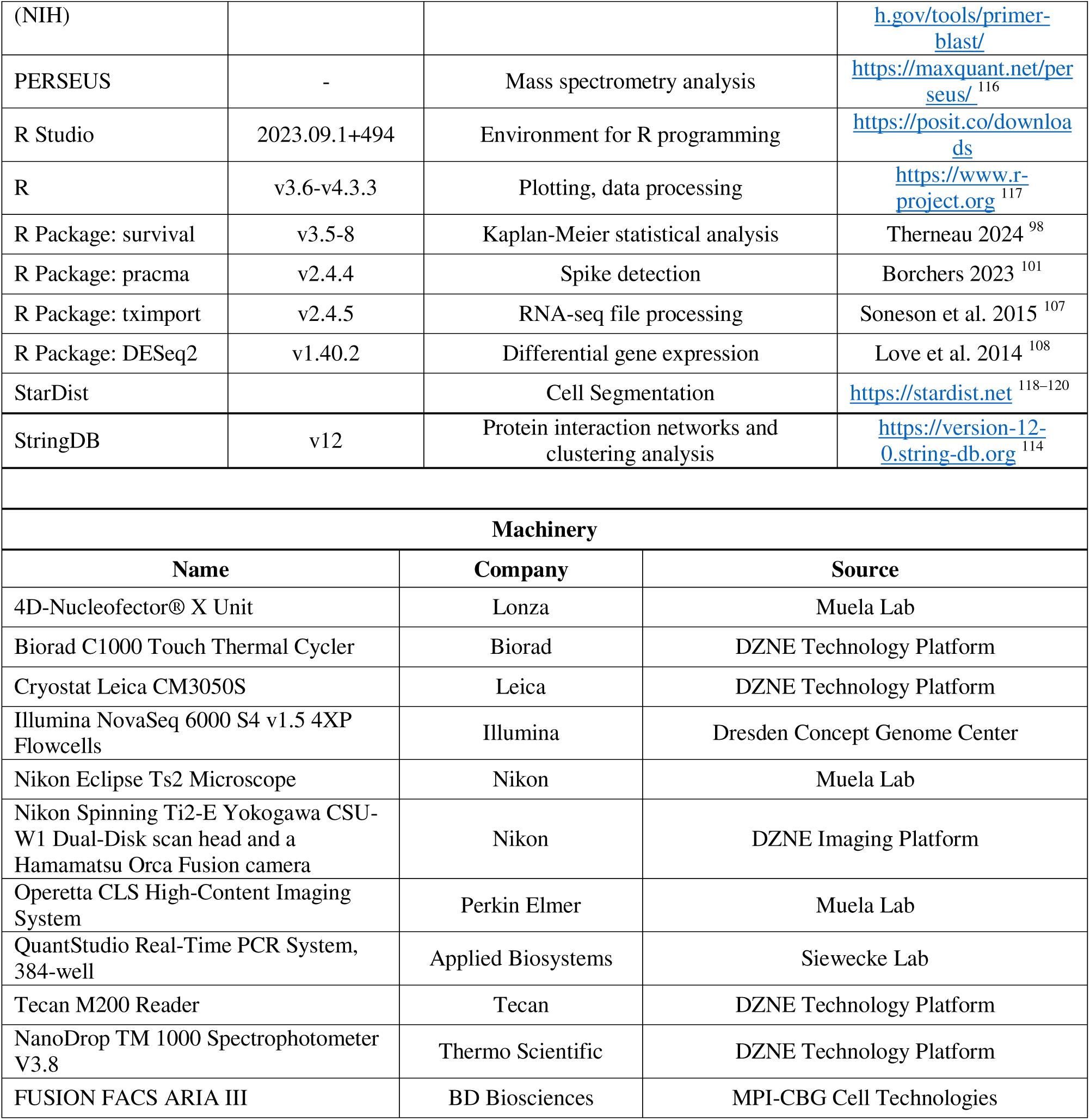

