## Supplementary figures and images for "Endogenous SMN Heterogeneity Defines Dynamic States of Motor Neuron Vulnerability and Resilience"

### Supplemental figure 1

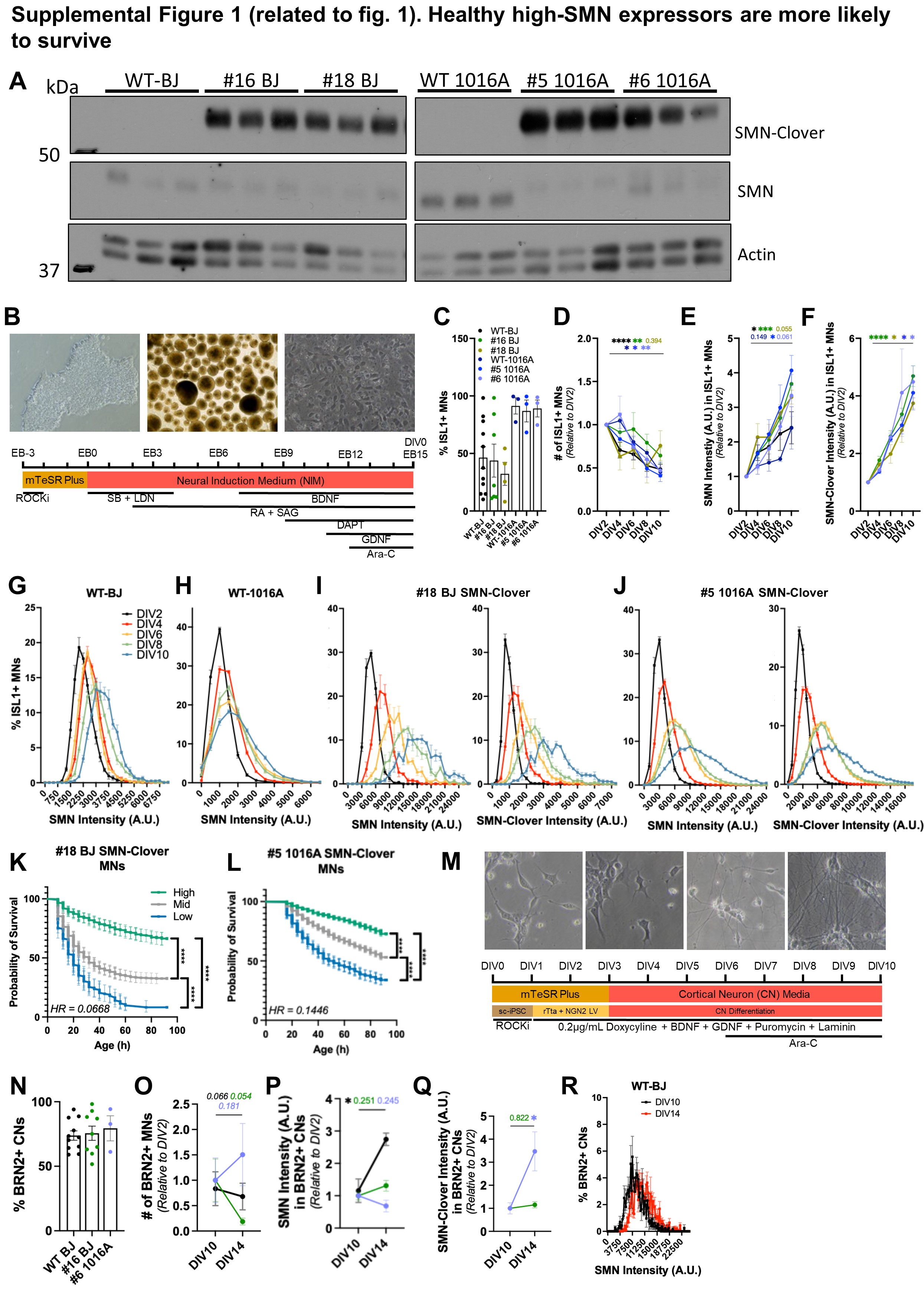

### Supplemental figure 2

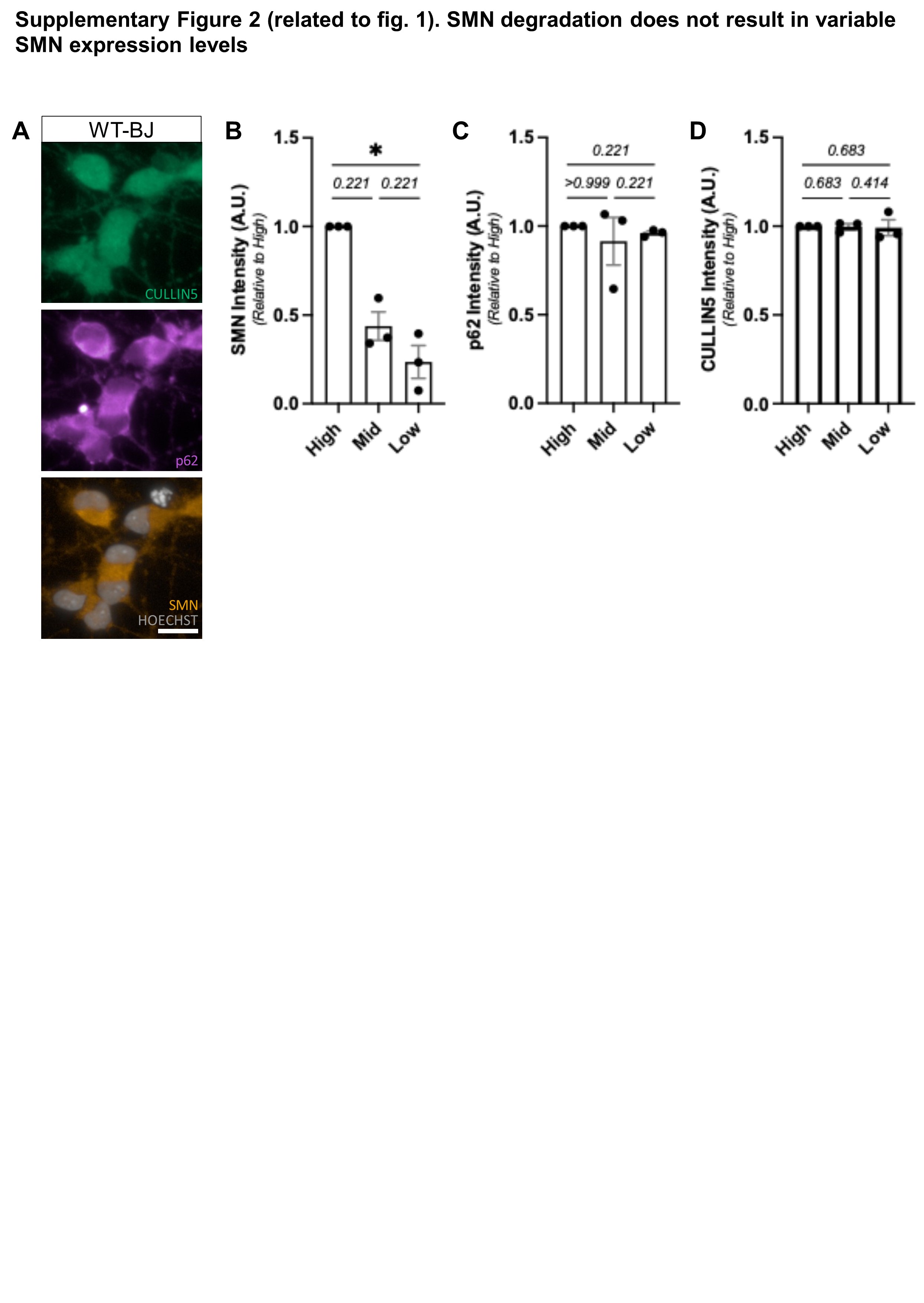

### Supplemental figure 3

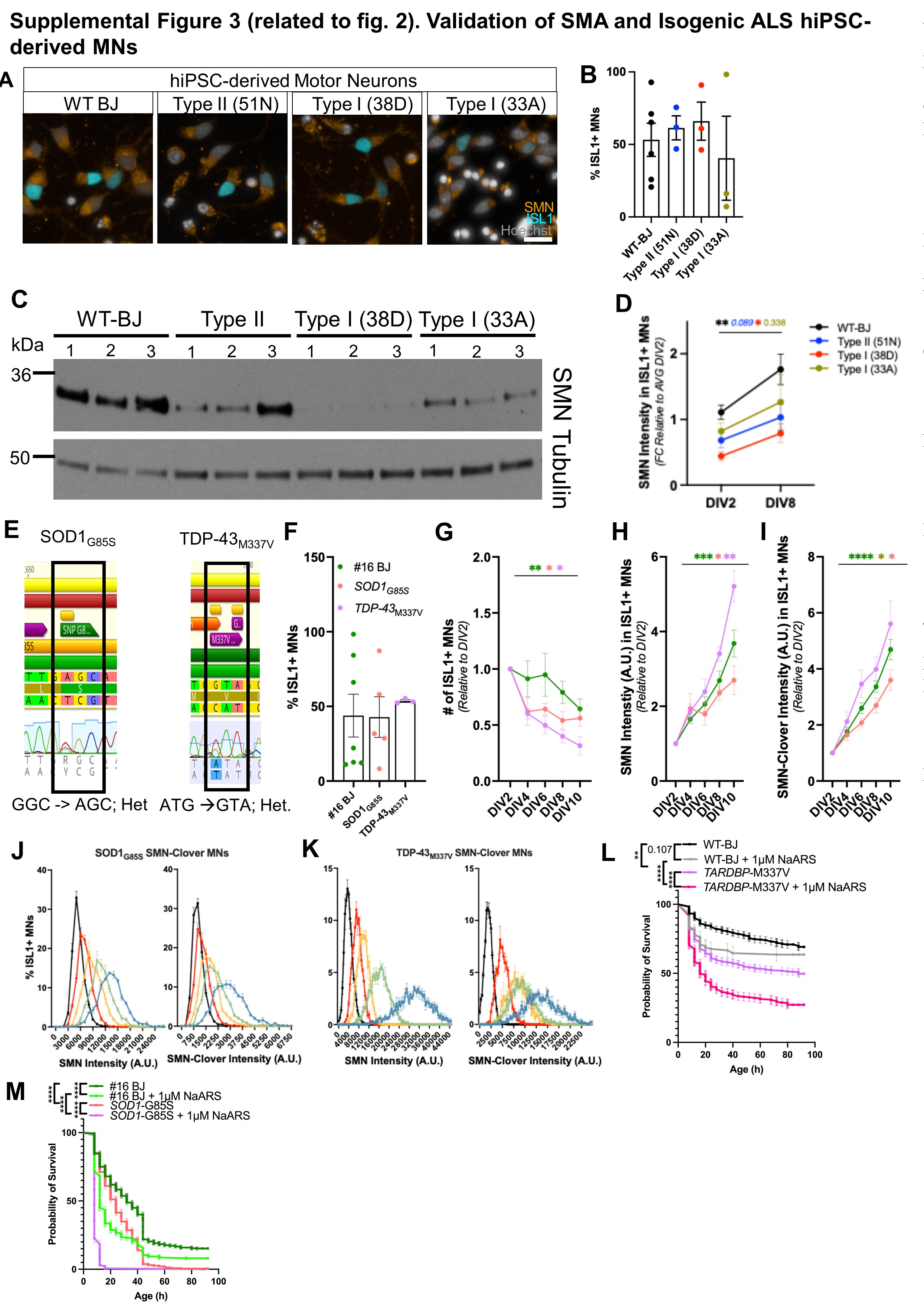

### Supplemental figure 4

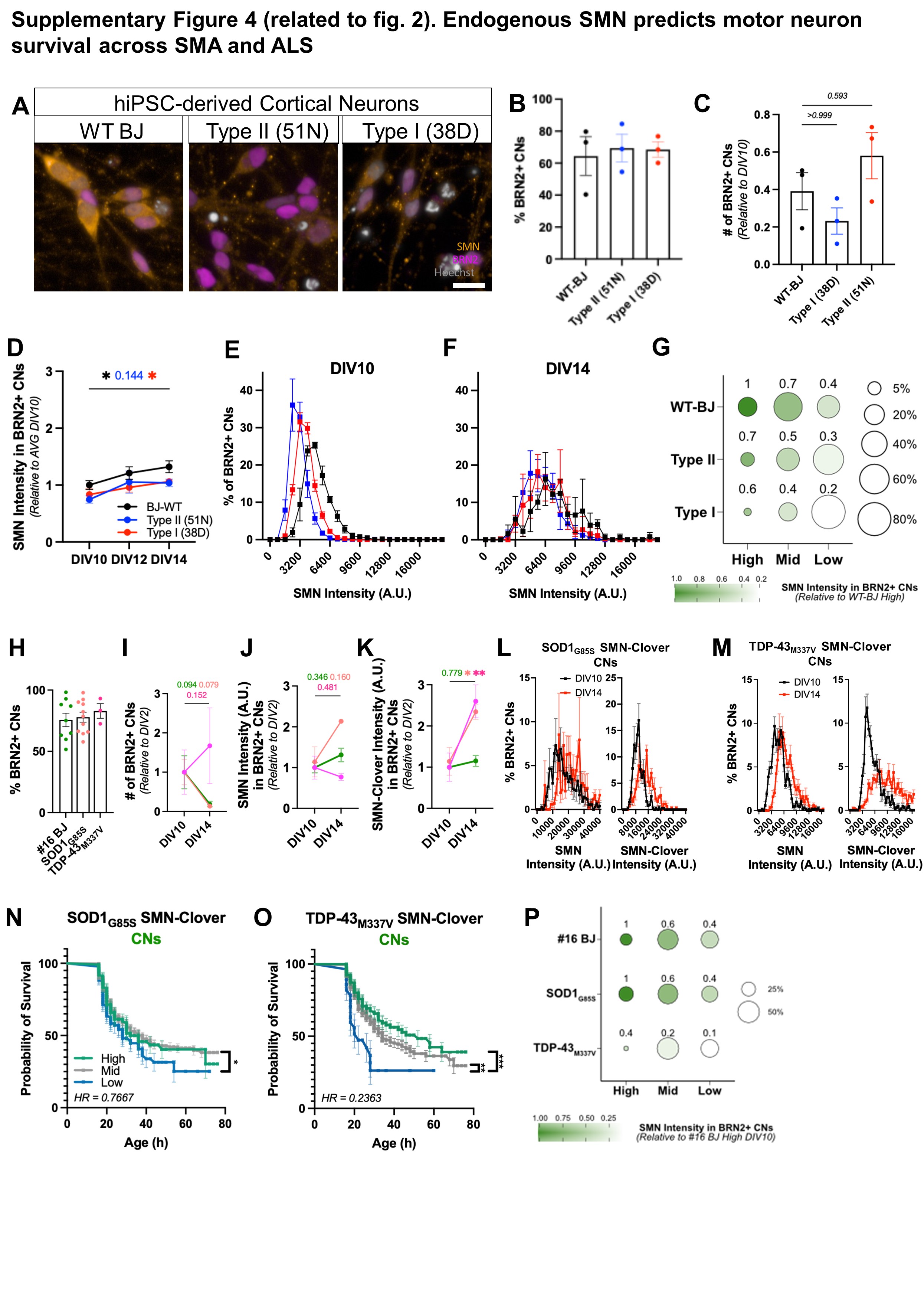

### Supplemental figure 5

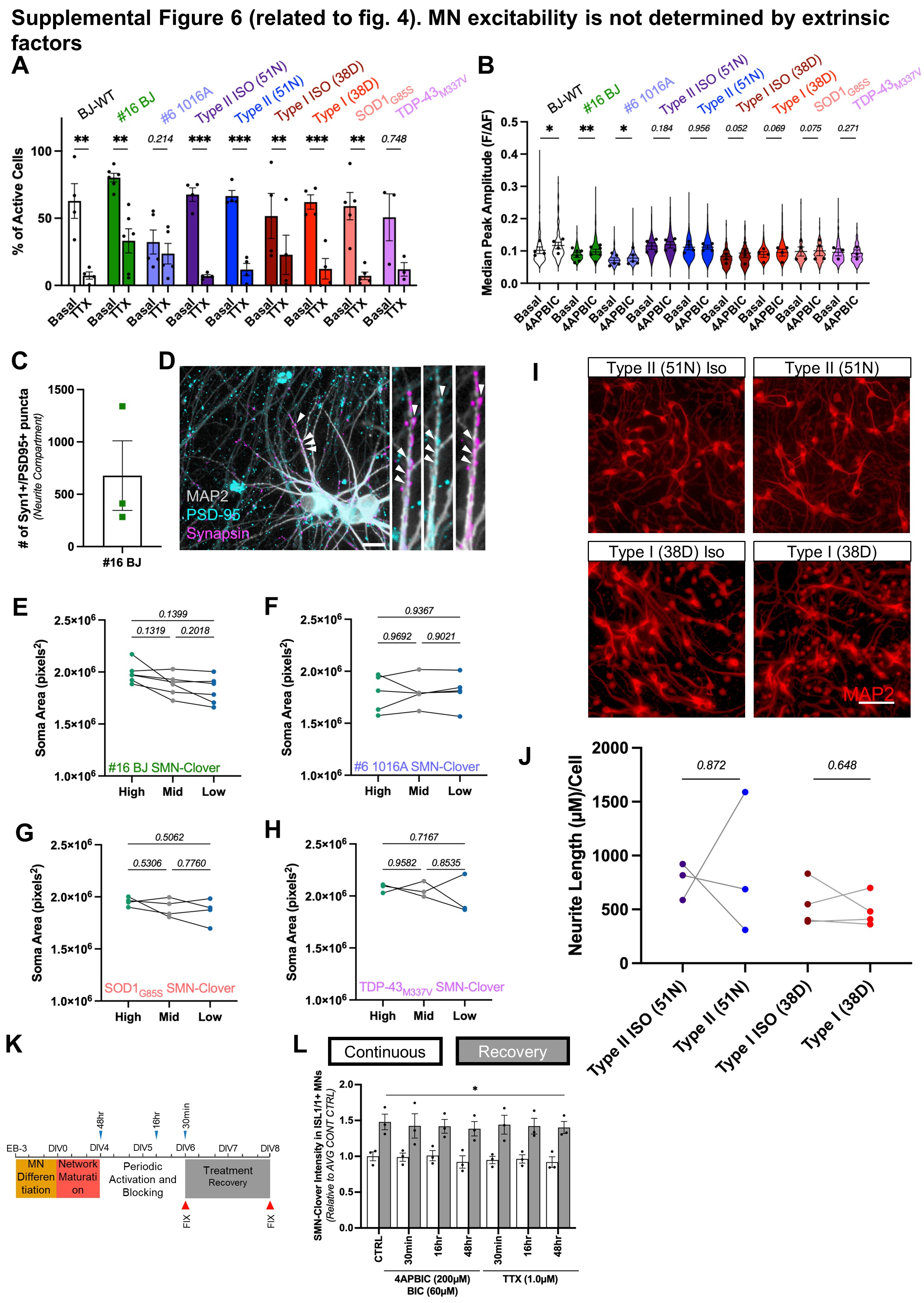

### Supplemental figure 7

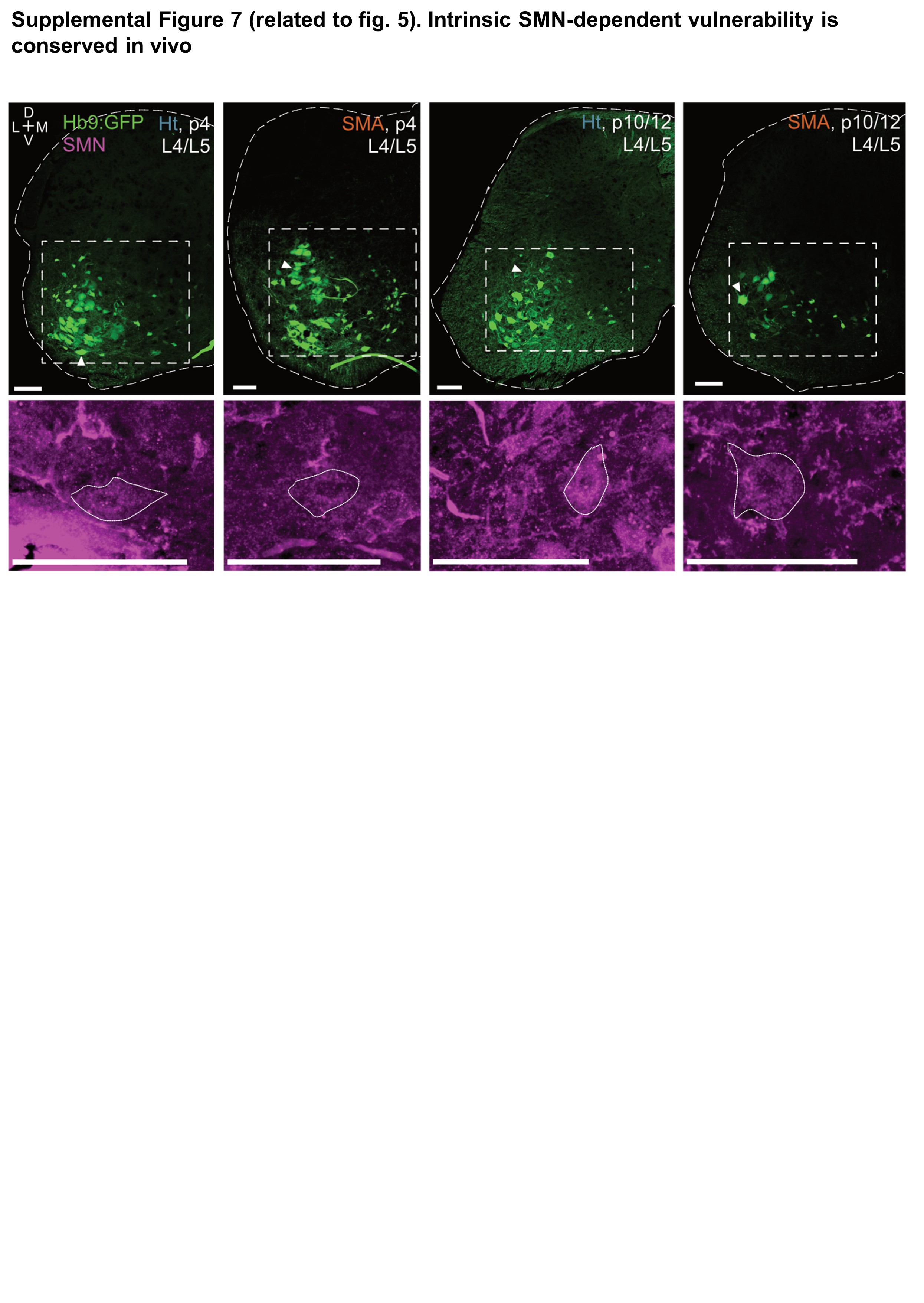

### Supplemental figure 8

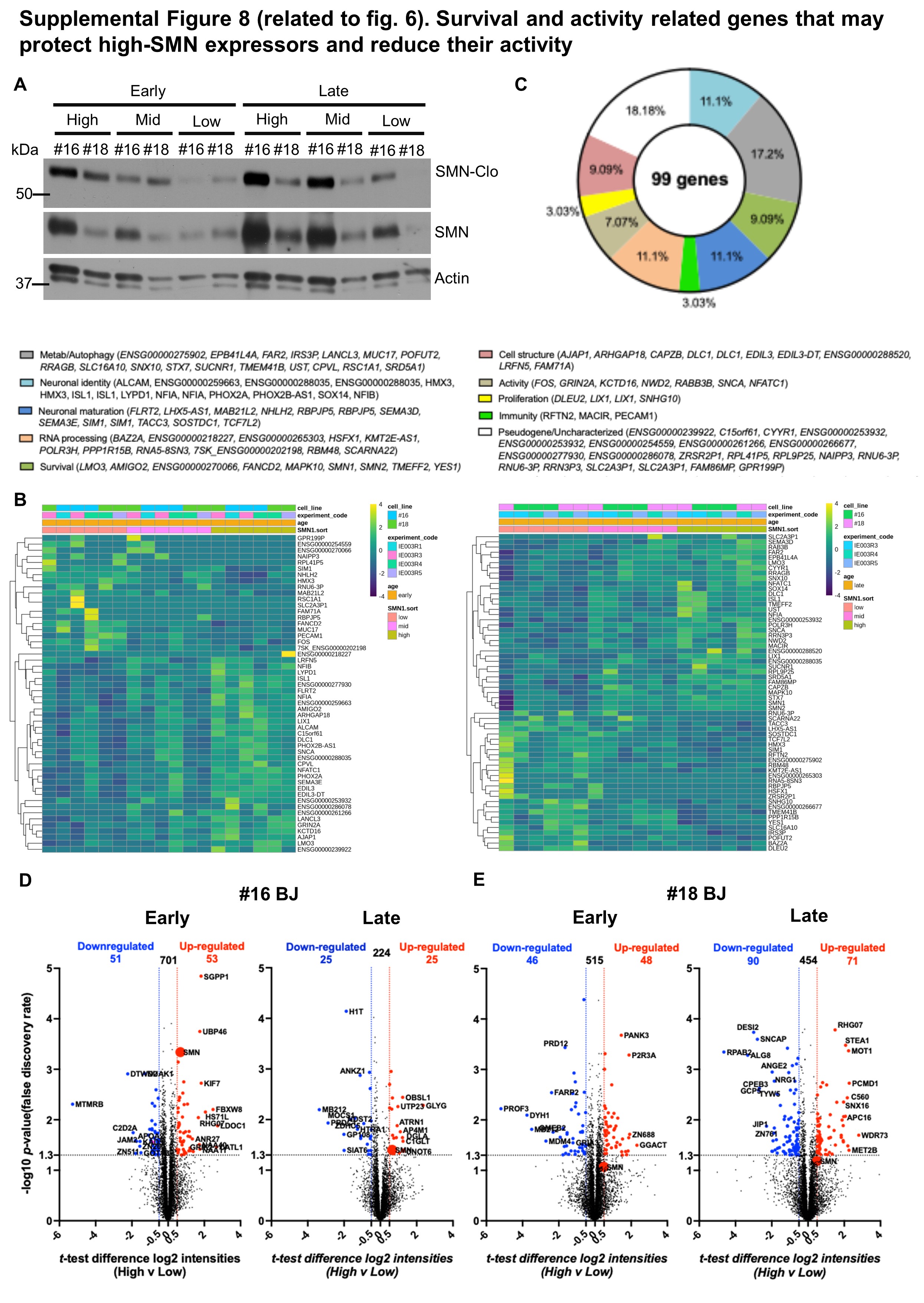

### Supplemental figure 9

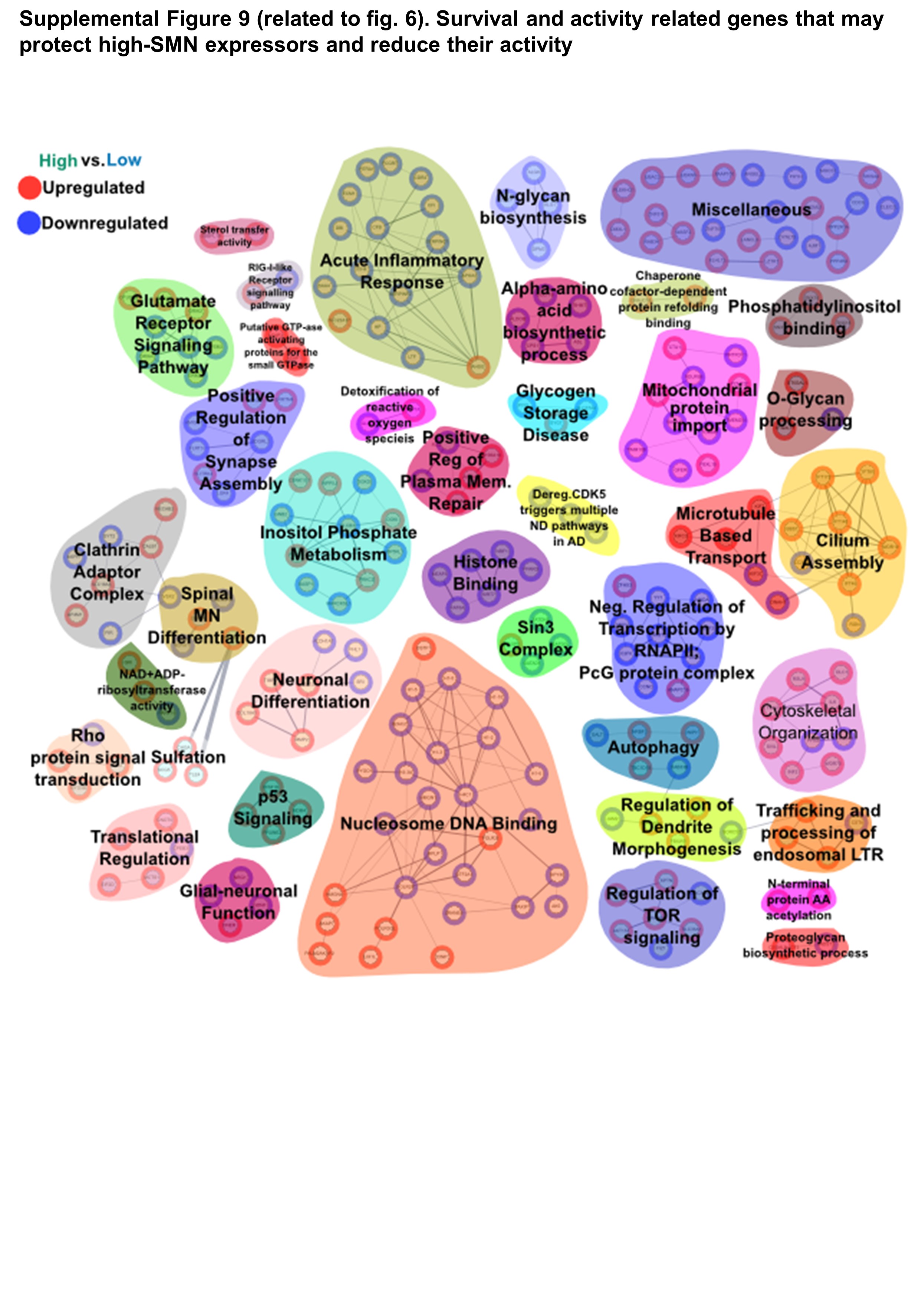

### Supplemental table 1

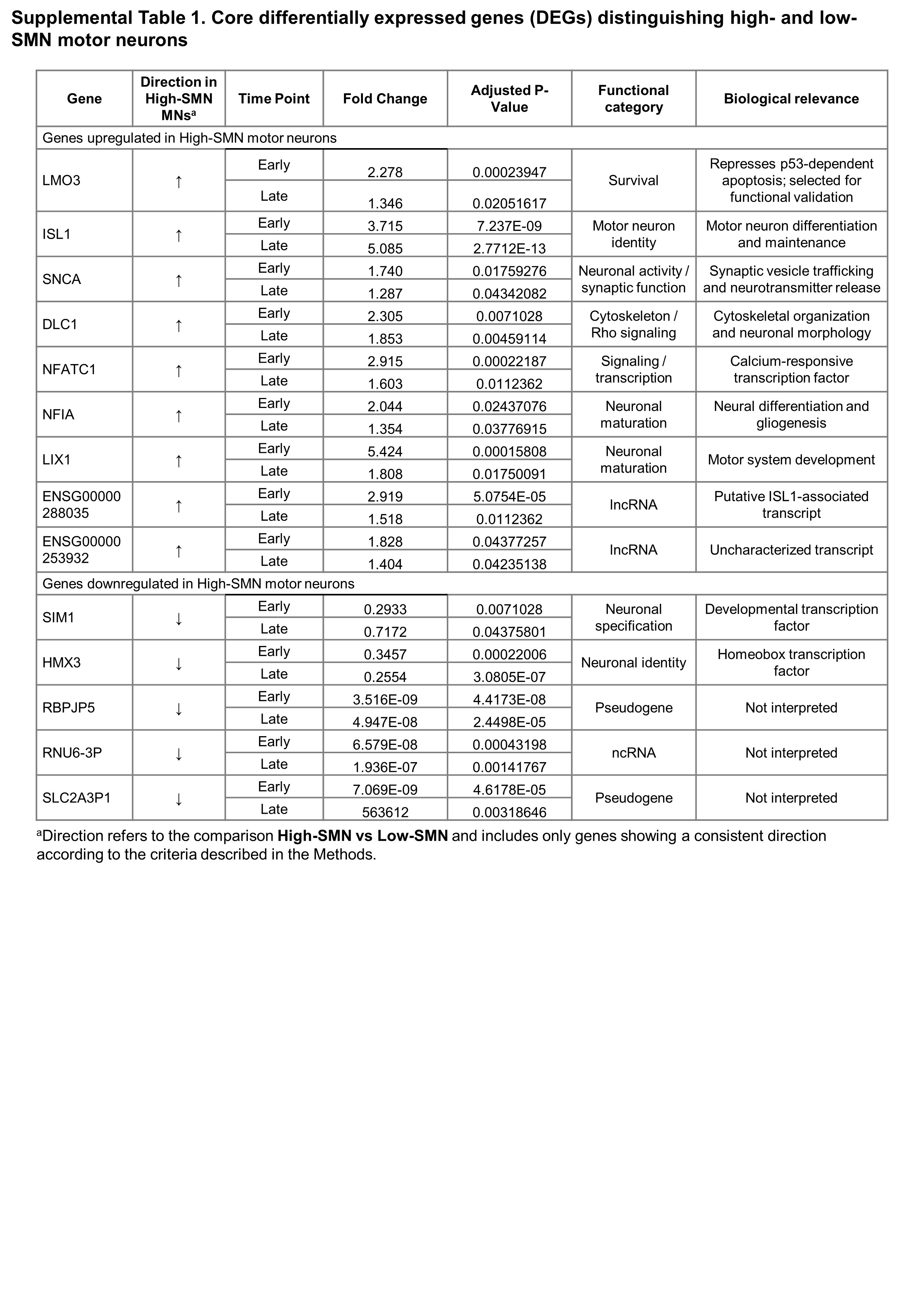

### Supplemental table 2

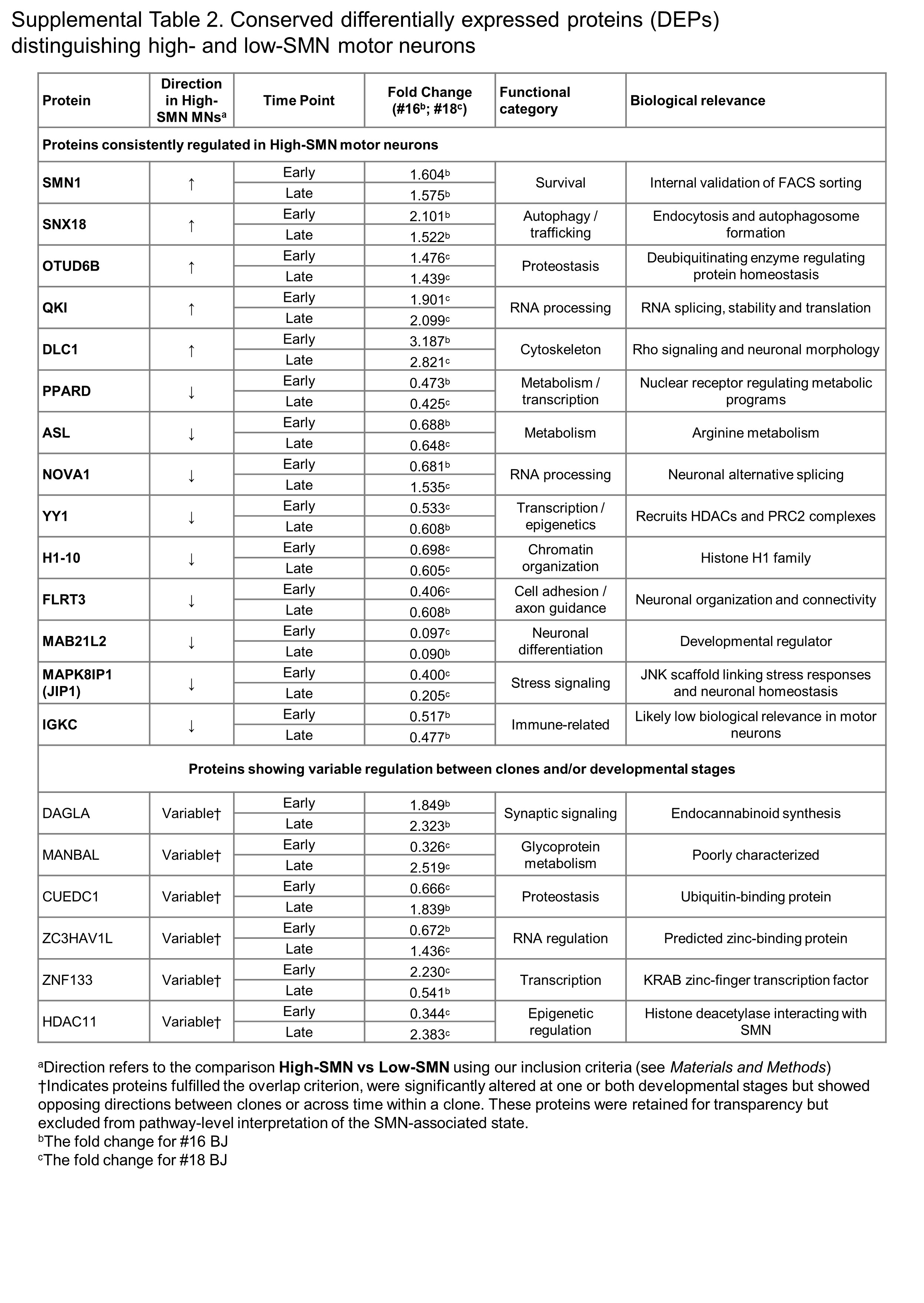
